# High-Quality Predicted Pathway Annotations Greatly Improve Pathway Enrichment Analysis of Metabolomics Datasets

**DOI:** 10.1101/2025.11.18.689105

**Authors:** Erik D. Huckvale, P. Travis Thompson, Robert M. Flight, Hunter N.B. Moseley

## Abstract

**Background/Objectives:** Metabolism-level interpretation of metabolomics datasets requires aggregation analyses across metabolites. One highly-used aggregation analysis is pathway enrichment analysis (PEA), which involves detecting pathways enriched with metabolites that are differential between experimental groups. Annotating metabolites with pathway associations is a prerequisite for PEA. While several knowledgebases define pathways and include metabolite-pathway annotations, these definitions are often partially or even grossly incomplete due to limitations in current metabolic knowledge and its curation, which greatly limits the effectiveness of PEA.

**Methods:** In this work, we used a novel multitask classification, graph convolutional-like neural network to generate high-quality metabolite-pathway annotations for pathways defined across KEGG, MetaCyc, and Reactome. We then included these predicted metabolite-pathway annotations when performing PEA on 990 datasets deposited in Metabolomics Workbench.

**Results:** We demonstrate an 8-fold increase in the median number of enriched pathways detected across these datasets compared to using only knowledgebase-derived annotations.

**Conclusions:** The significant increase in enriched pathways substantially improves the biological and biomedical interpretability of metabolomics datasets.

## 1. Introduction

Metabolism refers to the interconnected network of chemical reactions within cells and organisms that sustain life processes. Metabolism is also directly involved in many biological and disease processes, so it is biologically and biomedically meaningful to detect and understand changes in metabolism. Since metabolites are reactants and products of chemical reactions in life processes[1–3], their systematic detection and characterization, i.e. metabolomics, provides many details about metabolism. Changes in metabolite abundances and fluxes between experimental groups can provide biological and biomedical insights, e.g. testing the effects of a drug, examining differences between diseased versus healthy groups, etc [4,5].

However, it is very difficult to directly and systematically evaluate changes across hundreds to thousands of measured metabolites to generate biological and biomedical interpretations. To aid in systemic biological and biomedical interpretation of metabolomics datasets, data at the metabolite abundance and flux level can be aggregated to the pathway level, where a pathway can represent metabolic pathways, biological pathways, or any other collection of metabolites associated with some biological or biomedical concept. The metabolites that are part of such a collection are “associated with the pathway”. The most common pathway aggregation analysis of metabolomics datasets is pathway enrichment analysis (PEA), which is an annotation enrichment analysis (AEA) of pathway annotations, where each metabolite is annotated with the set of pathways it’s associated with [6],[7],[8],[9],[10],[11],[12],[13],[14],[15],[16],[17]. In PEA, evaluation metrics of metabolite abundance differences, like p-values, log-fold changes, etc., are analyzed to detect “enrichment” of metabolites associated with a given pathway. When the metabolites associated with a pathway collectively are enriched at a desired level of statistical significance (i.e., very low probability to be due to random chance), the pathway is defined as enriched.

In order to perform PEA, detected metabolites must have associated pathway annotations. Knowledgebases such as the Kyoto Encyclopedia of Genes and Genomes (KEGG) [18–20], MetaCyc [21], and Reactome [22] contain compound entries with pathway annotations, but only for some metabolites, and even then the metabolites with pathway annotations do not always have a complete set of annotations. So, a recurring problem with PEA is a lack of pathway annotations available or accessible from such knowledgebases [11],[23],[24]. A main reason for this incompleteness is that the knowledgebases are at best based on “known” metabolism, if knowledgebase curation is up-to-date with current peer-reviewed scientific literature. Both enzyme promiscuity and unknown (moonlighting) enzymatic functions represent significant portions of “unknown” metabolism that are still being discovered. This is evidenced by a significant fraction of compound entries in metabolic knowledgebases not being annotated to at least one pathway or enzymatic reaction. For example, over 60% of KEGG compound entries lack pathway annotations [25]. To further quantify these effects on incompleteness, we see that there are roughly 20,000 reactions in KEGG and MetaCyc combined, which is only 50% of the known enzymatic reactions in other knowledgebases like BRENDA, which has over 40,000 reactions [26]. Another reason is that different knowledgebases focus on different parts or aspects of metabolism [24]. Finally, many metabolites detected in metabolomics datasets are not easily corresponded to the correct knowledgebase entries without the database-specific compound identifier or some universal identifier like the IUPAC International Chemical Identifier (InChI) [27],[28],[29]. When performing PEA, missing pathway annotations greatly limit the number of enriched pathways detectable in metabolomics datasets, therefore limiting the biological and biomedical interpretability and insight gained from metabolomics datasets.

Solutions to reducing missing pathway annotations include improving data curation to more comprehensively document the metabolite-pathway annotations that are known, as well as to perform the wet lab research necessary to discover the metabolite-pathway associations that are still unknown. Currently, both tasks are prohibitively time consuming and financially costly. In the future, data curation of known metabolism may be automated using deep learning models, as is being demonstrated in the generation of gene annotations [30],[31]. However, there is no obvious approach for speeding up and reducing the cost of wet lab discovery of “unknown” metabolism. To more directly address the issue of missing pathway annotations, our lab is actively developing machine learning (ML) models for predicting the pathway associations of metabolites using chemical structures of metabolites as input [32],[25],[33],[34],[35],[36],[37]. Our current best approach, which is also the best demonstrated so far in the field [38], uses a graph convolutional-like multilayer perceptron (MLP) model that can predict metabolite-pathway annotations for a total of 22,265 pathway IDs defined across KEGG, MetaCyc, and Reactome where 8,517 of these are unique pathway definitions [38]. While a pathway ID is the label or name assigned to a pathway by its respective knowledgebase, a pathway definition is the set of metabolites associated with the pathway. Since pathways of different IDs can have equivalent pathway definitions, the model can predict more pathway IDs than unique pathway definitions. In this work, we focus on the unique pathway definitions to avoid redundancy and overly optimistic performance results.

The single binary classification model uses paired metabolite and pathway atom coloring features [39–41] representing all 1-bond, 2-bond, and 3-bond subgraphs in the metabolite and across metabolites annotated to a pathway. This enumeration of all chemical subgraphs provides similar intermediate results to a graph convolutional neural network [42],[43], with the resulting features providing a representation of the chemical graph that can be used for global graph classification tasks. The pairing of metabolite-specific and pathway-specific chemical subgraph features creates a multitask classification model [44],[45],[46] and more specifically a multitask extreme classification model [47],[48],[36], enabling transfer learning [49],[50],[51] between specific pathway prediction tasks [34],[35],[37],[38]. Huckvale and Moseley demonstrated a model performance with a mean Matthew’s Correlation Coefficient (MCC)[52] of 0.9036 ± 0.0033(sd), computed by 100 train-test splits for a dataset with roughly 50,000,000 feature vectors[38]. Also, the model became significantly more generalizable when using InChI standardization[27–29] of the chemical structure representations prior to training, testing, and novel prediction[38]. In this work, we demonstrate how the resulting predictions increase the median number of enriched unique pathway definitions 8-fold across a large analysis that reuses 990 metabolomics datasets obtained from the Metabolomics Workbench (MW) repository[53]. We fully expect the increased number of enriched pathways, detectable as a result of the increased pathway annotations, will substantially improve the results of future metabolomics PEA and meaningfully improve the biological and biomedical interpretability and insight obtainable in the field of metabolomics.

## 2. Materials and Methods

### 2.1. Downloading, cleaning, and filtering metabolomics datasets

To analyze the improvement in PEA afforded by our ML model’s pathway annotation predictions, we used the mwtab Python package[54,55], previously developed and recently improved in our lab, to download 4,372 metabolomics datasets in mwTab format from the Metabolomics Workbench (MW) repository[53] on April, 26, 2024. We then used a prototype rcha_metab (repair, clean, harmonize, augment) Python package, being developed in our lab but not yet publicly available, to repair unparsable mwTab files, clean the resulting files, and harmonize field names and values, in order to make them usable in our meta-analyses. Due to the large number of formatting and harmonization issues in MW deposited datasets [56],[55], the vast majority of the downloaded datasets would not have been usable in the PEA meta-analyses presented here without the application of the prototype rcha_metab package.

Connecting metabolites in the MW datasets to compound identifiers (IDs) in a major metabolic or chemical knowledgebase like KEGG and PubChem[57],[58] is a crucial step to detecting enriched pathways. Since the chemical structure information is not provided with the MW data, the compound ID must be known to retrieve the corresponding mol-file[59] containing the chemical structure data needed by the ML model to predict pathway annotations. And since our analysis involved comparing the effects of the known annotations to the predicted annotations, we also needed the compound IDs for at least some of the metabolites to access the known pathway annotations from KEGG, MetaCyc, and Reactome. While the KEGG, MetaCyc, and Reactome (ChEBI) compound IDs enabled retrieving both a molfile and known pathway annotations, when only a PubChem ID was available, we could only use the predictions. Very few MW datasets have compound IDs for MetaCyc or ChEBI[60] (and thus to Reactome), but IDs from these knowledgebases can be retrieved from KEGG or PubChem IDs via cross referencing. This meant we at least needed either a KEGG ID or a PubChem ID and this requirement significantly reduced the number of usable datasets from the MW repository. Another requirement not met by all MW datasets was needing a sufficient number of compound IDs even if some were available. For a pathway to be reasonably detectable as enriched at an adjusted p-value ≤ 0.01, it must be associated with at least 15 metabolites. In some cases, the MW dataset did not contain at least 15 metabolites at all, regardless of whether their IDs were provided or not. Thus, if there were less than 15 compound IDs in the dataset, it is pragmatically impossible for any pathways to be detectable, and we filtered out the dataset. Other relatively minor requirements not met included the metabolite intensities being at an appropriate scale (i.e. the minimum intensity being greater than or equal to 0 and the maximum intensity being greater than or equal to 20), the file being readable by the mwtab Python package, and the data processing pipeline being capable of finishing without software errors. Figure 1 details the checks made to determine whether to keep or filter an MW dataset in the order that they were checked. Table S1 details the amount of MW datasets filtered per requirement not met, totaling to 3,382 datasets filtered, reducing the number of MW datasets usable for our analysis from 4,372 to 990.

**Figure 1.**
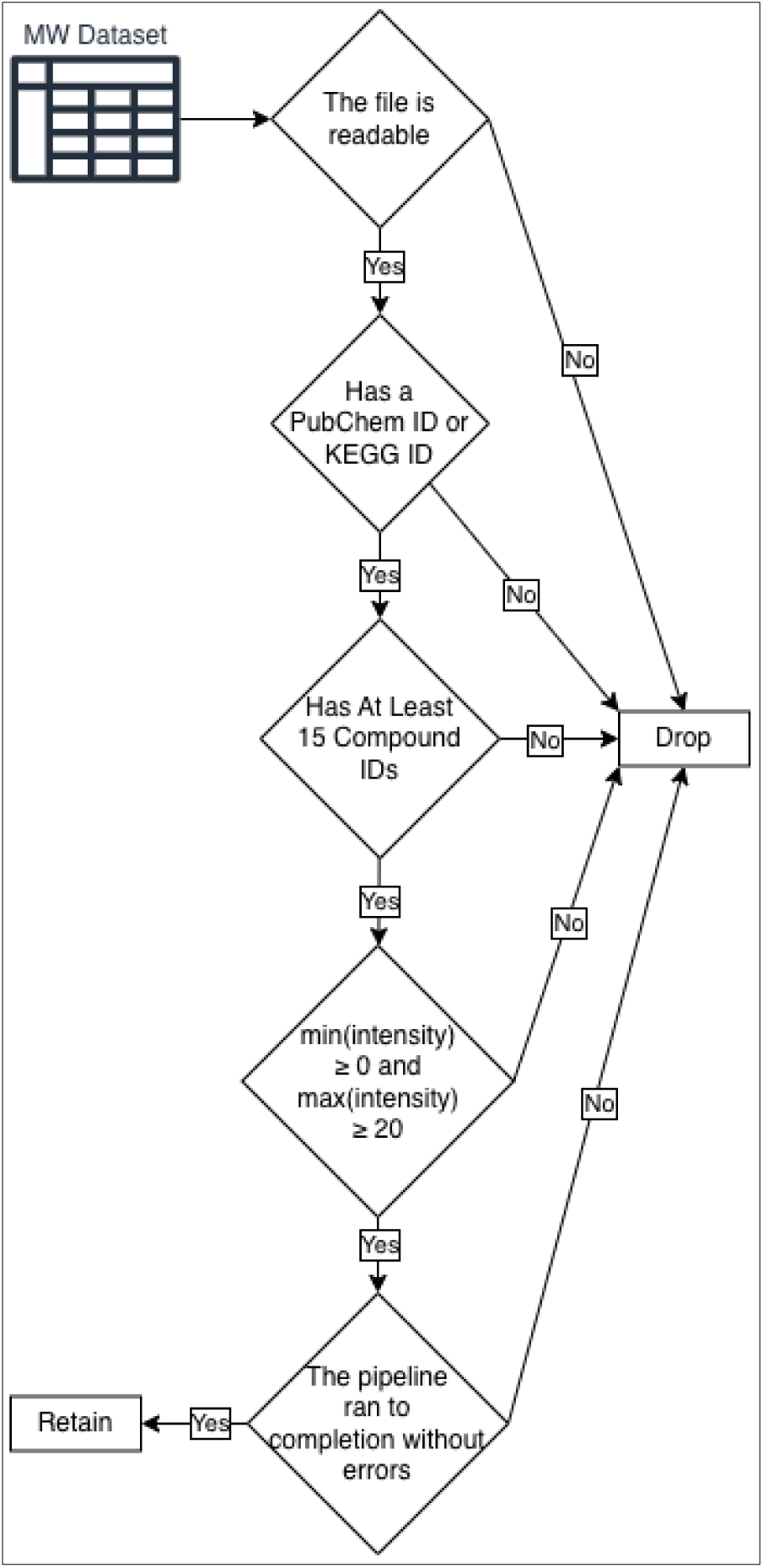
Checks made to determine whether to filter or keep an MW dataset for the analysis.

### 2.2. Resolving PubChem SIDs and CIDs

While PubChem does not contain pathway annotations[57,58], the annotations of pathways defined in the KEGG, MetaCyc, or Reactome knowledgebases can be obtained by cross-referencing the PubChem ID to a compound ID in one of these three knowledge-bases as well as by using the ML model on the provided PubChem molfiles[59] to predict pathway annotations. However, one specific issue that came up when using PubChem IDs was that PubChem has both CIDs (compound IDs) and SIDs (substance IDs)[57],[58]. This is a problem because both sets of IDs are just positive integers, they overlap, and most MW datasets do not indicate whether they are using CIDs or SIDs. So, if you are given an arbitrary PubChem ID, it could be either a CID or an SID. Since PubChem provides metabolite cross-references to KEGG, MetaCyc, and Reactome (ChEBI) via SIDs, and the file names of their molfiles contain CIDs, it must be known whether a PubChem ID in an MW file is an SID or a CID. This required us to write a Python script to compare the metadata of the metabolites in the MW datasets to both CID and SID records in PubChem to determine the correct type of PubChem ID. The script first compared the synonyms or names for the metabolite with the name given in the MW dataset. If a name strongly matched a CID record, we recorded the PubChem ID as a CID, otherwise it was recorded as an SID. If the name was neither matched to a CID record nor an SID record, then the chemical formula and molecular mass, if available in the metadata, were compared to make a match. Molecular mass and chemical formula matching were simple equality comparisons, but the name matching drew on code developed for the rcha_metab package. The name comparisons included Levenshtein distance matching (fuzzy matching using the fuzzywuzzy Python package[61]), and various other matching techniques based on specific naming conventions in different sub classifications of metabolites, such as lipids, isotopologues, and isomers. If a PubChem ID neither matched to an SID nor CID due to limitations or inconsistencies in the metadata (e.g. the ID corresponded to a CID while the synonyms in fact matched an SID), then the PubChem ID could not be used.

### 2.3. Chaining compound ID cross-references

We used the web-based cross-referencing functionality of PubChem to obtain compound IDs for KEGG, MetaCyc, and ChEBI/Reactome, converting PubChem CIDs to Pub-Chem SIDs as needed. PubChem provides cross references via its REST API mapping SIDs to external IDs in KEGG, MetaCyc, and ChEBI. If a metabolite in an MW dataset has an SID available, we can use the SID to map directly to cross references. However, if a CID is available, we first must map to its corresponding SIDs. Iterating through a list of SIDs corresponding to a given CID, we selected the cross reference of the first SID in the list with a cross reference available, if any. We did this for KEGG, MetaCyc, and ChEBI cross references for each metabolite with either an SID or CID available. Likewise, we used the web-based cross-referencing functionality available in KEGG, MetaCyc, and ChEBI to map compound IDs back to PubChem and each other. Finally, we implemented a Python script to perform chained cross-referencing i.e. cross-referencing back-and-forth between all four knowledgebases and seeing if the number of available metabolite IDs increased. The Python script repeated the back-and-forth cross-referencing, further augmenting the number of PubChem, KEGG, MetaCyc, and ChEBI IDs in the MW dataset until there was no further increase in compound IDs.

### 2.4. Accessing known metabolite-pathway annotations

We used the KEGG IDs directly available in an MW dataset to access known metabolite-pathway annotations in KEGG. We refer to these known metabolite-pathway annotations as the “ground truth”. We then used the KEGG, MetaCyc, and ChEBI compound IDs generated from the chained cross-referencing or already present in the MW dataset to access known metabolite-pathway annotations in KEGG, MetaCyc, and Reactome. We refer to these known metabolite-pathway annotations as the “ground truth with cross-referencing”. As of November 19^th^ 2024, there are 23,110 pathway IDs available across these three knowledgebases. However, many of these pathway IDs have redundant pathway definitions i.e. the set of metabolites associated with the pathway. While there are 23,110 pathway IDs, 8574 of these pathways have unique pathway definitions (Table 1).

**Table 1.** Comparing the number of pathways available in different contexts.

| #Known Pathway IDs | #Predictable Pathway IDs | #Known Unique Pathway Definitions | #Predictable Unique Pathway Definitions |
| --- | --- | --- | --- |
| 23,110 | 22,265 | 8574 | 8517 |

#### Predicting metabolite-pathway annotations

We used the KEGG compound IDs available in an MW dataset to predict pathway annotations for KEGG, MetaCyc, and Reactome using the molfiles provided by KEGG and standardized using InChI canonicalization. Next, we used the ML model to predict annotations to the pathways defined in KEGG, MetaCyc, and Reactome using the InChI standardized molfiles from the PubChem CID records for all PubChem CIDs available in the MW datasets, converting PubChem SIDs to PubChem CIDs as needed. PubChem molfiles are only available by their corresponding CID. If a CID is available for a metabolite in an MW dataset, we obtained the molfile directly. If an SID was available, we mapped the SID to its corresponding CID. We refer to these metabolite-pathway annotations obtained by predicting on chemical structure information from the molfiles as the “predicted annotations”. Next, we used the KEGG, MetaCyc, ChEBI, and PubChem compound IDs generated from the chained cross-referencing or already present in the MW dataset to access molfiles in the respective knowledgebase, which we standardized using InChI canonicalization, and then used to predict metabolite-pathway annotations for KEGG, MetaCyc, and Reactome defined pathways. We refer to these predicted metabolite-pathway annotations as the “predicted annotations with cross-referencing”. Figure 2 details the process of obtaining pathway annotations, including looking up known pathway annotations and predicting additional pathway annotations for both the original and cross-referenced datasets. While there were molfiles used by the ML model to predict pathway annotations for the respective metabolites, some of the molfiles used for constructing the dataset to train the model could not be successfully standardized. Because of this, some of the pathways in the dataset lost associated metabolites in their definitions. And some of these pathways lost all associated metabolites, resulting in those pathways being removed from the dataset. The ML model can only predict pathways available in the dataset it was trained on. Therefore, while 23,110 pathway IDs and 8574 pathway definitions are known and available for the ground truth, 22,265 pathway IDs and 8517 unique pathway definitions remained predictable by the ML model (Table 1).

**Figure 2.**
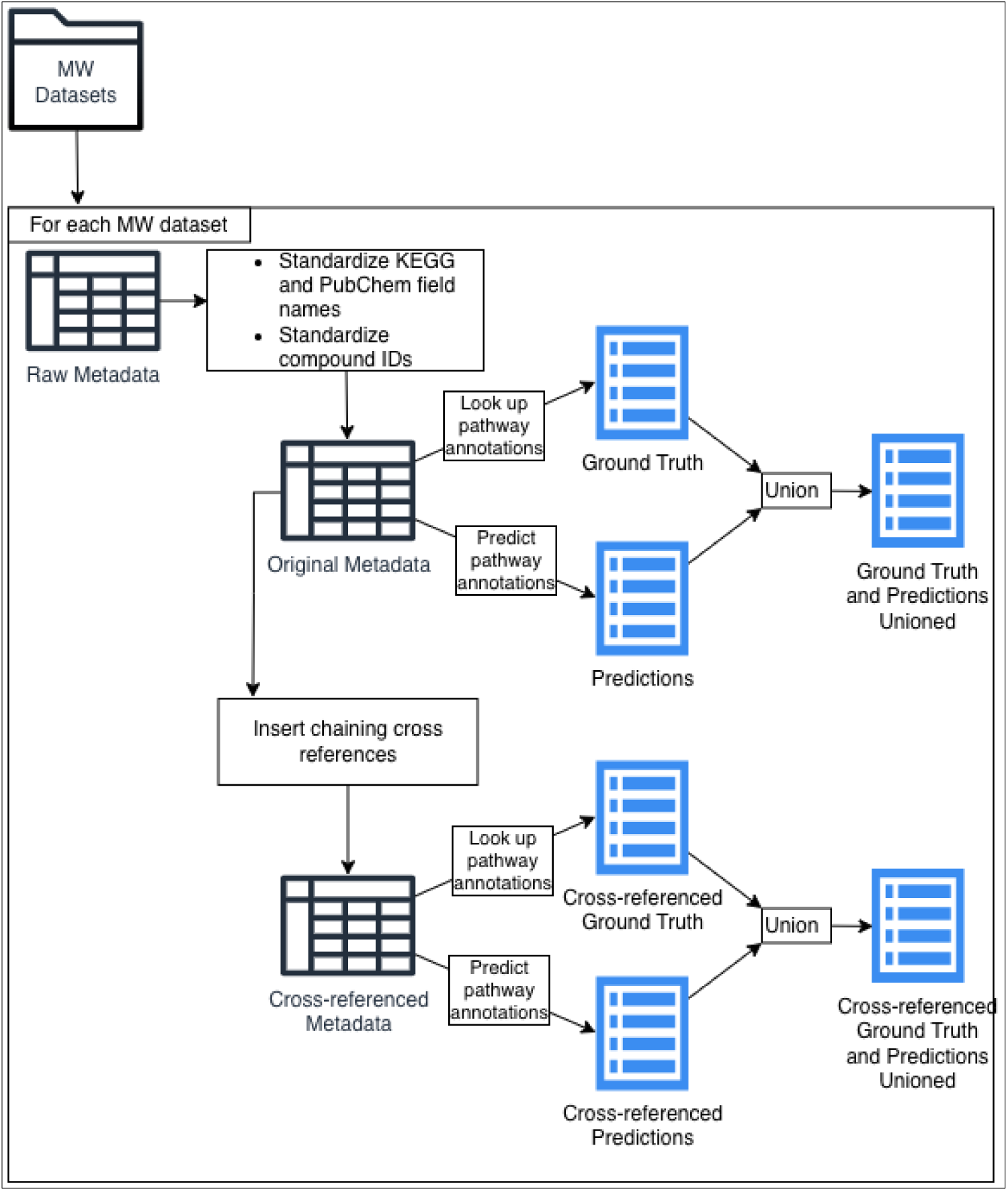
Using the MW datasets’ metadata to determine compound IDs and map them to pathway annotations.

### 2.5. Pathway enrichment analysis

Since the increase in enriched pathways could be attributed to the increase in available cross references, to prove the utility of the ML model beyond our chained cross-referencing algorithm, we computed enriched pathways both on the cross-referenced version of the MW datasets and the original dataset. We also compared the impact on the number of enriched pathways using the ground truth alone, the predictions alone, and the ground truth unioned with the predictions.

For all combinations of the original and cross-referenced MW datasets with ground truth annotations, predicted annotations, and ground truth unioned with predictions, we used the categoryCompare2 R package[62,63] developed in our lab to compute the number of enriched pathways for all 990 usable MW datasets. Specifically, we performed PEA using the Gene Set Enrichment Analysis (GSEA)[64] algorithm implemented in fgsea [65] and wrapped by categoryCompare2. GSEA requires the features (metabolites in our case) to be ranked. We performed Principal Component Analysis (PCA)[66] on the log-transformed metabolite intensity values in the MW datasets. To prevent values of negative infinity resulting from a log of 0, we replaced 0s with the minimum non-zero value divided by 10 prior to the log transform. Next, we used the resulting principal component (PC) loadings as the ranks for GSEA. Only the top PCs adding up to at least 75% of the total variance in the given dataset were used. With the categoryCompare2 package, we performed GSEA on each list of PC loadings followed by Benjamini-Hochberg multiple testing correction [67]. Pathways with an adjusted p-value ≤ 0.01 were considered enriched. This approach obviates the need to determine the experimental design of a given metabolomics dataset, since differences between experimental groups (i.e., systematic variance) are expected to be represented in the top PCs. However, analytical variance will also be detected as differences in certain PCs. But this does not affect the overall purpose of evaluating effects of predicted metabolite-pathway annotations on pathway enrichment analysis. Furthermore, this approach enables the analysis of all 990 metabolomics datasets to be fully automated. Finally, we statistically compared the number of enriched pathways between the original and cross-referenced datasets as well as between the datasets using ground truth annotations, predictions, and ground truth unioned with predictions using Mann-Whitney U tests[68]. While the matched-pair case-control experimental design allows the use of a Wilcoxon signed-rank test, we conservatively chose the Mann-Whitney U test, since the dataset had ample sample size for high statistical power. Figure 3 details the process of detecting enriched pathways. Note that when counting the number of detected pathways for a given set of pathway annotations for a given MW dataset, the counts are summed across the PCs selected to explain 75% of the variance. The summation of enriched pathways across PCs is not double counting; because, each PC is an orthogonal component of correlated variance which is separately being evaluated for improvement in statistical sensitivity and power.

**Figure 3.**
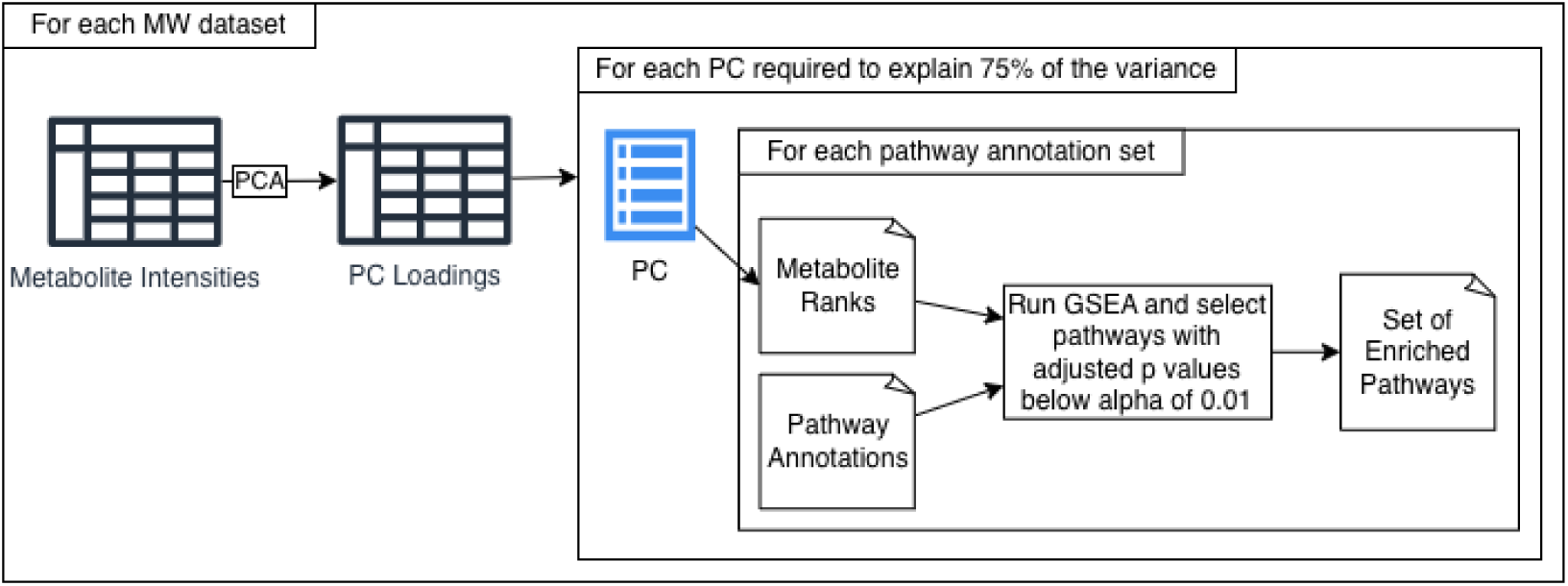
Detecting enriched pathways for each combination of PC and pathway annotation set.

### 2.6. Description of software packages and hardware used

While the categoryCompare2 (v0.200.5) package[62],[63], used to perform PEA via the GSEA algorithm, is written in the R programming language[69], all other code was written in the Python (v3.10.12) programming language[70], including the mwtab (v1.2.5) package[54,55]. The basic data processing was performed using the Pandas (v2.2.3)[71] and NumPy (1.26.4)[72] Python packages. PCA was performed using the Scikit-learn (v1.3.0) Python package[73]. The Mann-Whitney U tests were performed using the SciPy (v1.16.3)[74] Python package. We generated all of the data visualizations of results using the Tableau Public business intelligence software (v2024.3.3)[75] as well as the seaborn (0.12.2) Python package[76] built on top of the Matplotlib package (v3.7.2)[77] within Jupyter Notebooks (v1.1.1)[78].

The hardware for this work included computers with 64 gigabytes (GB) of random access memory (RAM) and central processing units (CPUs) ranging from 3.4 gigahertz (GHz) to 3.6 GHz, each with 6 hyperthreaded (HT) cores. The CPU chips included ‘Intel(R) Core(TM)i7-2600 CPU@3.40 GHz’, ‘Intel(R) Core(TM) i7-5930K CPU@3.50 GHz’, ‘Intel(R) Core(TM) i7-4930K CPU@3.40 GHz’, and ‘Intel(R) Core(TM) i7-6850K CPU@3.60 GHz’.

## 3. Results

### 3.1. Pre-GSEA

The following results describe the annotations prior to GSEA where the pathways counted are unique pathway definitions rather than pathway IDs. For a pathway to be detectable, the given MW dataset must have at least 15 metabolites with pathway annotations to the specific pathway. Figure 4a shows how the median number of detectable pathways across MW datasets changes depending on the set of metabolite-pathway annotations used. We see that using the ML model’s predictions greatly increases the number of pathways that are detectable. We also see that the median does not change whether using the predictions only or using the predictions unioned with ground truth. Figure 4b shows the same quantities as Figure 4a but rearranged to compare how adding metabolite ID cross references impacts the median number of detectable pathways. We see regardless of the annotations we use, the cross referenced datasets have a meaningfully higher number of detectable pathways. Figure 4c shows the median of the total number of metabolite-pathway annotations across MW datasets i.e. the sum of all metabolite associations across all pathways in a dateset. We likewise see that adding predictions greatly increases the median number of metabolite-pathway annotations and Figure 4d shows that adding cross references does the same; however, the increases from predictions is substantially larger. Figure S1 shows these results as violinplots to illustrate the distribution of the number of detectable pathways and total number of metabolite-pathway annotations across the datasets. While Figure 4 shows these results for unique pathway definitions, Figure S2 shows these results for individual pathway IDs. And while these are the results of KEGG, MetaCyc, and Reactome combined, Figure S3 shows this information for each knowledgebase separately (unique pathway definitions). The largest increases occur for MetaCyc– and Reactome-defined pathways, but these two knowledgebases represent over 90% of the uniquely defined pathways. Figure S4 shows these per knowledgebase results for individual pathway IDs.

**Figure 4.**
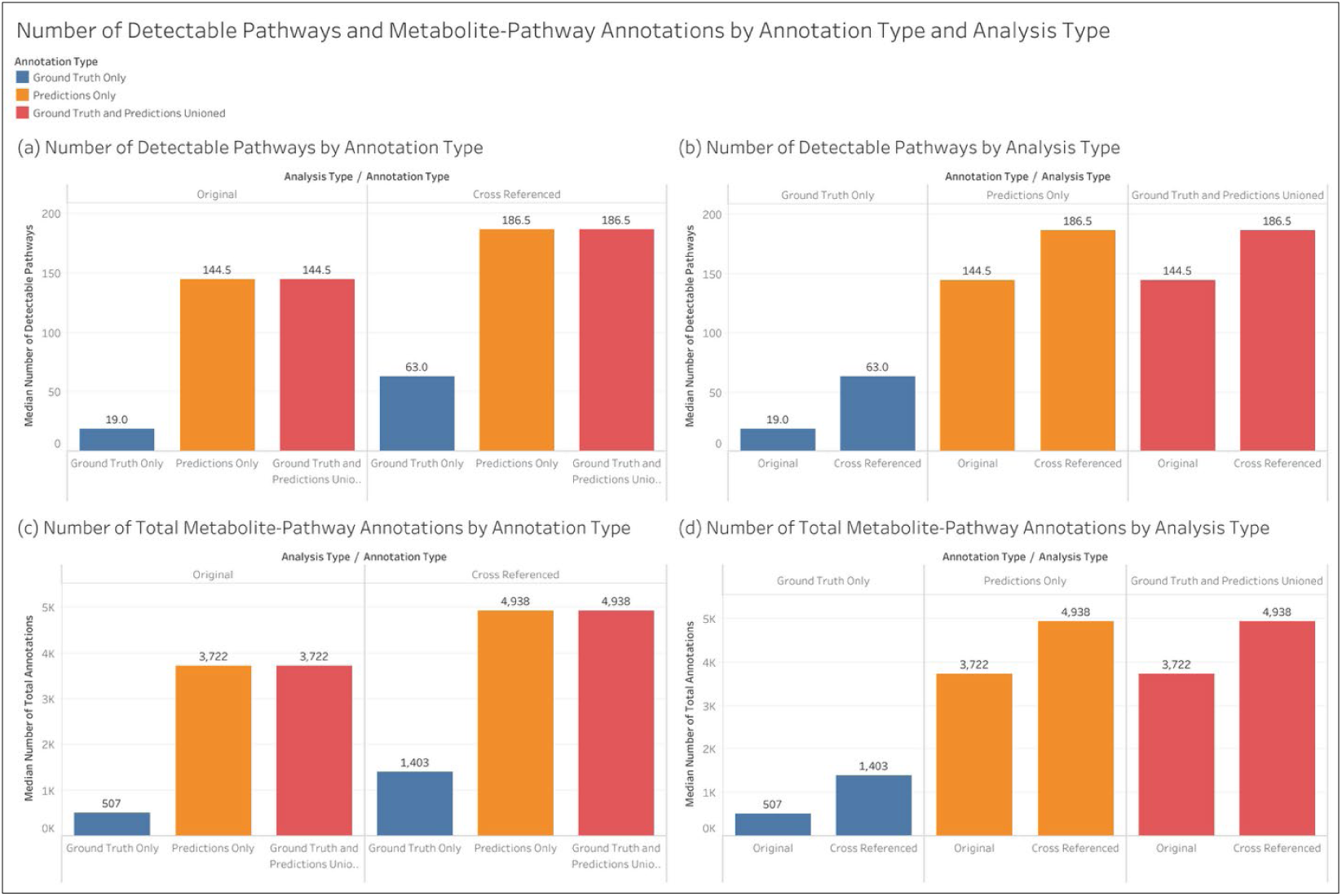
The median across MW datasets of the number of pathways that are detectable (associated with at least 15 metabolites) and the total number of metabolite-pathway associations. Results are for unique pathway definitions.

Figure 5 shows, on a per MW dataset basis, the change in detectable pathways and metabolite-pathway associations between ground truth and the ground truth unioned with predictions on log-log scatterplots. The red line in each scatterplot represents y=x with a slope of 1. We see that whether using the original or the cross-referenced compound IDs, while there are a few MW datasets that experienced little or no change, nearly all datasets experienced a meaningful increase in both the number of detectable pathways and metabolite-pathway associations. Figure S5 shows these results for individual pathway IDs. Figures S6 through S9 show this information for each knowledgebase separately. Again, the largest increases occur for MetaCyc-defined and Reactome-defined pathways, which represent over 90% of the uniquely defined pathways. Figures S10 through S13 show the same as figures S6 through S9 but for individual pathway IDs.

**Figure 5.**
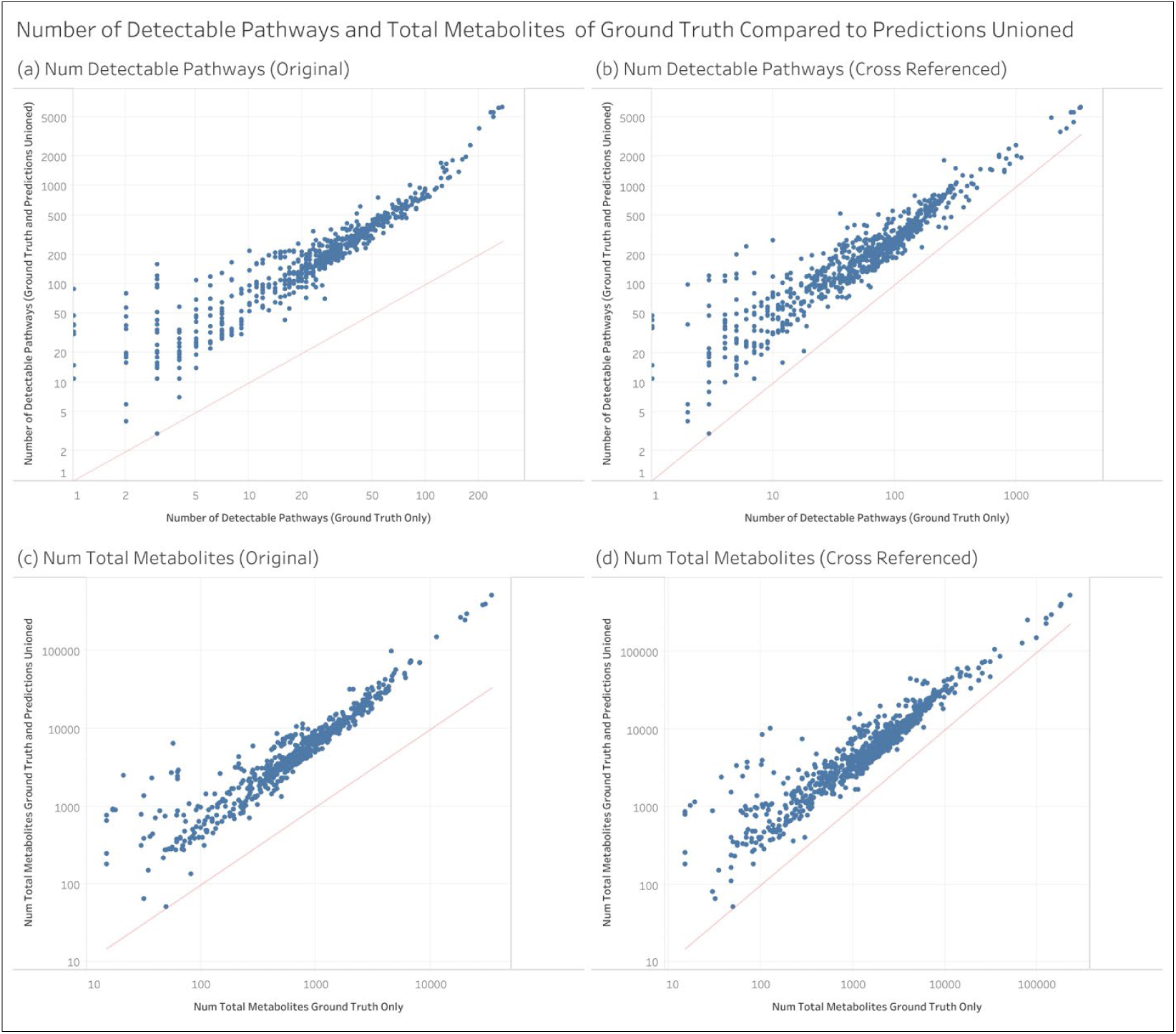
Per dataset view of the amount of increase in the number of detectable pathways and the total number of metabolite-pathway associations between ground truth annotations and the ground truth unioned with the predictions. Note that the axes are on different log scales across the 4 scatterplots. Each point is a separate MW dataset. Results are for unique pathway definitions.

### 3.2. Post-GSEA

While the above results demonstrate how the ML model’s predictions increase the number of pathways that could be detected, the following results demonstrate the actual change in enriched pathways at an adjusted p-value ≤ 0.01. Again, these results are for unique pathway definitions.

#### 3.2.1. Changes in the number of enriched pathways

Figure 6 shows the change in the median number of enriched pathways across MW datasets after adding more compound IDs via cross referencing. We see that when using the ground truth annotations alone, adding the cross references significantly increases the median number of enriched pathways. However, when using the predictions, either the predictions only or the predictions unioned with the ground truth, the cross-references do increase the median number of enriched pathways, but the change is not statistically significant. Figure S14 shows these results for individual pathway IDs.

**Figure 6.**
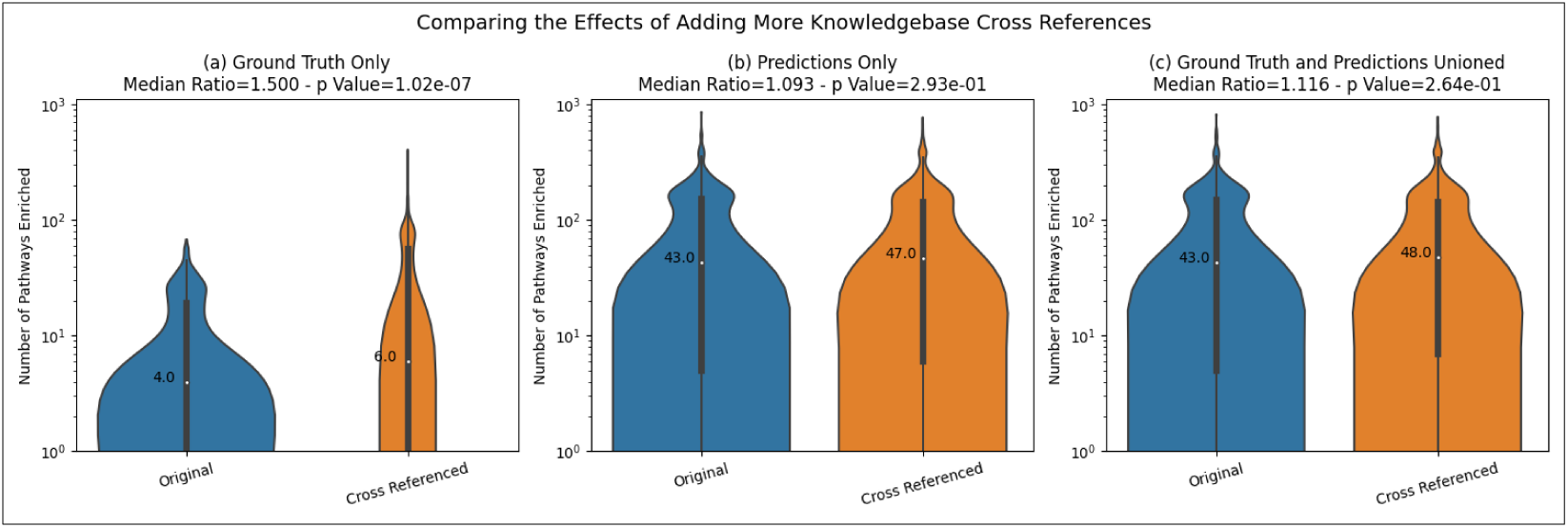
The effect on the number of enriched pathways of adding metabolite ID cross references to the MW datasets for each type of annotations. Results are for unique pathway definitions.

Figure 7 compares the number of enriched pathways when using the predictions by themselves and when unioning the ground truth with the predictions. We see that for both the original and cross-referenced MW datasets, there is not a significant difference between the two approaches. Figure S15 shows these results for individual pathway IDs.

**Figure 7.**
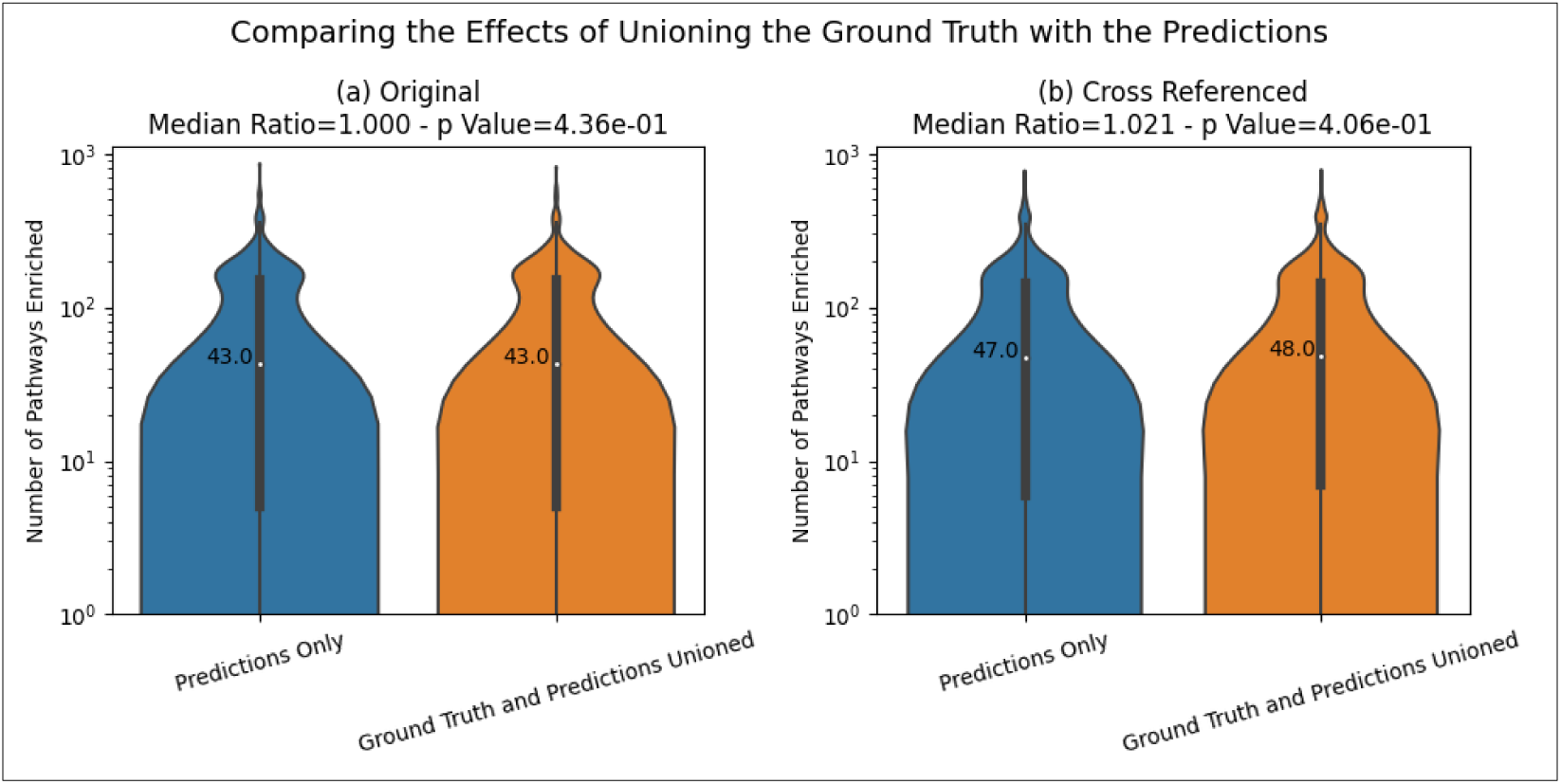
The effect on the number of enriched pathways when unioning the predictions with the ground truth. Results are for unique pathway definitions.

Figure 8 demonstrates the high quality of the ML model’s predictions by the substantial increase in the median number of enriched pathways detected at an adjusted p-value ≤ 0.01 when adding the predicted pathway annotations. Although the effect size when comparing the original datasets to the datasets with predictions (10.75 for both predictions only and predictions unioned with ground truth) is larger than the effect size when comparing the cross-referenced datasets with predictions (7.833 for predictions only and 8.0 for predictions unioned with ground truth), all effect sizes still have an over 7-fold increase in the median number of enriched pathways after adding predicted pathway annotations. Figure S16 shows these results for individual pathway IDs wherein we see an over 10-fold increase. Figures S17 and S18 show this information for each knowledgebase separately. Again, the largest increases were observed for MetaCyc and Reactome-defined pathways, which represent over 90% of the uniquely defined pathways. Figures S19 and S20 show the same results as Figures S17 and S18 but for individual pathway IDs.

**Figure 8.**
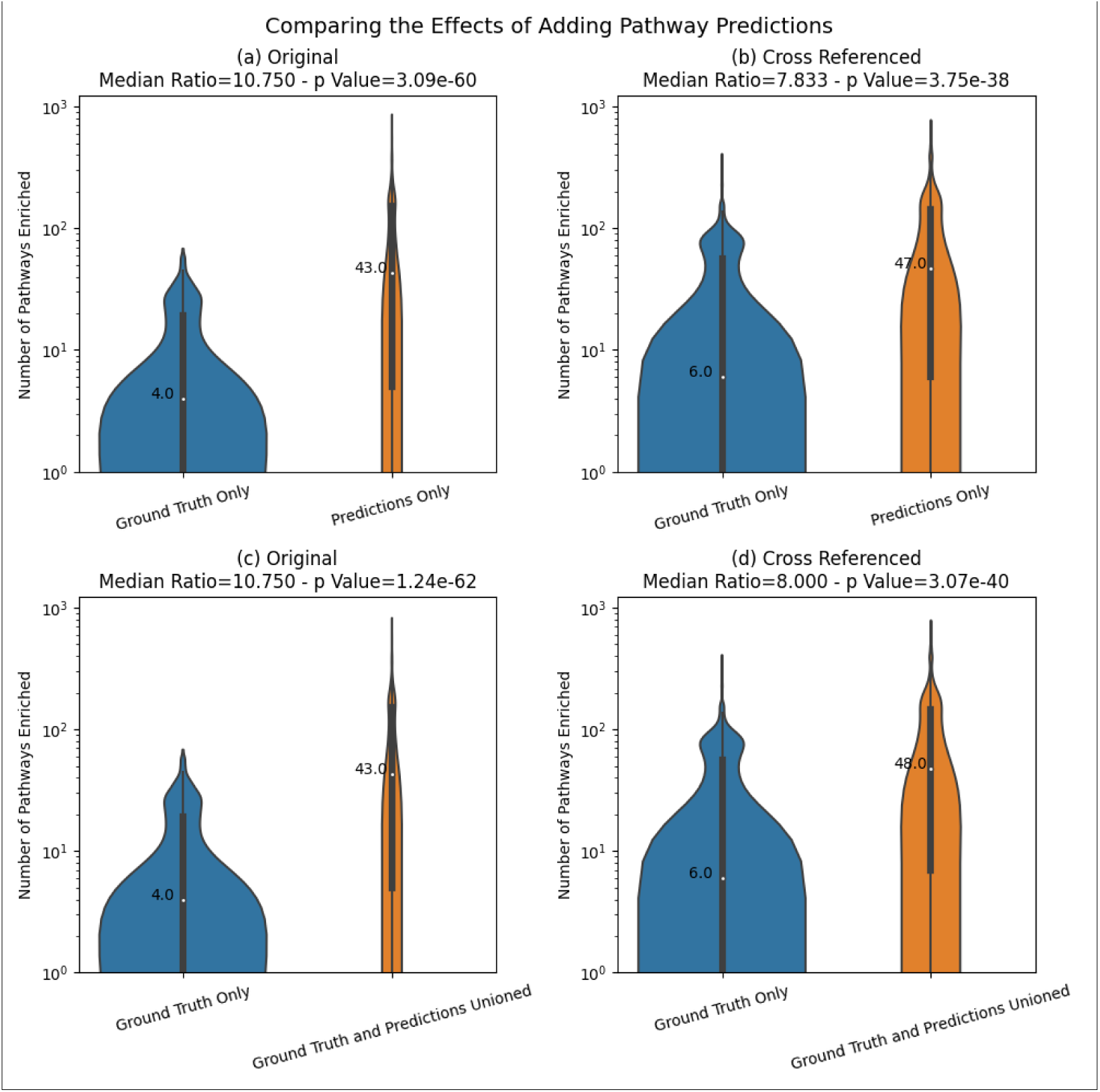
The increase of the number of enriched pathways when adding predicted annotations. Results are for unique pathway definitions.

#### 3.2.2 Information loss and gain

While the above results demonstrate that adding the predictions greatly increases the overall number of enriched pathways, in some cases, incorporating the pathway predictions results in a relatively minor decrease in the number of enriched pathways. Figure 9 shows the per MW dataset change in the number of enriched pathways between the ground truth and that unioned with predictions as log-log scatterplots. The red line is a slope of 1, meaning data points above the line indicate an increase in enriched pathways after adding predictions and those below the line indicate a decrease. Consistent with prior results, we see that a majority of MW datasets experienced an increase in the number of enriched pathways as a result of adding predictions (71.3% to 89.9%), but for some datasets, adding the predictions actually resulted in less enriched pathways than when using the ground truth alone (1.3% to 4.5%). Figure S21 shows these results for individual pathway IDs. Figures S22 and S23 show this information for each knowledgebase separately. Again, the largest increases were observed for MetaCyc and Reactomedefined pathways, which represent over 90% of the uniquely defined pathways. Figures S24 and S25 show the same results as figures S22 and S23 but for individual pathway IDs.

**Figure 9.**
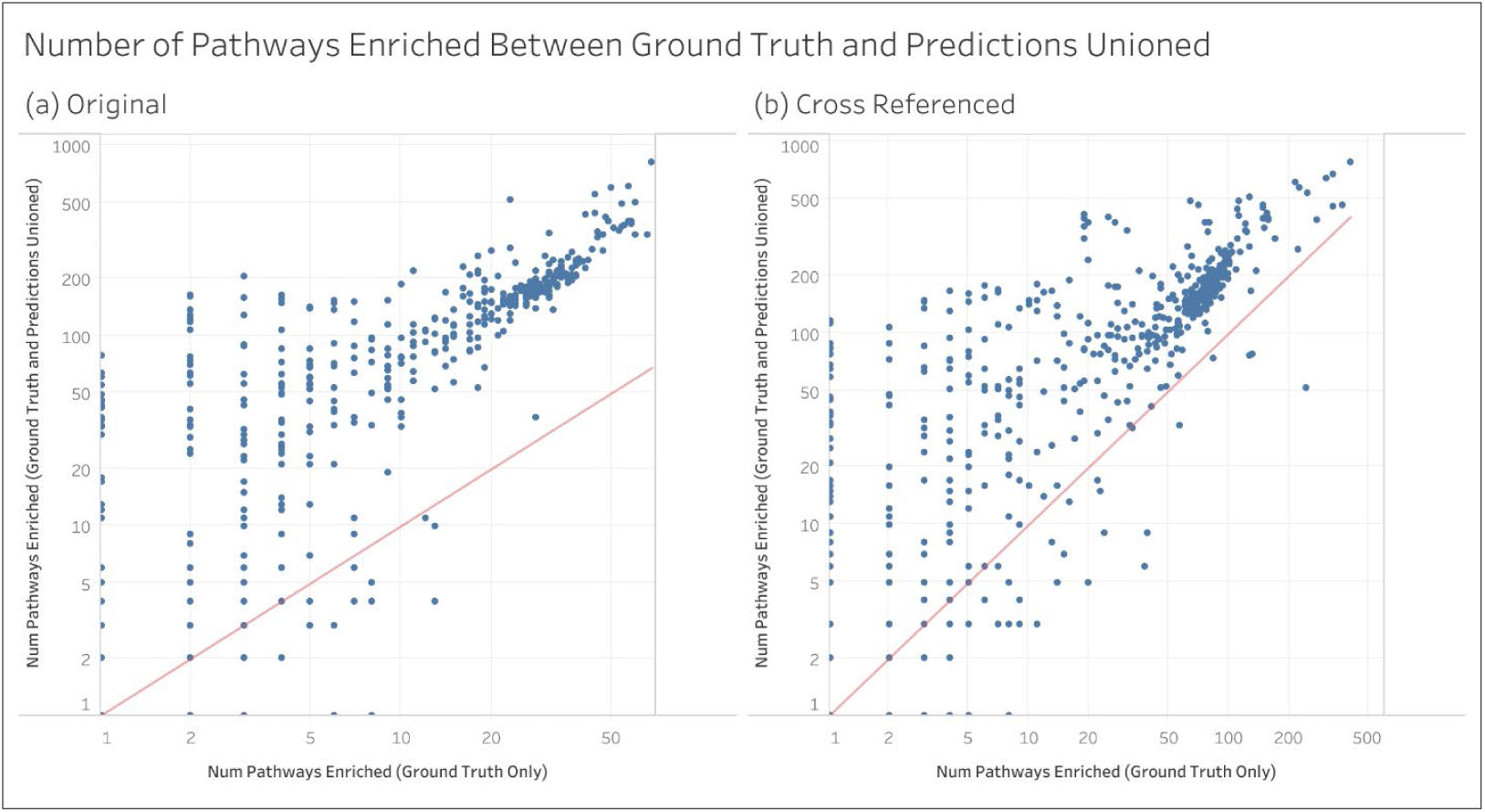
The per dataset change in the number of enriched pathways between the ground truth and the ground truth unioned with the predictions. Each point is a separate MW dataset. Results are for unique pathway definitions.

**Figure 10.**
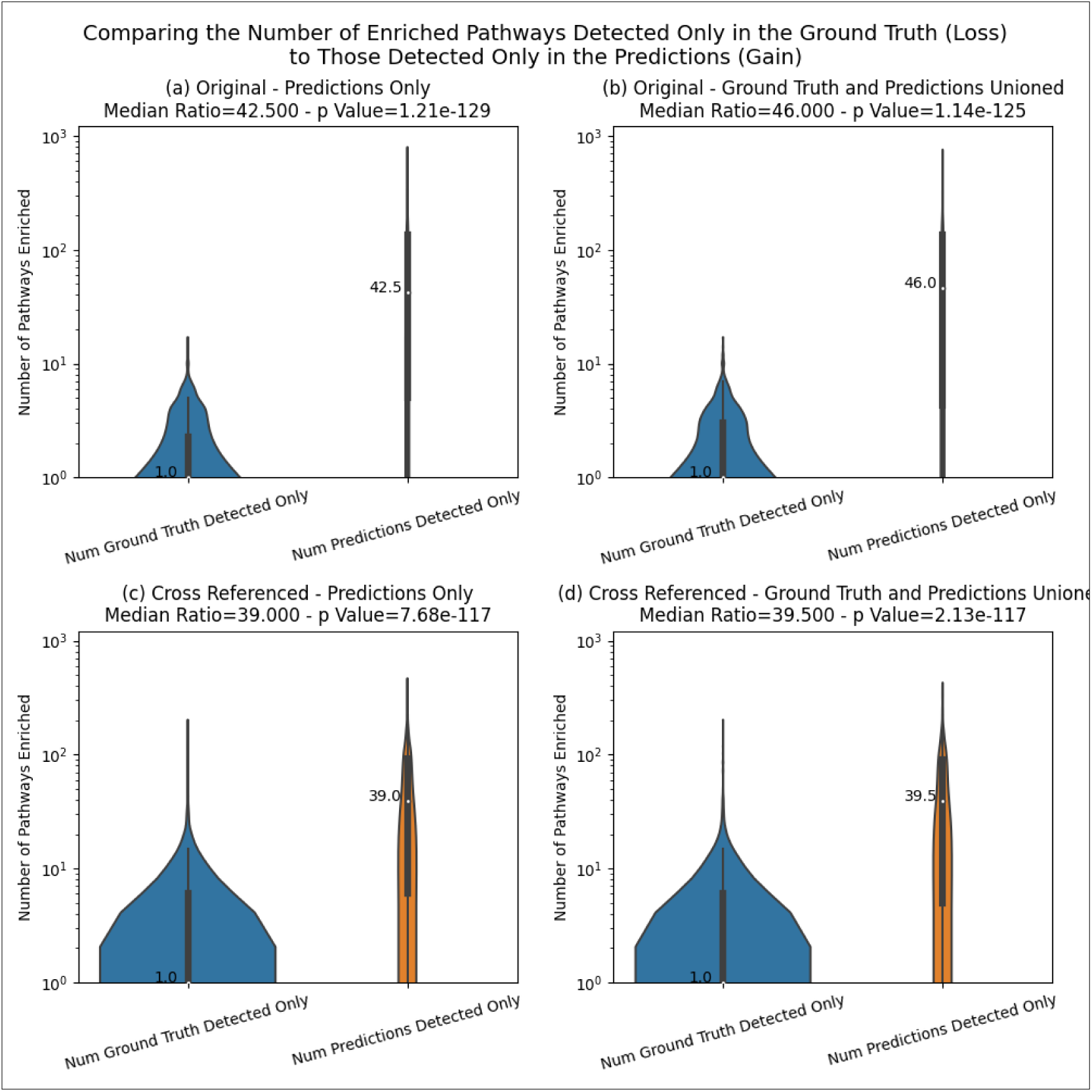
Comparing the distribution of information loss to that of information gain across MW datasets. Results are for unique pathway definitions.

Table 2 quantifies the information gain and information loss in terms of enriched pathways across all MW datasets. Information loss is indicated when a pathway is enriched in the ground truth but no longer enriched after adding the predictions for a given MW dataset. Likewise, information gain is indicated when a pathway is not enriched in the ground truth but is enriched after adding the predictions for a given MW dataset. There is no change in information when a pathway is enriched from both sets of annotations for a given MW dataset. Total information loss, total information gain, and total unchanged information is summed up across all pathways across all MW datasets. Adding up these three sums equals the total number of enriched pathways across all MW datasets. The total can then be used to calculate the percentage of information loss, information gain, and unchanged information. Table 2 shows that whether using the original MW datasets or the cross-referenced datasets and whether using the predictions only or those unioned with ground truth, information gain has the highest percentage by far with the percent unchanged coming in second. However, we do observe a small amount of information loss across MW datasets as a result of adding the predictions. Table S2 shows these results for individual pathway IDs.

**Table 2.**
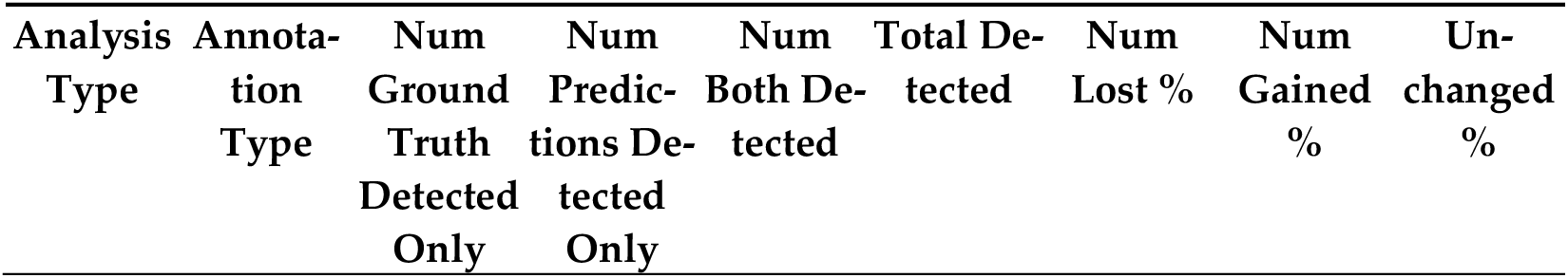
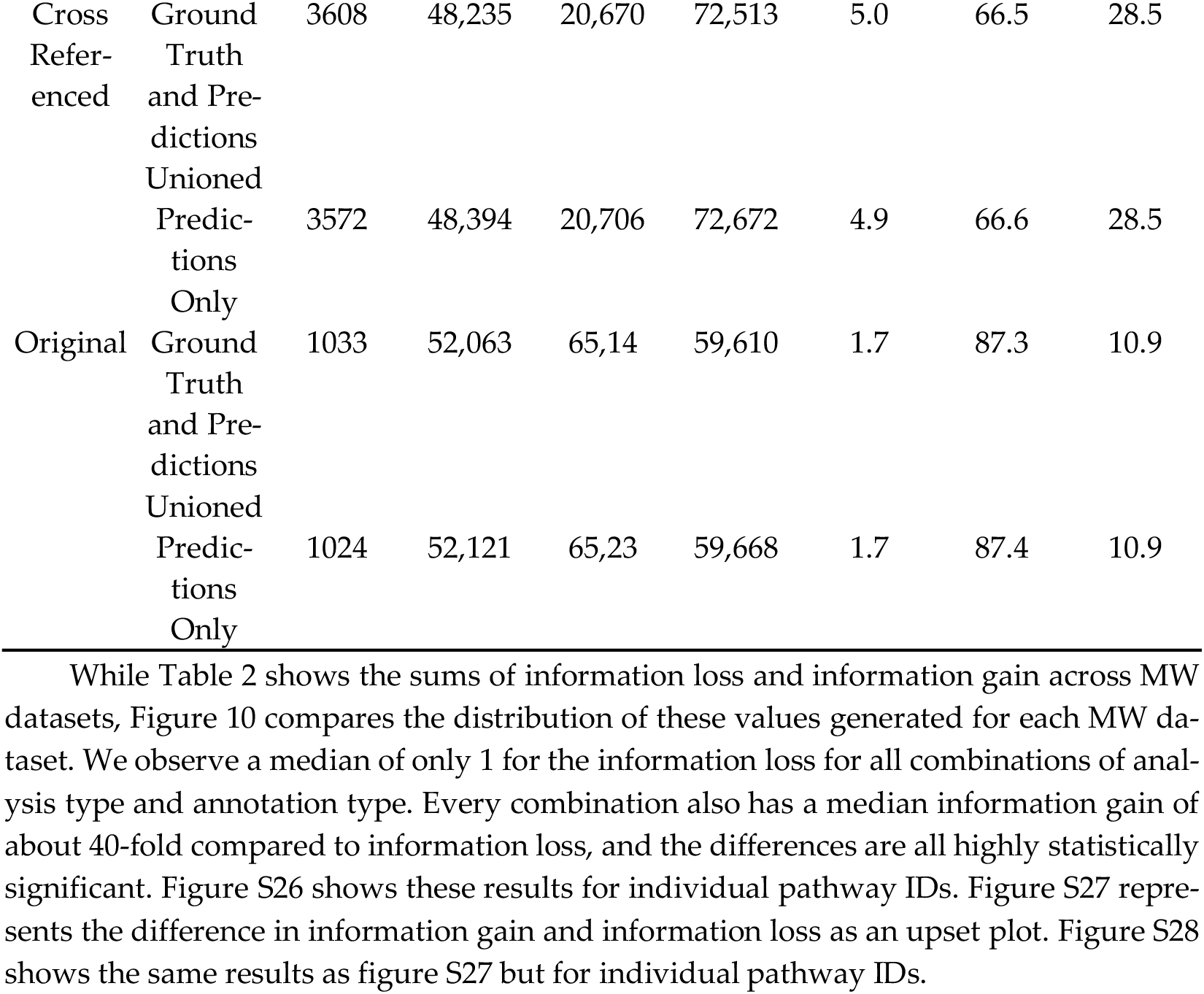
The amount of information gain and information loss summed across all enriched pathways and MW datasets. Results are for unique pathway definitions.

In Figure 11, each pathway’s overall MCC, as obtained in the work of Huckvale and Moseley [38], is plotted against the sum of the number of times the pathway was enriched across all MW datasets only after adding predicted pathway annotations, but not enriched when using ground truth annotations alone. We see that for all combinations of original or cross-referenced datasets and predictions only or predictions unioned with ground truth, the number of times that pathways are enriched greatly increases when the predictive performance of pathways reaches an MCC of 0.7. We see an increase in the number of times enriched between pathways below an overall MCC of 0.7 and pathways equal to or above an overall MCC of 0.7 and the difference between the two groups are highly statistically significant (all p-values from a Mann-Whitney U test ≤ 10-19), with mean ratios ranging from 5.5– to 5.8-fold. We use mean ratios here since the median number of times enriched for pathways below an overall MCC of 0.7 was consistently 0, making it impossible to calculate ratios using the medians. These results illustrate a strong dependence of pathway enrichment on pathway prediction performance. Specifically, high enrichment does not occur until pathway prediction overall MCC is roughly 0.7 or above.

**Figure 11.**
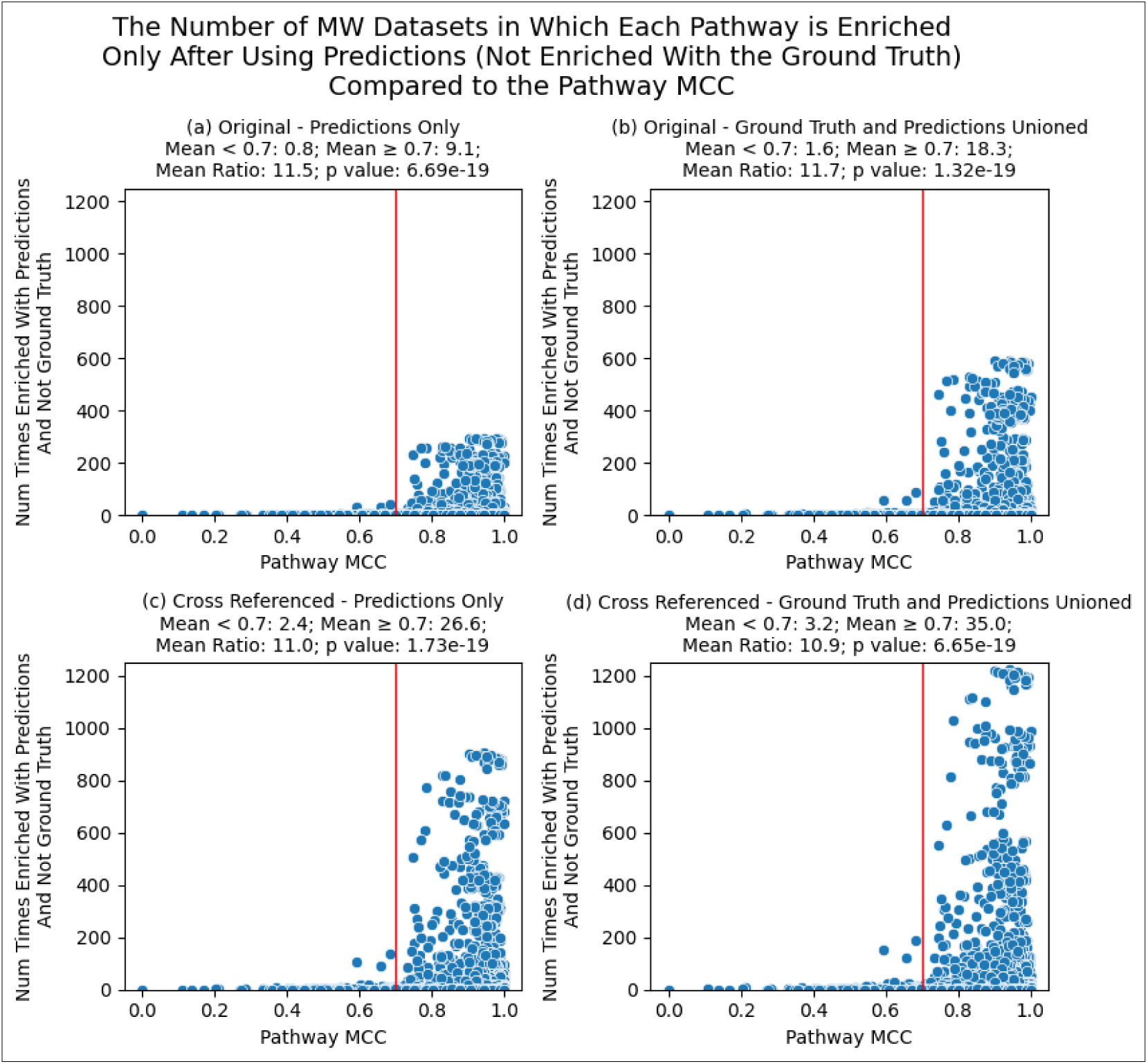
Comparing the pathway MCC to the number of MW datasets wherein each pathway was not enriched when using ground truth annotations but was enriched when using predicted annotations. The red line is drawn at overall MCC = 0.7. Each point is a different pathway enriched in at least one dataset. Results are for unique pathway definitions.

Table 3 demonstrates the coverage of metabolism in terms of the number of unique metabolites present within the metabolite-pathway annotations across MW datasets before and after using predictions. We see that the increase in the number of unique metabolites is greater for the original datasets as compared to the cross referenced datasets. Note that the number of datasets used to compute the summary statistics is less than 990 since datasets that had 0 metabolites present for both ground truth and predictions were filtered out. Table S3 shows these stats for all MW datasets unfiltered. Also note that it does not matter whether pathway IDs or pathway definitions are used since these counts are the union of pathway definitions (associated metabolites) and redundant pathway definitions leave the union unchanged.

**Table 3.**
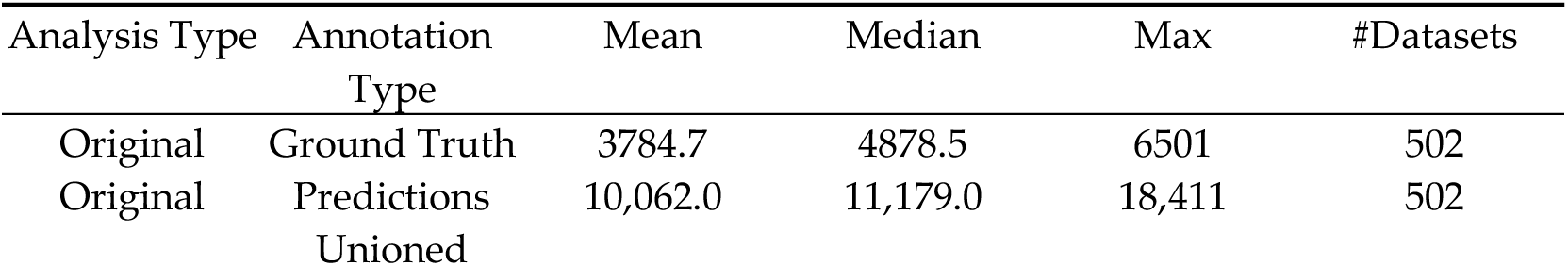
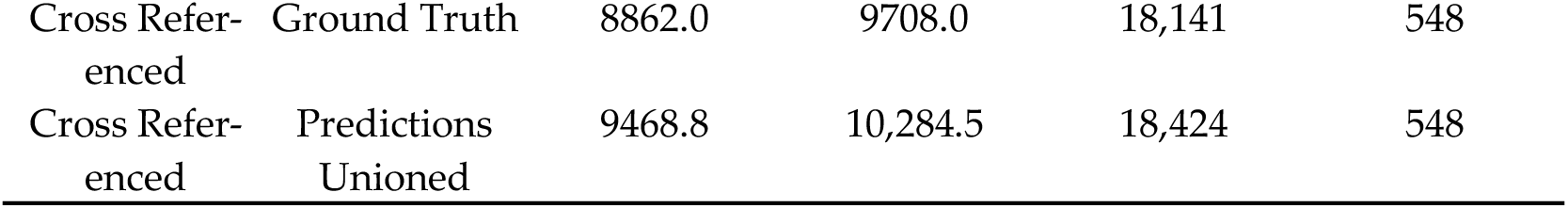
The number of unique metabolites as part of metabolite-pathway annotations across MW datasets. Datasets with 0 metabolites present for both ground truth and predictions are filtered out.

Table 4 demonstrates information gain by quantifying the uniqueness of enriched pathways across MW datasets by calculating Jaccard indices of their associated definitions (associated metabolites). Specifically, this is calculated by summing the pathway uniqueness scores across enriched pathways in an MW dataset, the uniqueness score being calculated by averaging 1 minus the Jaccard distance between its definition to the definition of every other enriched pathway in the dataset. Again, datasets with scores of 0 for both ground truth and predictions are filtered out (Table S4 shows these results unfiltered) and it does not matter whether pathway IDs or pathway definitions are used since redundant definitions would add a 1 minus Jaccard index of 0 to the uniqueness score.

**Table 4.**
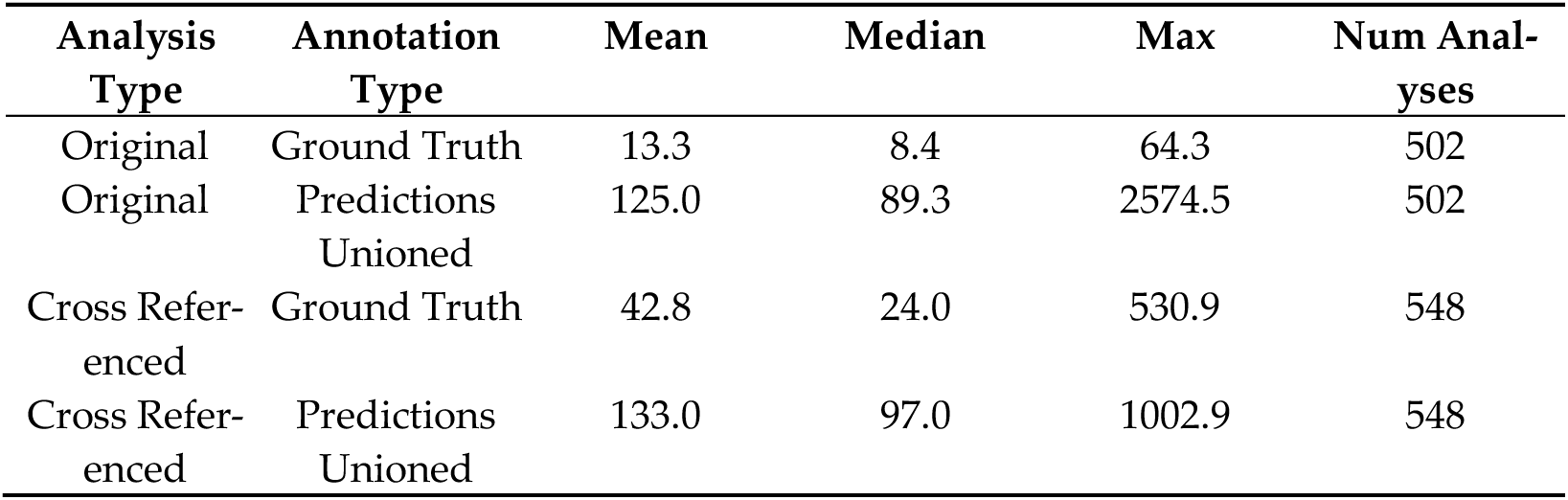
Sum of 1 minus average Jaccard index between each enriched pathway definition and every other enriched pathway definition. Datasets with 0 enriched pathways in both ground truth and predictions are filtered out.

| Analysis<br>Type | Annotation<br>Type | Mean | Median | Max | Num Anal-<br>yses |
| --- | --- | --- | --- | --- | --- |
| Original | Ground Truth | 13.3 | 8.4 | 64.3 | 502 |
| Original | Predictions<br>Unioned | 125.0 | 89.3 | 2574.5 | 502 |
| Cross Refer-<br>enced | Ground Truth | 42.8 | 24.0 | 530.9 | 548 |
| Cross Refer-<br>enced | Predictions<br>Unioned | 133.0 | 97.0 | 1002.9 | 548 |

## 4. Discussion

Our results demonstrate the high utility of using machine learning to predict pathway annotations that enhance the results from PEA. Using an ML model with the highest prediction performance to date along with the highest number of pathways capable of being predicted, we greatly increased the number of pathway annotations and pathways that are detectable (Figure 4 and Figure 5). The quality of the model’s pathway predictions, whether predicting unique pathway definitions or pathway IDs, is indicated by its ability to increase the number of pathways detected i.e. pathways being associated with changes in metabolites across test groups at a statistically significant extent (Figure 8). We expect the 8-fold increase in the number of enriched unique pathways has high potential for expanding the biological and biomedical interpretability and insight of metabolomics datasets.

We consider the ability to accurately predict a higher number of pathway annotations to be a substantial milestone in resolving the lack of compound IDs in metabolomics datasets and the lack of pathway annotations for the corresponding metabolites in knowledgebases such as KEGG, MetaCyc, and Reactome. We also demonstrate that the issues surrounding the lack of compound IDs and associated pathway IDs can be ameliorated by chaining cross references. However, we further demonstrate increases in the number of enriched pathways far beyond what the crossreferencing can accomplish alone. From Figure 8, we see an over 7-fold increase in enriched pathways whether or not cross-referencing is performed in the pipeline, as well as an over 7-fold increase in enriched pathways whether or not ground truth annotations are available at all. This is especially useful for researchers that either do not have access to compound ID cross-references or do not have access to pathway annotations at all, especially for duplicate metabolite entries of different compound IDs where one ID has pathway annotations and the other does not or where intra-knowledgebase cross references are not provided by the knowledgebase. These findings are additionally useful when a compound ID is not available since the model can predict pathway annotations from the chemical structure information alone.

We see that while the ML model’s predictions improve the results of PEA overall, there are some cases of information loss alongside the information gain (Figure 9 and Table 2). While the amount of information gain is about 40-fold higher than information loss (Figure 10), we expect the few cases of information loss to be attributed mainly to inaccuracies in the predictions of the model. This is somewhat corroborated by the higher gains in Reactome vs KEGG and MetaCyc, with Reactome pathway predictions having an overall MCC of 0.94 versus 0.87 for KEGG and 0.88 for MetaCyc [35],[36],[37],[38], providing evidence that more accurate predictions result in more enriched pathways. More specifically, Figure 11 shows a pathway’s potential to be enriched increases between 10.9 and 11.5 fold when reaching a predictive performance of at least 0.7 overall MCC. As of now, we recommend that researchers analyzing metabolomics datasets try both the ground truth annotations with chaining cross-references, if they have access to them, as well as our model’s predictions to gain the largest number of enriched pathways while avoiding any information loss. As an additional precaution, predictions for pathways with overall MCC less than 0.7 can simply be excluded. We anticipate that further improvements to the performance and generalizability of the model, especially to increase the number of pathways with overall MCC ≥ 0.7, will likely result in less information loss as well as higher information gain, further improving the interpretability of and insight from metabolomics analyses. Beyond information gain in the increased number of enriched pathways, we also see a gain in the amount of unique metabolic information (Table 4).

## 5. Conclusions

Our results demonstrate the utility of an ML model that predicts pathway annotations. The increased amount of pathway annotations greatly expands the discovery and interpretability of metabolomics datasets via PEA.

## Supplementary Materials

Table S1: The number of MW datasets filtered by reason for filtering; Figure S1: Violin plots comparing the number of detectable pathways and total number of metabolite-pathway annotations for the combined data; Figure S2: The median across MW datasets of the number of pathways that are detectable (associated with at least 15 metabolites) and the total number of metabolite-pathway associations. Results are for individual pathway IDs; Figure S4: Comparing the median number of detectable pathways and total metabolite-pathway annotations between original and cross-referenced MW datasets and between ground truth and predictions for each knowledgebase. Results are for unique pathway definitions; Figure S4: Same as Figure S3 but for individual pathway IDs; Figure S5: Per dataset view of the amount of increase in the number of detectable pathways and the total number of metabolite-pathway associations between ground truth annotations and the ground truth unioned with the predictions. Note that the axes are on different scales across the 4 scatterplots. Each point is a separate MW dataset. Results are for individual pathway IDs; Figure S7: Per analysis view of the change in the number of detectable pathways between the ground truth and that unioned with predictions for each knowledgebase using cross-referenced MW datasets. Results are for unique pathway definitions. Results are for unique pathway definitions; Figure S7: Per analysis view of the change in the number of detectable pathways between the ground truth and that unioned with predictions for each knowledgebase using the original MW datasets. MetaCyc and Reactome are not shown here since there were not any metabolite IDs available for these in the original datasets. Results are for unique pathway definitions; Figure S8: Per analysis view of the change in the number of metabolite-pathway annotations between the ground truth and that unioned with predictions for each knowledgebase using cross-referenced MW datasets. Results are for unique pathway definitions. Results are for unique pathway definitions; Figure S9: Per analysis view of the change in the number of metabolite-pathway annotations between the ground truth and that unioned with predictions for each knowledgebase using the original MW datasets. MetaCyc and Reactome are not shown here since there were not any metabolite IDs available for these in the original datasets. Results are for unique pathway definitions; Figure S10 – Same as Figure S6 but for individual pathway IDs; Figure S11: Same as Figure S7 but for individual pathway IDs; Figure S12: Same as Figure S8 but for individual pathway IDs; Figure S13: Same as Figure S9 but for individual pathway IDs. Figure S15: The effect on the number of enriched pathways of adding metabolite ID cross references to the MW datasets for each type of annotations. Results are for individual pathway IDs. Figure S15: The effect on the number of enriched pathways when unioning the predictions with the ground truth. Results are for individual pathway IDs; Figure S17: The increase of the number of enriched pathways when adding predicted annotations. Results are for individual pathway IDs; Figure S17: Comparing the distribution across MW analyses of the number of enriched pathways between the ground truth pathway annotations and those unioned with the predicted annotations for the cross referenced MW datasets for each knowledgebase. Combined results included for comparison. Results are for unique pathway definitions; Figure S18: Comparing the distribution across MW analyses of the number of enriched pathways between the ground truth pathway annotations and those unioned with the predicted annotations for the original MW datasets for each knowledgebase. Note that MetaCyc and Reactome had no ground truth annotations available since their metabolite IDs were not available in the original MW datasets. Combined results included for comparison. Results are for unique pathway definitions. Results are for unique pathway definitions; Figure S19: Same as Fiigure S17 but for individual pathway IDs; Figure S20: Same as Figure S18 but for individual pathways; Figure S21: The per dataset change in the number of enriched pathways between the ground truth and the ground truth unioned with the predictions. Each point is a separate MW dataset. Results are for individual pathway IDs; Figure S22: Per analysis view of the change in the number of enriched pathways between the ground truth pathway annotations and those unioned with the predicted annotations for each knowledgebase on the cross-referenced MW datasets. Results are for unique pathway definitions; Figure S23: Per analysis view of the change in the number of enriched pathways between the ground truth pathway annotations and those unioned with the predicted annotations for each knowledgebase on the original MW datasets. Note that MetaCyc and Reactome are not shown because these had no metabolite IDs available in the original datasets. Results are for unique pathway definitions; Figure S24: Same as Figure S22 but for individual pathway IDs; Figure S25: Same as Figure S23 but for individual pathway IDs; Table S2: The amount of information gain and information loss summed across all enriched pathways and MW datasets. Results are for individual pathway IDs; Figure S26: Comparing the distribution of information loss to that of information gain across MW datasets. Results are for individual pathway IDs; Figure S27: Represents the information loss and information gain of enriched pathways between ground truth and predictions by showing the overlap of detected pathways. Results are for unique pathway definitions. Results are for unique pathway definitions; Figure S28: Same as Figure S27 but for individual pathway IDs; Figure S29: Comparing the pathway MCC to the number of MW datasets wherein each pathway was not enriched when using ground truth annotations but was enriched when using predicted annotations. The red line is drawn at overall MCC = 0.7. Each point is a different pathway enriched in at least one dataset. Results are for individual pathway IDs; Table S3: The number of unique metabolites as part of metabolite-pathway annotations across MW datasets; Table S4: Sum of 1 minus average Jaccard distance between enriched pathway definitions and every other pathway definition; Table S5: Hyperparameters used for the ML model.

## Author Contributions

Conceptualization, EDH and HNBM; methodology, EDH, PTT, and RMF; software, EDH, PTT, and RMF; validation, EDH; formal analysis, EDH, PTT, RMF; investigation, EDH and HNBM; resources, EDH and HNMB; data curation, EDH and PTT; writing—original draft preparation, EDH and PTT; writing—review and editing, EDH, PTT, RMF, and HNBM; visualization, EDH; supervision, HNBM; project administration, HNBM; funding acquisition, HNBM. All authors have read and agreed to the published version of the manuscript.”

## Informed Consent Statement

Not applicable.

## Data Availability Statement

All data and code for reproducing the results of this manuscript are available in the following Figshare item: https://doi.org/10.6084/m9.figshare.30645875.

## Acknowledgments

We thank the University of Kentucky Institute for Biomedical Informatics, University of Kentucky Center for Computational Sciences, and National Science Foundation grant number 1626364 for their support and associated computing resources.

## Abbreviations

The following abbreviations are used in this manuscript:

PEA: Pathway Enrichment Analysis
AEA: Annotation Enrichment Analysis
KEGG: Kyoto Encyclopedia of Genes and Genomes
InChI: IUPAC International Chemical Identifier
ML: Machine Learning
MLP: Multilayer Perceptron
MCC: Matthew’s Correlation Coefficient
MW: Metabolomics Workbench
PCA: Principal Component Analysis
PC: Principal Component
GSEA: Gene Set Enrichment Analysis

## Supplemental Material for

### Filtering Counts

**Table S1.**
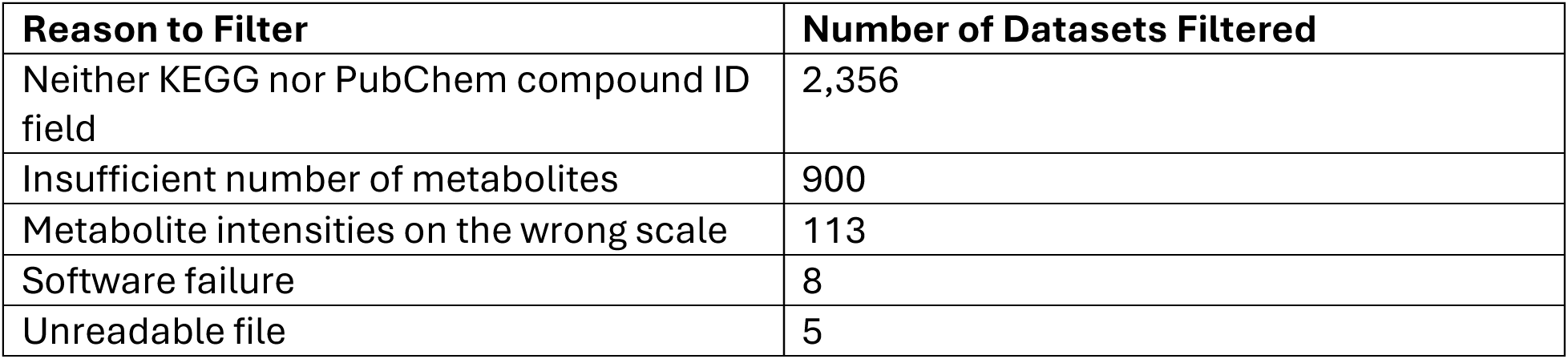
– The number of MW datasets filtered by reason for filtering.

| Reason to Filter | Number of Datasets Filtered |
| --- | --- |
| Neither KEGG nor PubChem compound ID field | 2,356 |
| Insufficient number of metabolites | 900 |
| Metabolite intensities on the wrong scale | 113 |
| Software failure | 8 |
| Unreadable file | 5 |

### Pre-GSEA

**Figure S1.**
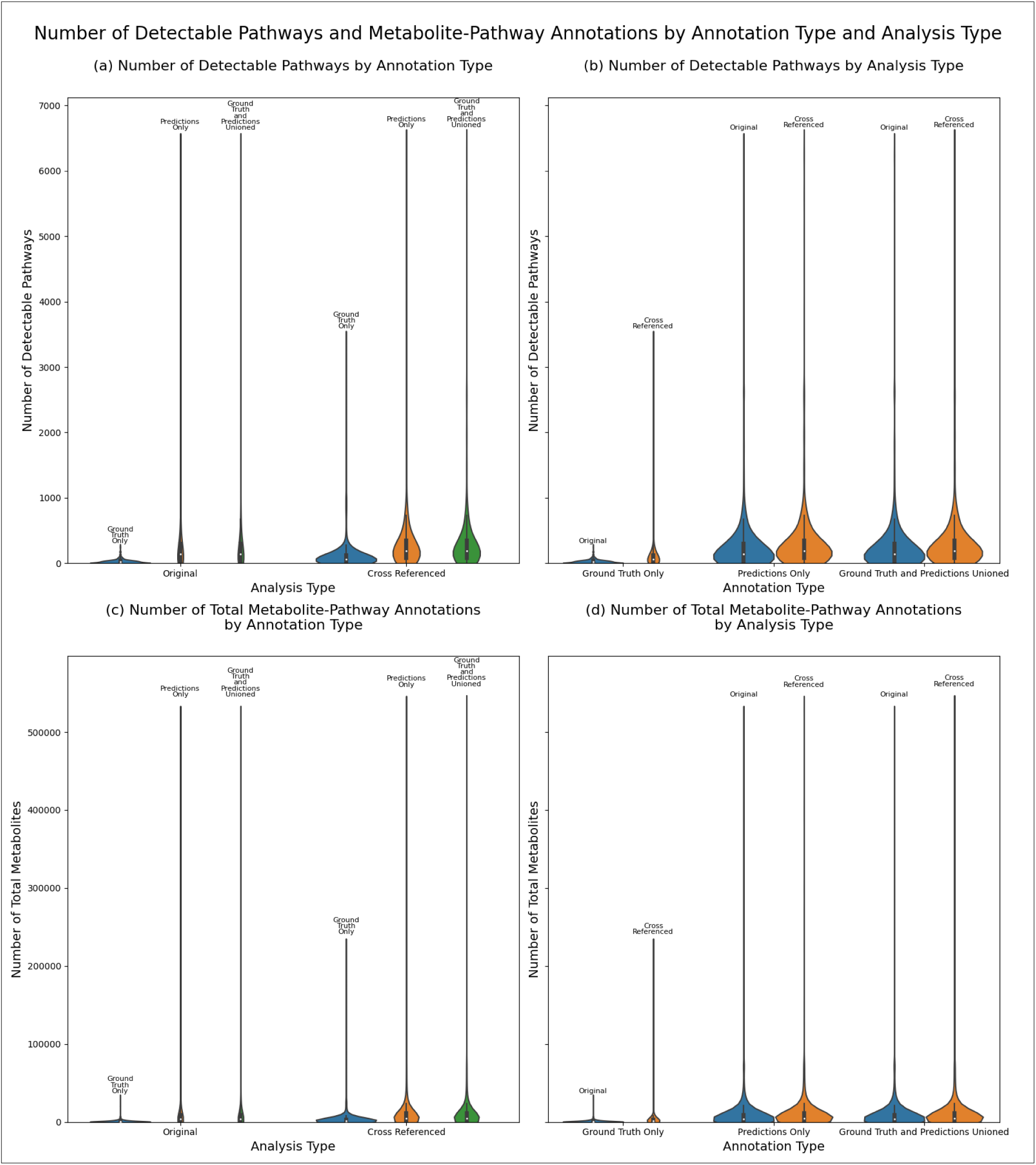
– Violin plots comparing the number of detectable pathways and total number of metabolite-pathway annotations for the combined data.

**Figure S2.**
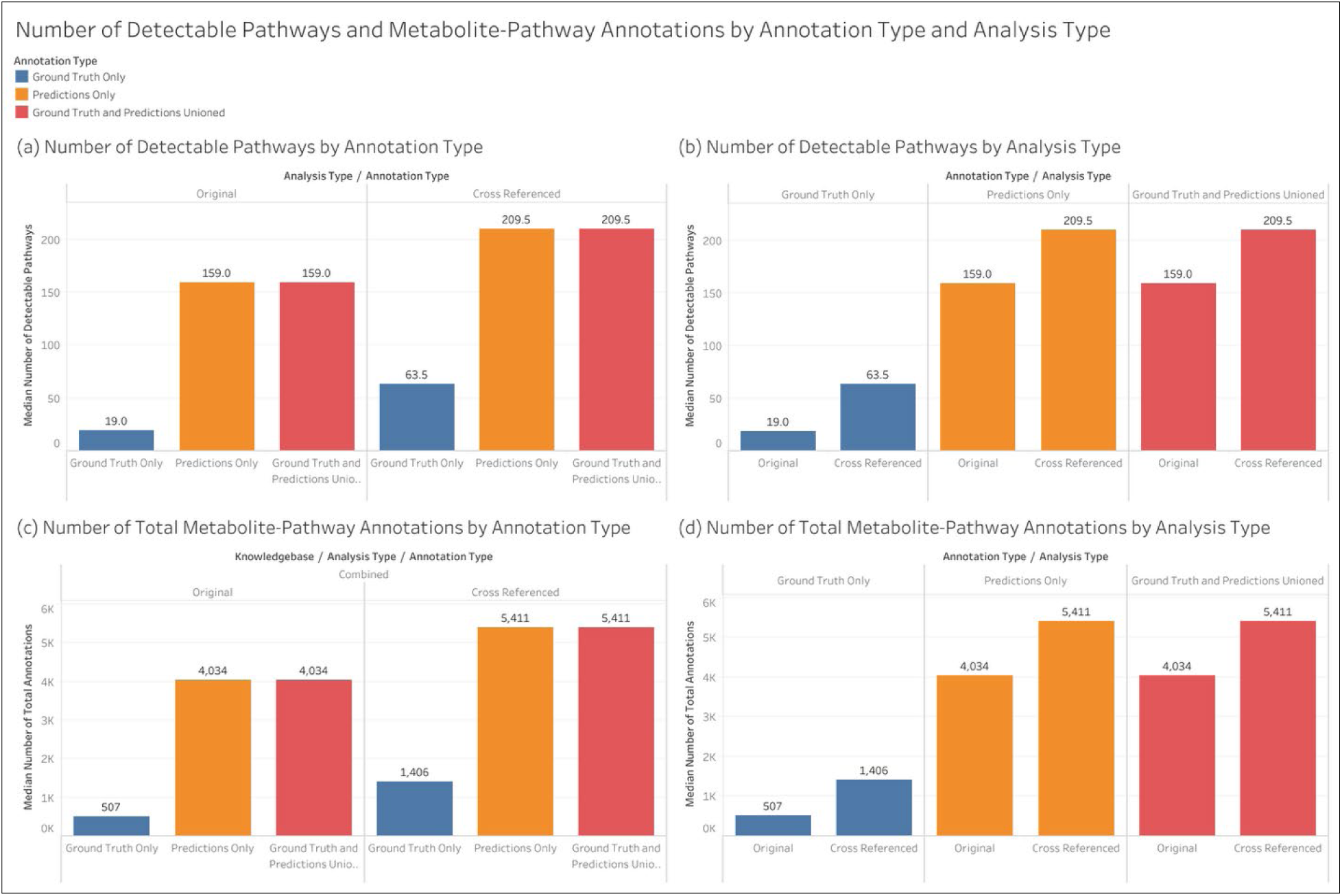
– The median across MW datasets of the number of pathways that are detectable (associated with at least 15 metabolites) and the total number of metabolite-pathway associations. Results are for individual pathway IDs.

**Figure S3.**
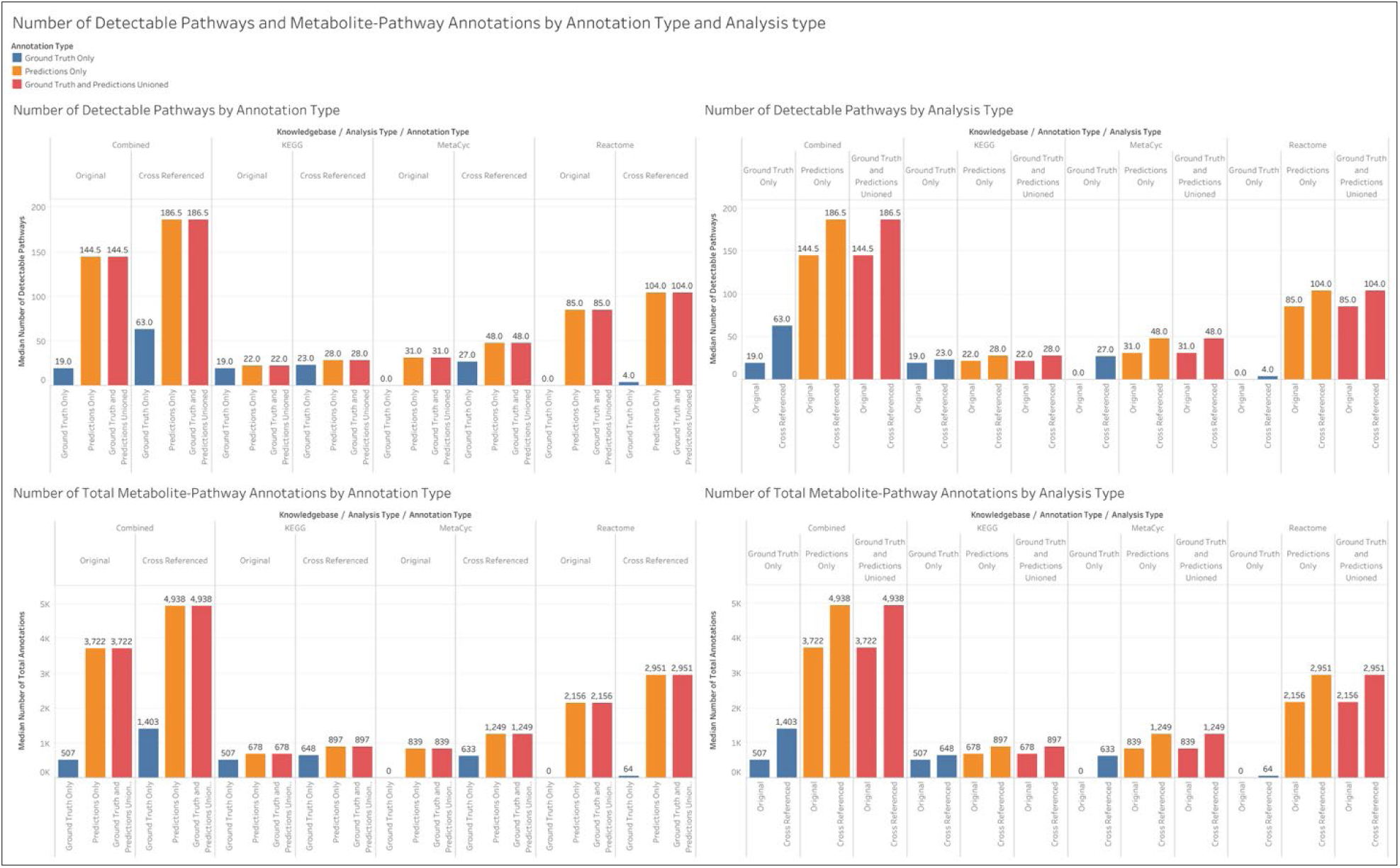
– Comparing the median number of detectable pathways and total metabolite-pathway annotations between original and cross-referenced MW datasets and between ground truth and predictions for each knowledgebase. Results are for unique pathway definitions.

**Figure S4.**
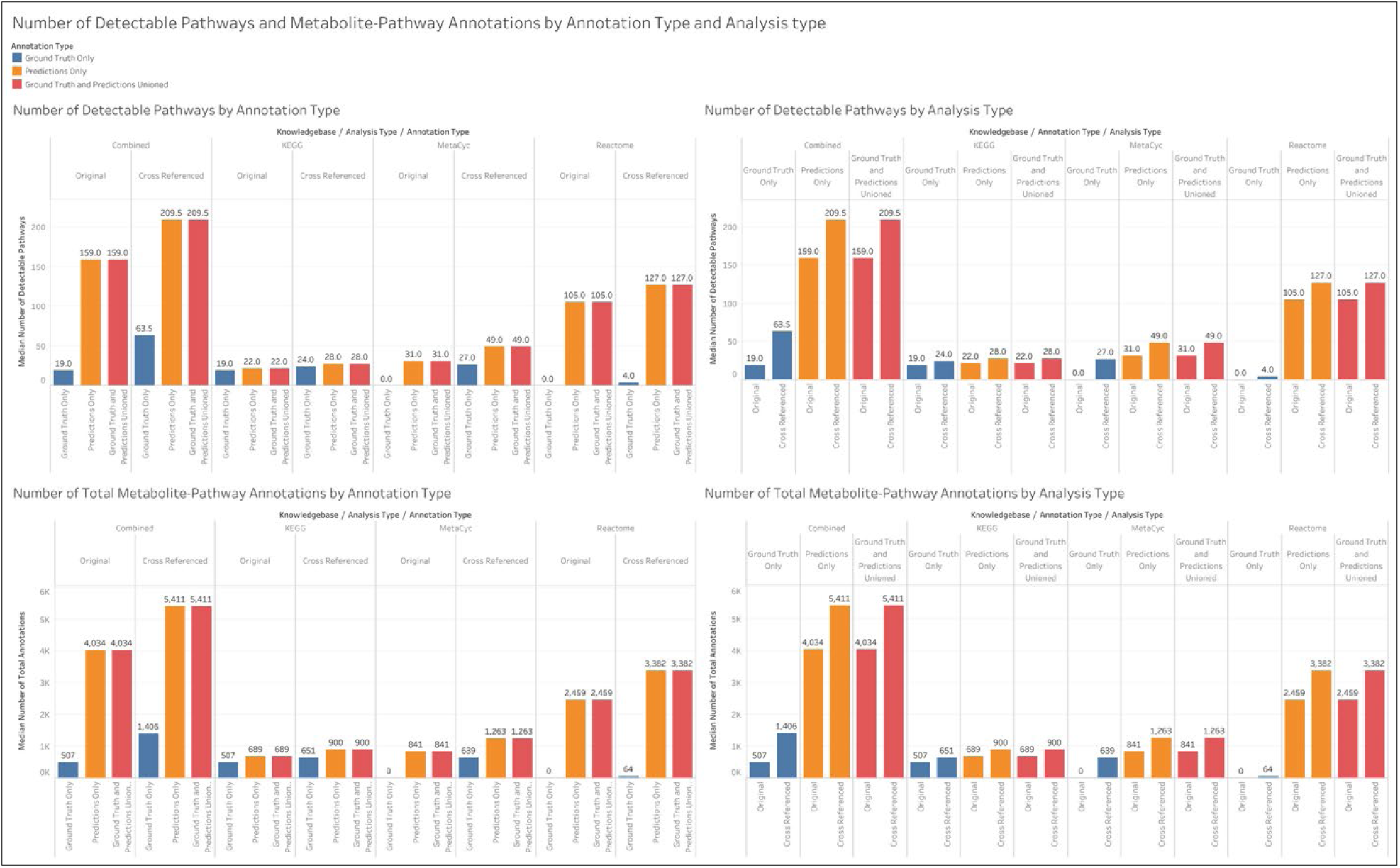
– Same as Figure S3 but for individual pathway IDs.

**Figure S5.**
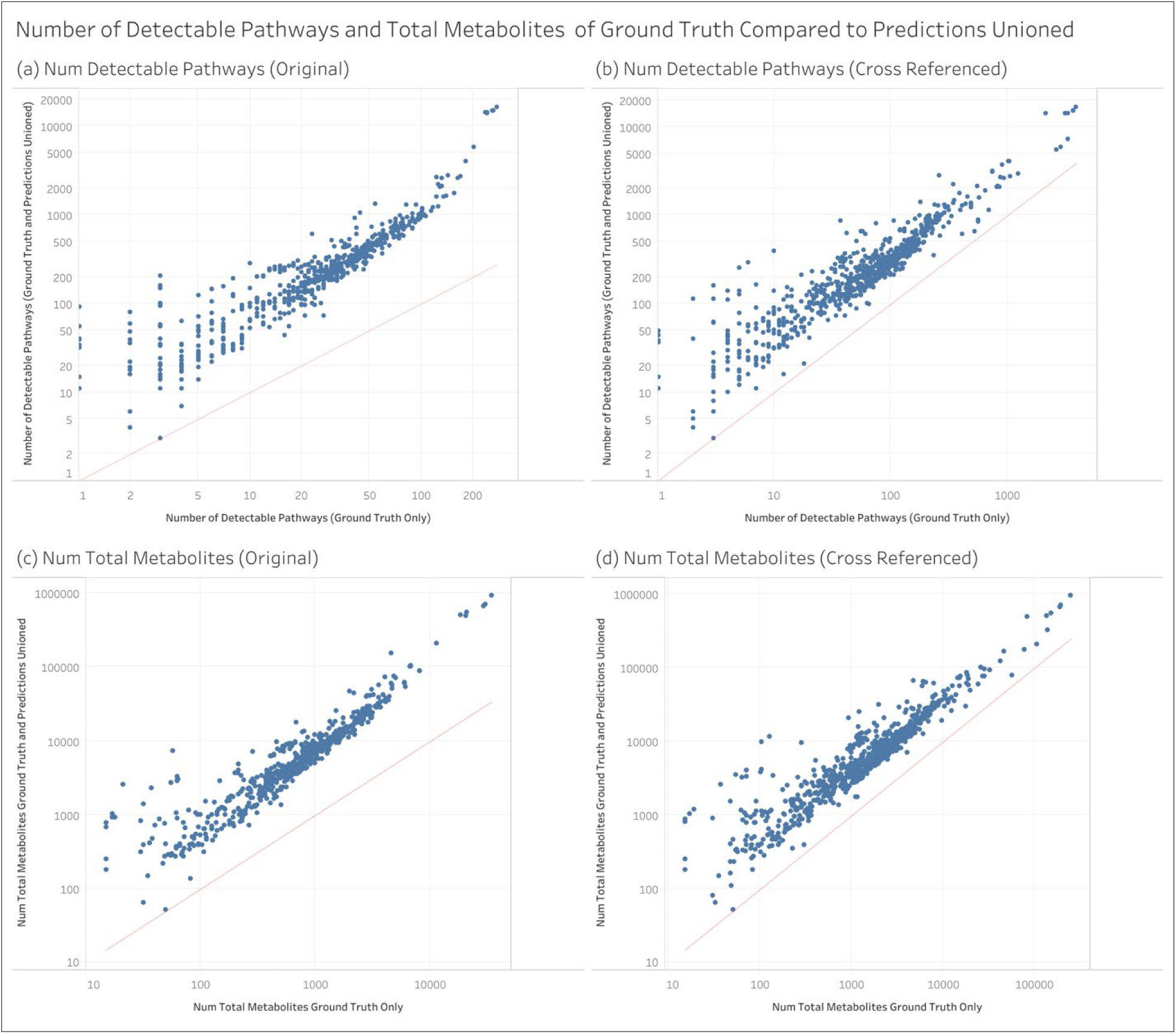
– Per dataset view of the amount of increase in the number of detectable pathways and the total number of metabolite-pathway associations between ground truth annotations and the ground truth unioned with the predictions. Note that the axes are on different scales across the 4 scatterplots. Each point is a separate MW dataset. Results are for individual pathway IDs.

**Figure S6.**
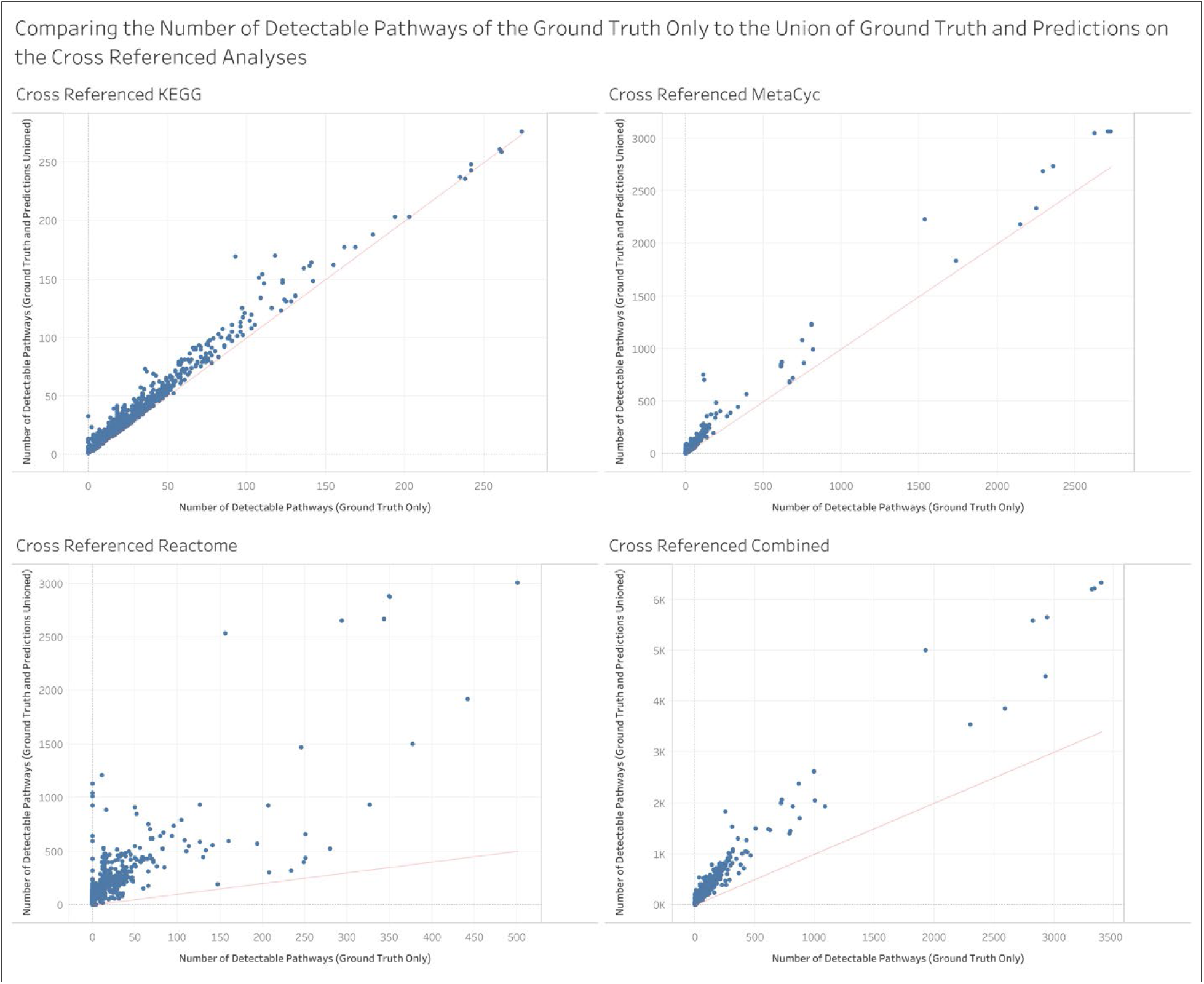
– Per analysis view of the change in the number of detectable pathways between the ground truth and that unioned with predictions for each knowledgebase using cross-referenced MW datasets. Results are for unique pathway definitions.

**Figure S7.**
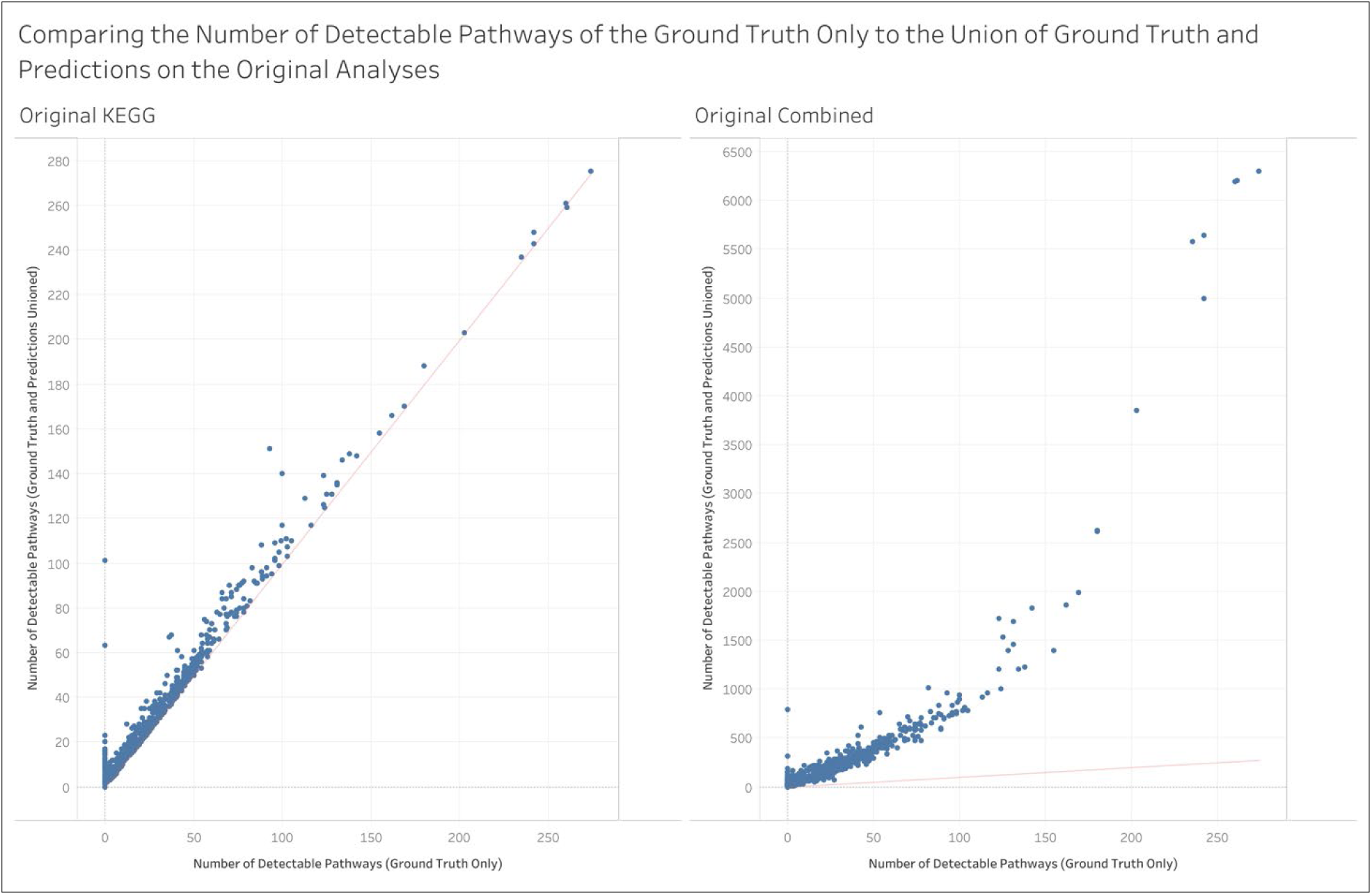
– Per analysis view of the change in the number of detectable pathways between the ground truth and that unioned with predictions for each knowledgebase using the original MW datasets. MetaCyc and Reactome are not shown here since there were not any metabolite IDs available for these in the original datasets. Results are for unique pathway definitions.

**Figure S8.**
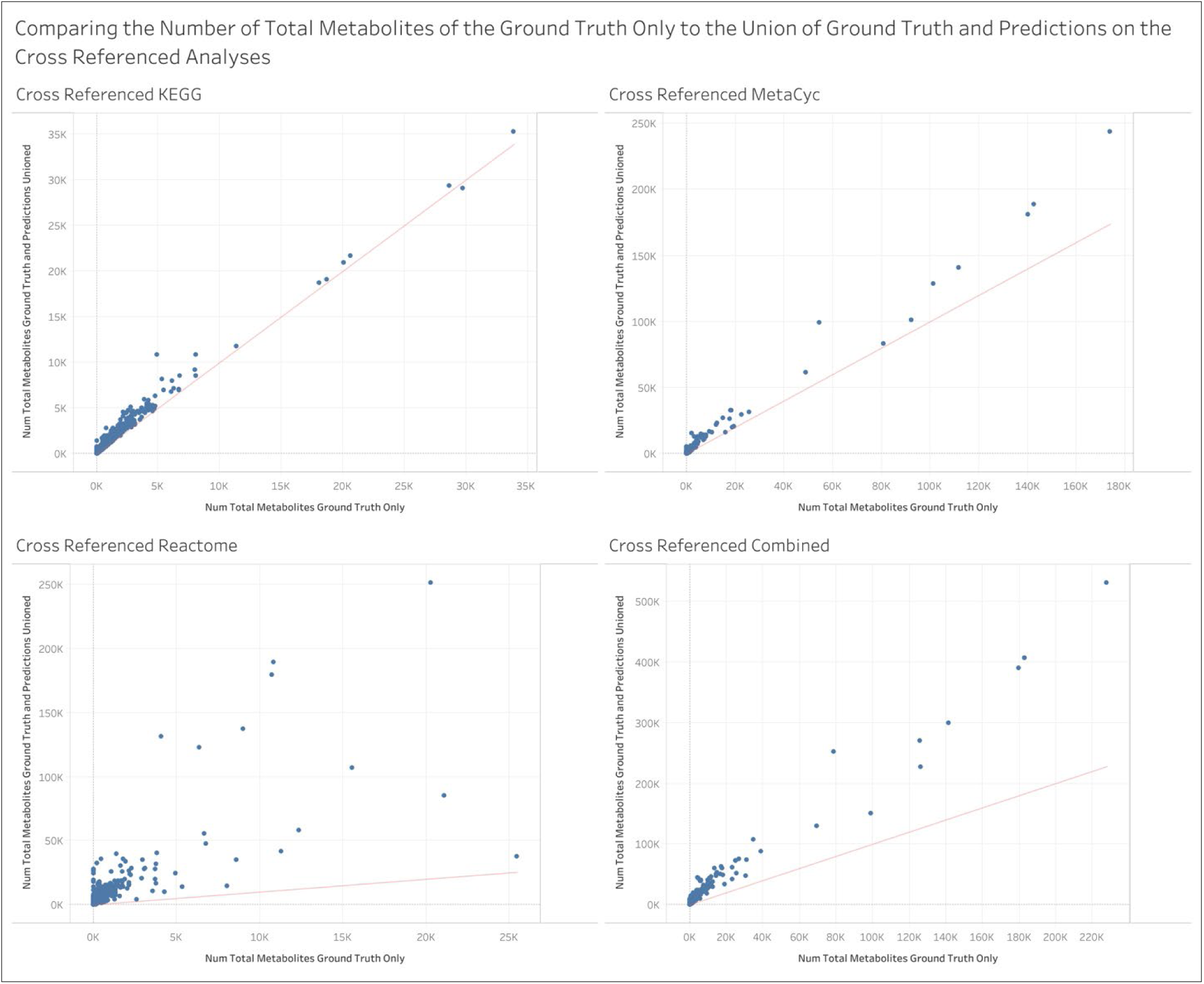
– Per analysis view of the change in the number of metabolite-pathway annotations between the ground truth and that unioned with predictions for each knowledgebase using cross-referenced MW datasets. Results are for unique pathway definitions.

**Figure S9.**
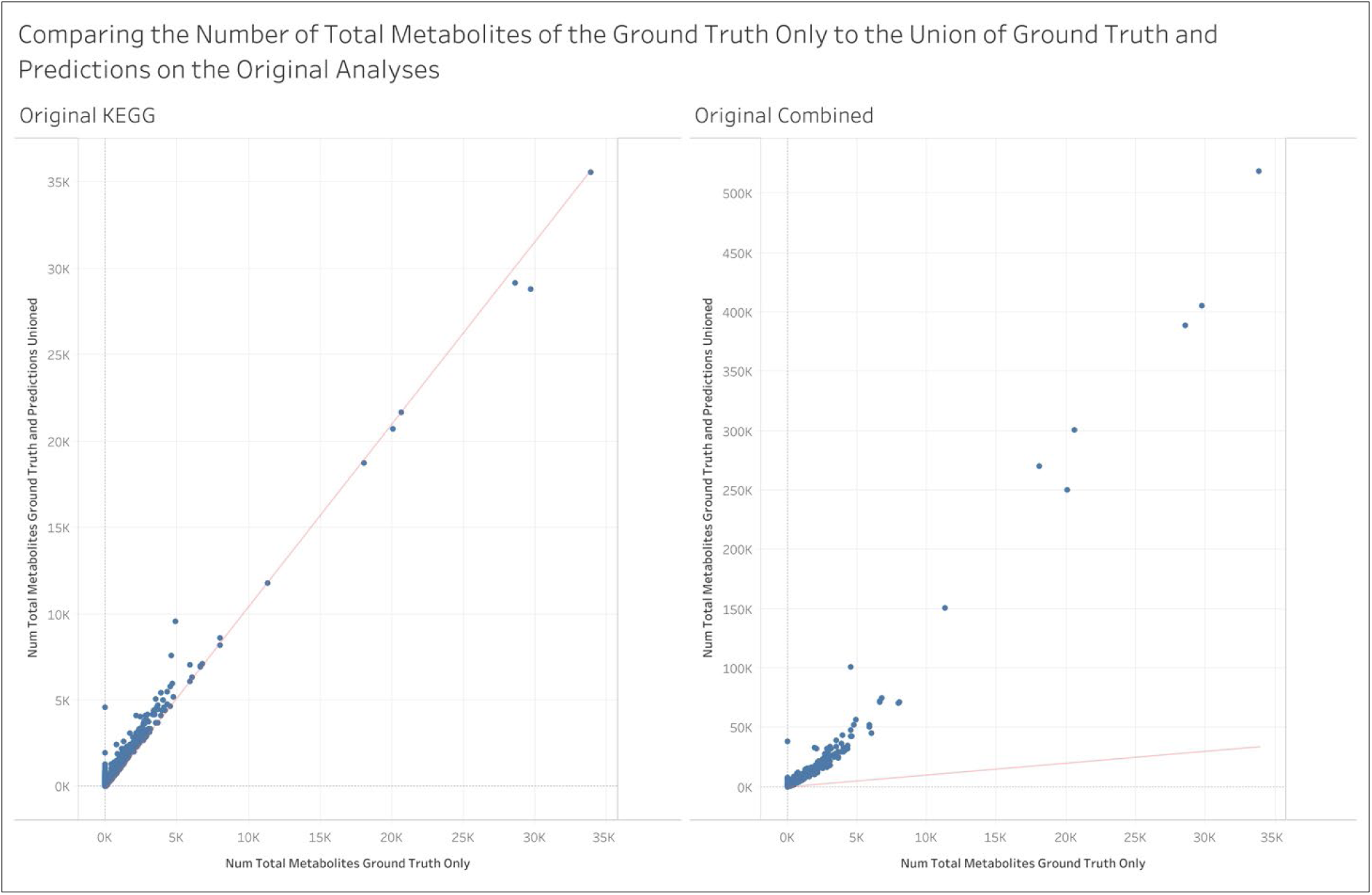
– Per analysis view of the change in the number of metabolite-pathway annotations between the ground truth and that unioned with predictions for each knowledgebase using the original MW datasets. MetaCyc and Reactome are not shown here since there were not any metabolite IDs available for these in the original datasets. Results are for unique pathway definitions.

**Figure S10.**
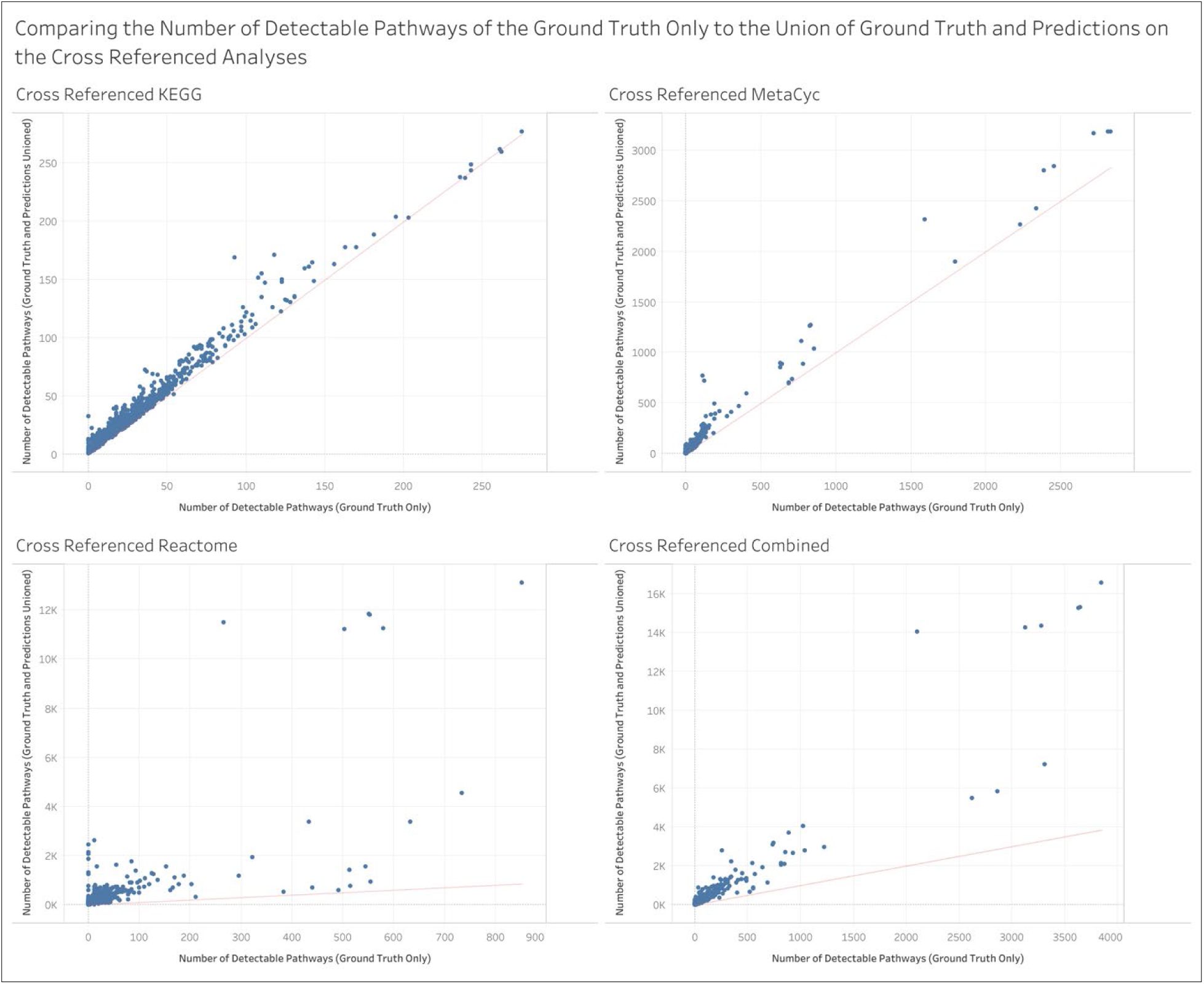
– Same as Figure S6 but for individual pathway IDs.

**Figure S11.**
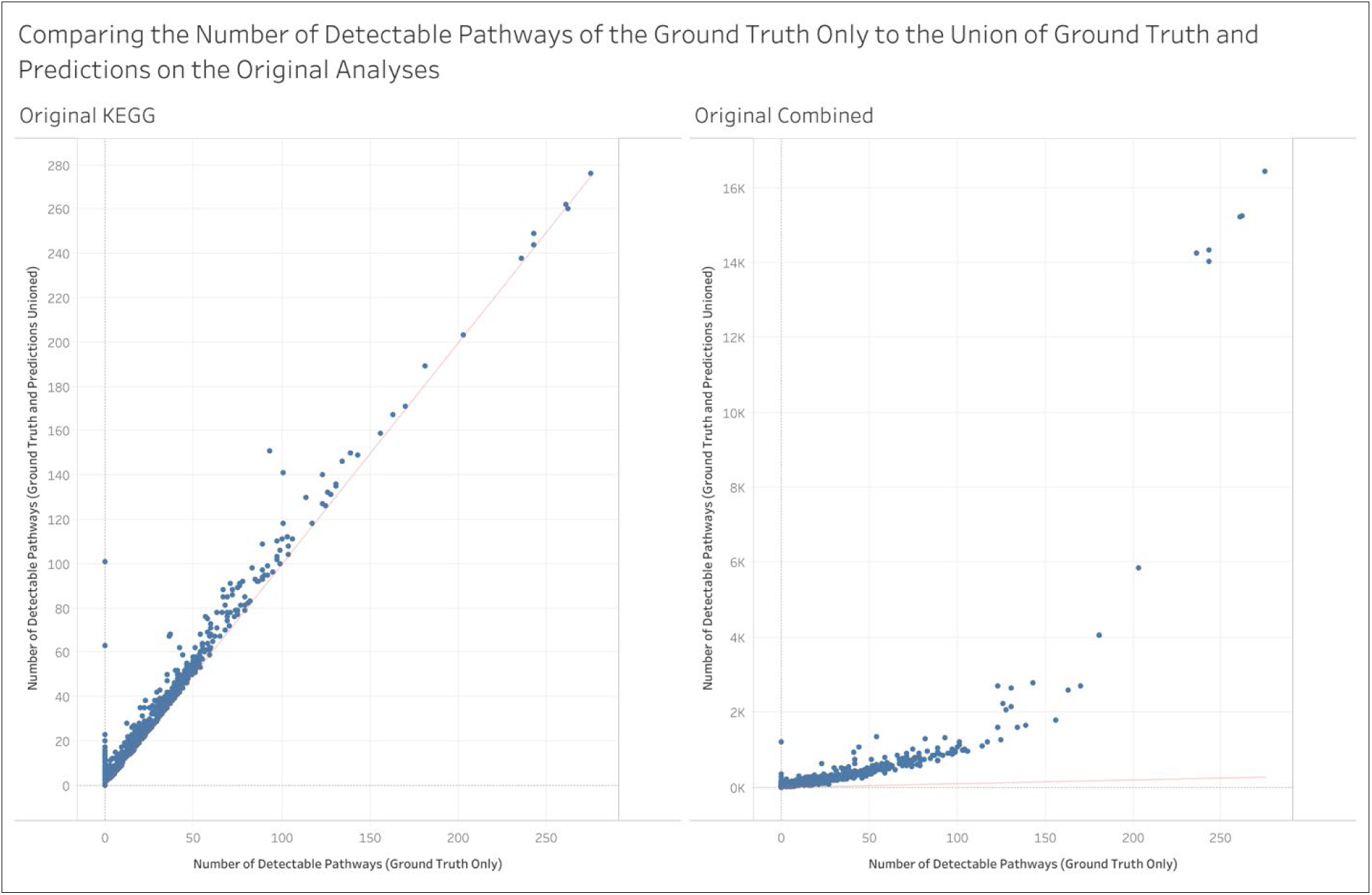
– Same as Figure S7 but for individual pathway IDs.

**Figure S12.**
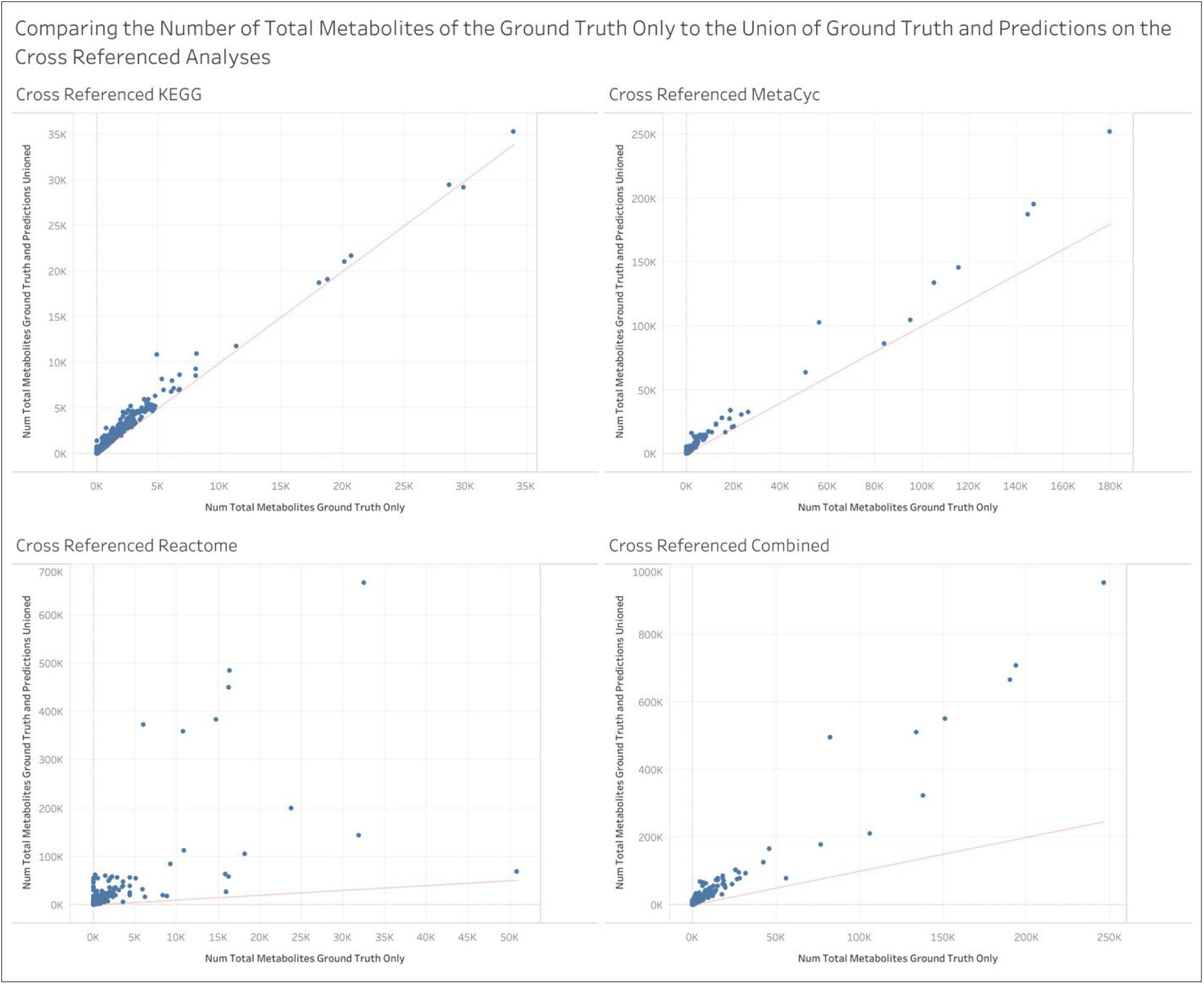
– Same as Figure S8 but for individual pathway IDs.

**Figure S13.**
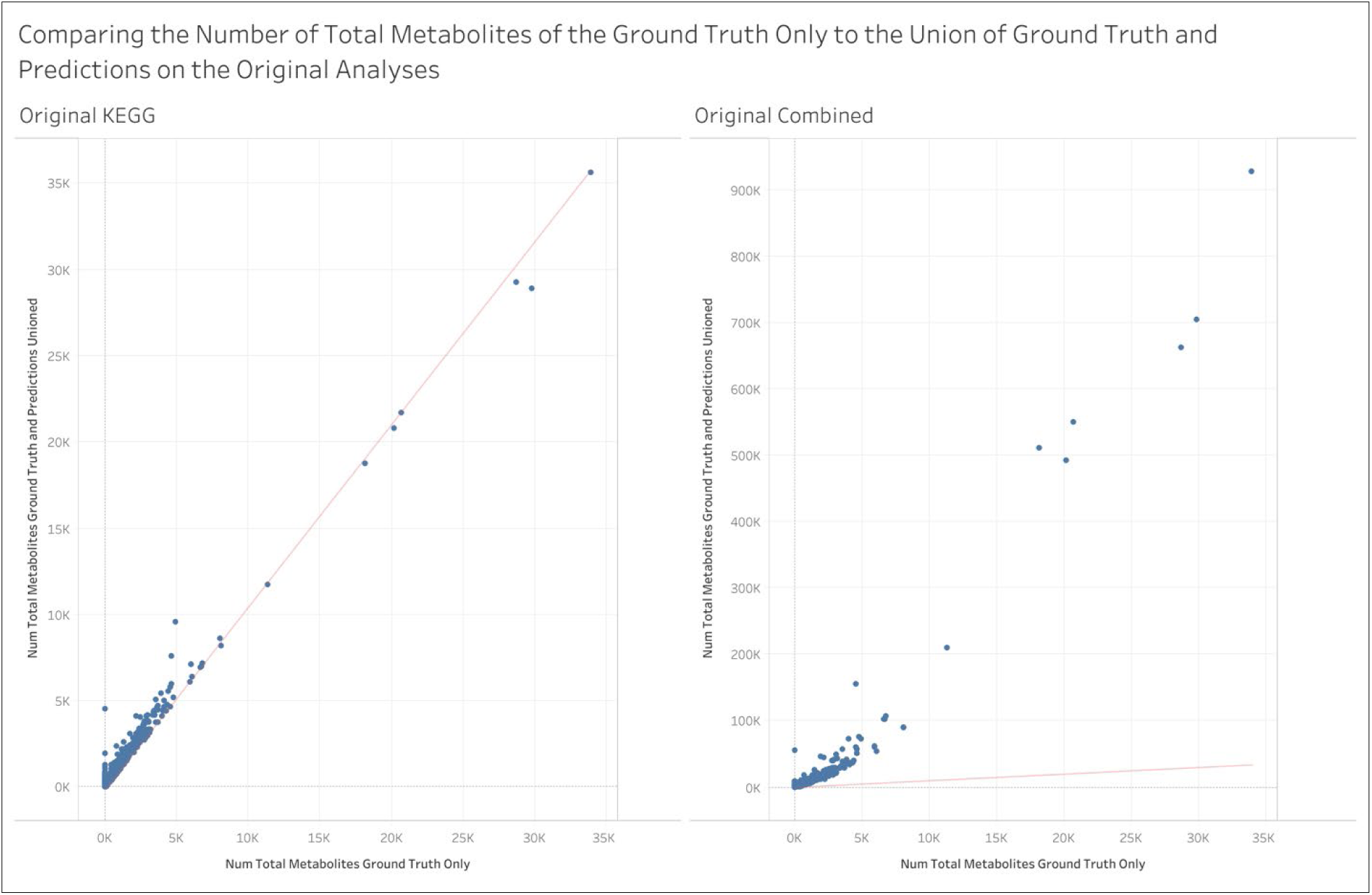
– Same as Figure S9 but for individual pathway IDs.

### Post-GSEA

**Figure S14.**
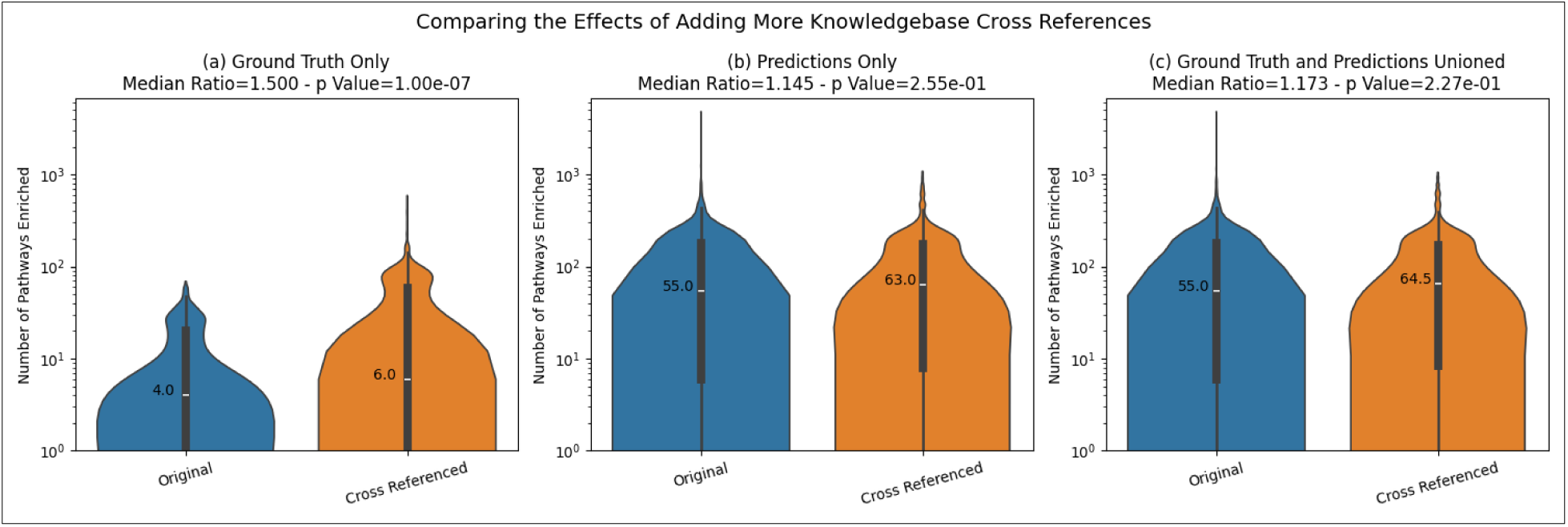
– The effect on the number of enriched pathways of adding metabolite ID cross references to the MW datasets for each type of annotations. Results are for individual pathway IDs.

**Figure S15.**
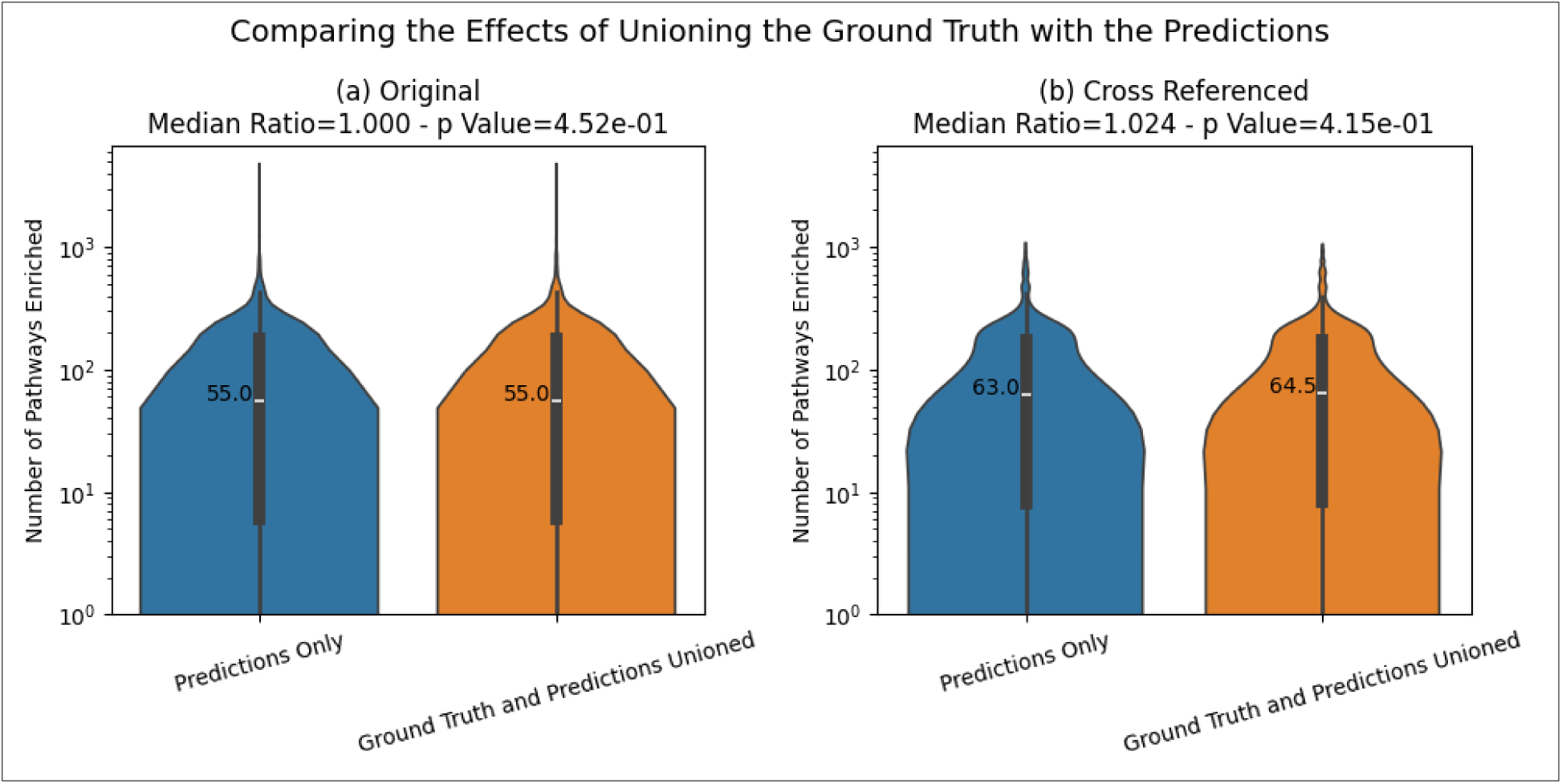
– The effect on the number of enriched pathways when unioning the predictions with the ground truth. Results are for individual pathway IDs.

**Figure S16.**
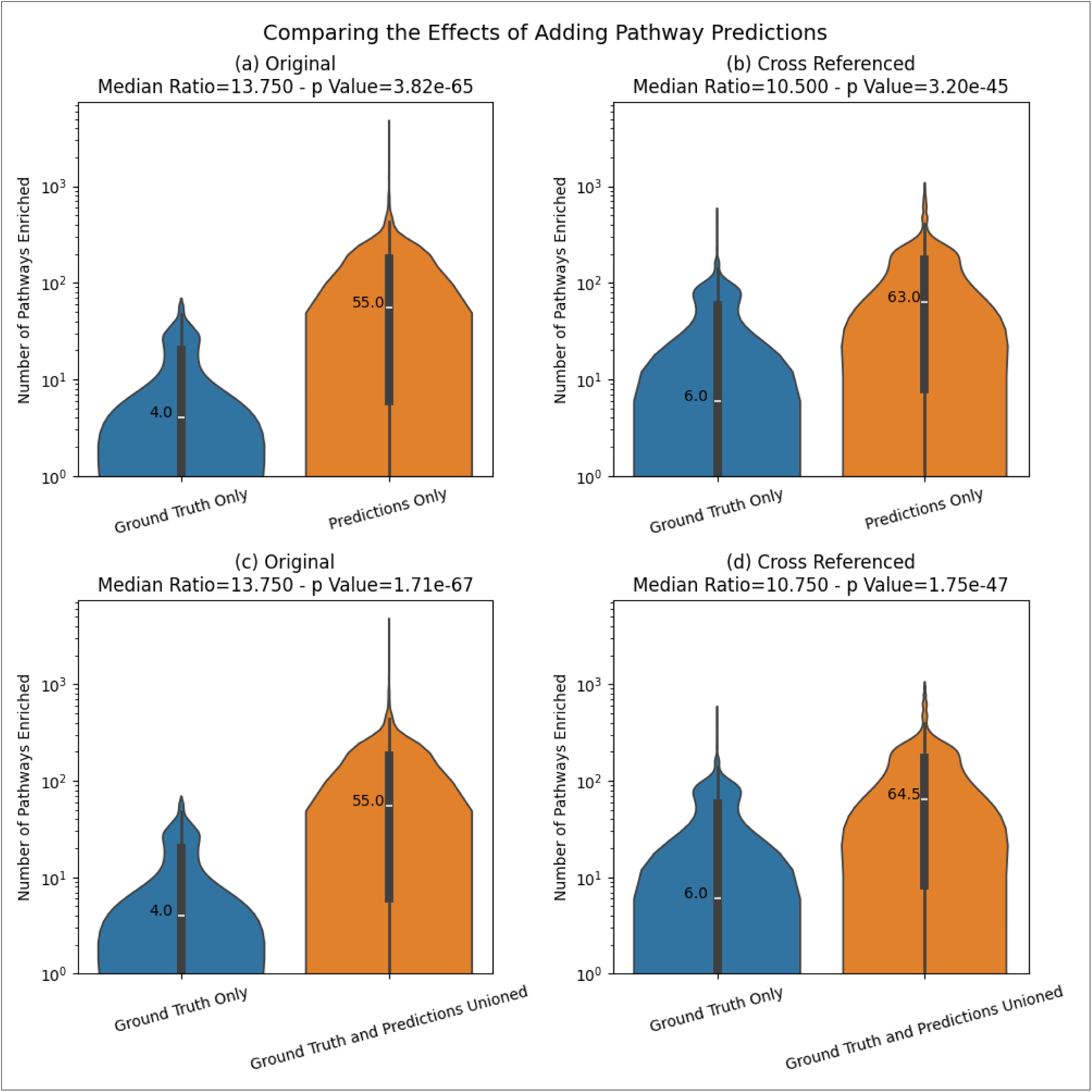
– The increase of the number of enriched pathways when adding predicted annotations. Results are for individual pathway IDs.

**Figure S17.**
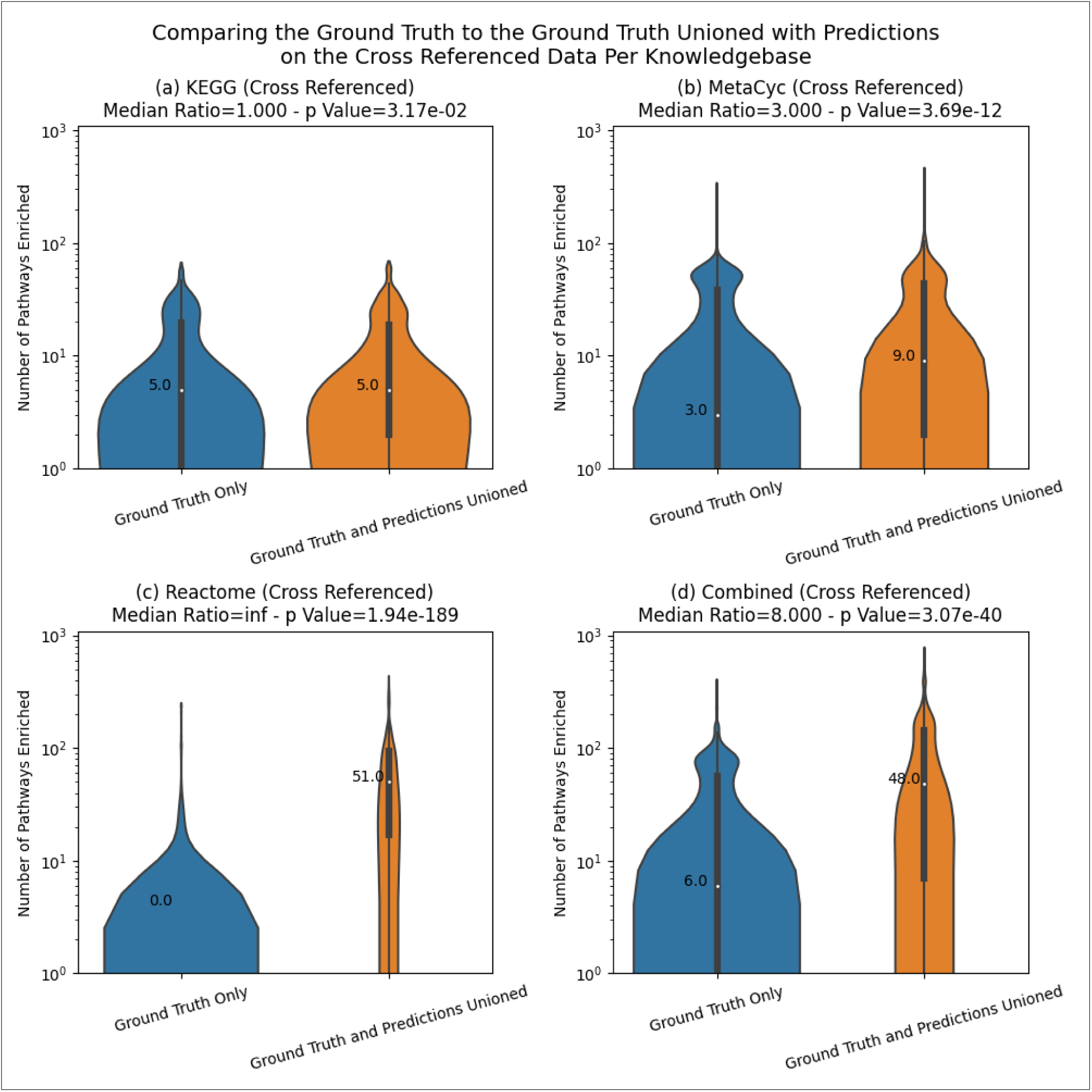
–Comparing the distribution across MW analyses of the number of enriched pathways between the ground truth pathway annotations and those unioned with the predicted annotations for the cross referenced MW datasets for each knowledgebase. Combined results included for comparison. Results are for unique pathway definitions.

**Figure S18.**
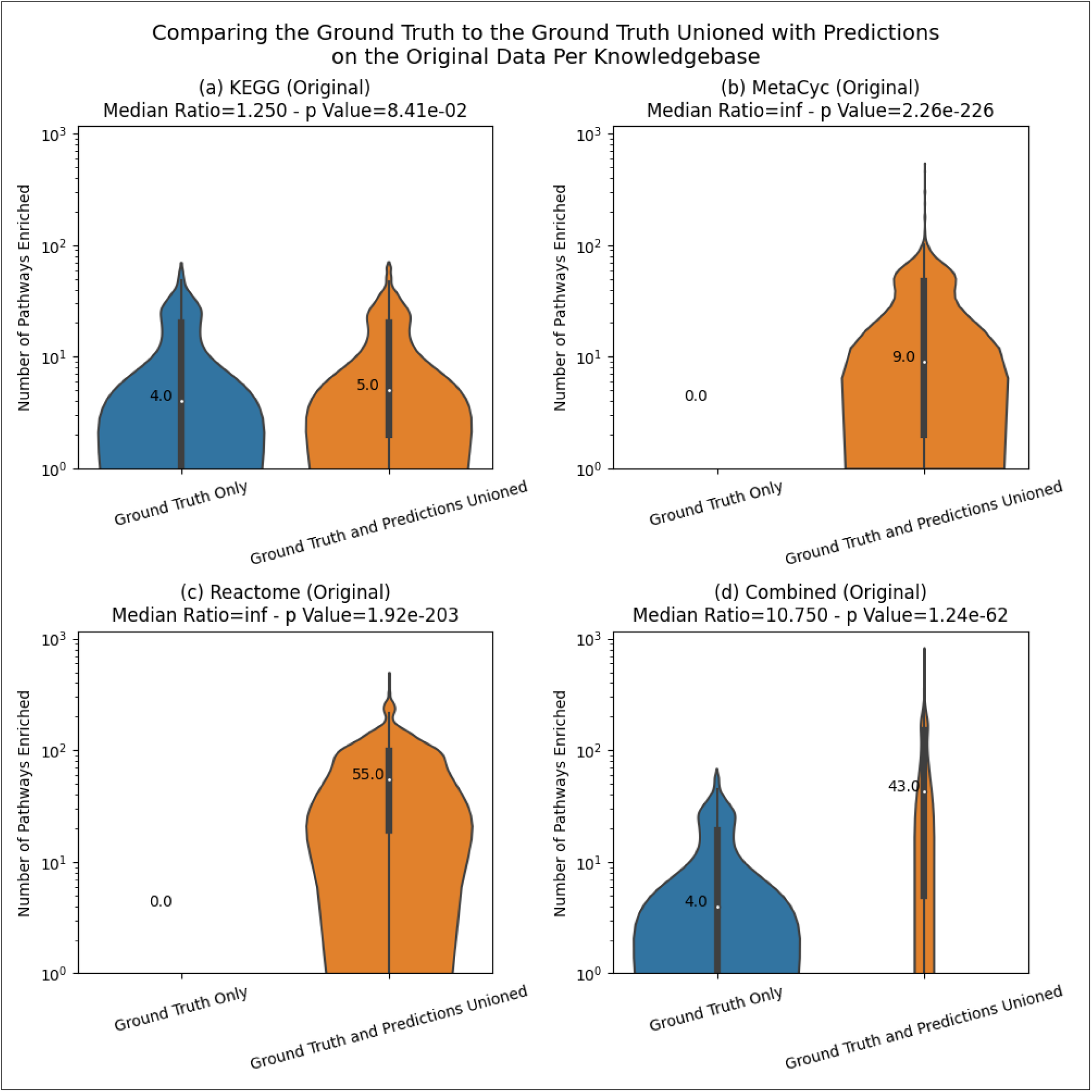
– Comparing the distribution across MW analyses of the number of enriched pathways between the ground truth pathway annotations and those unioned with the predicted annotations for the original MW datasets for each knowledgebase. Note that MetaCyc and Reactome had no ground truth annotations available since their metabolite IDs were not available in the original MW datasets. Combined results included for comparison. Results are for unique pathway definitions.

**Figure S19.**
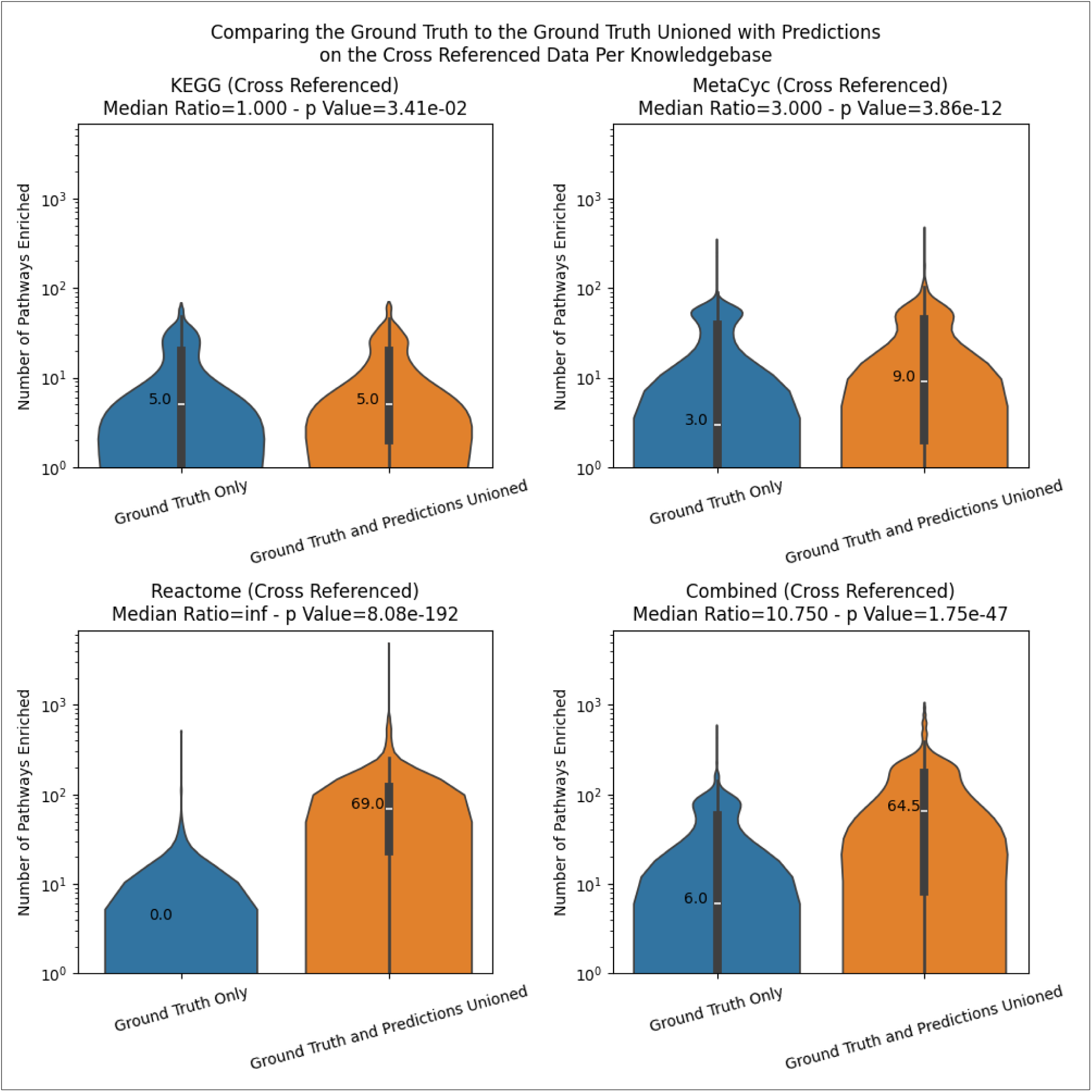
– Same as Figure S17 but for individual pathway IDs.

**Figure S20.**
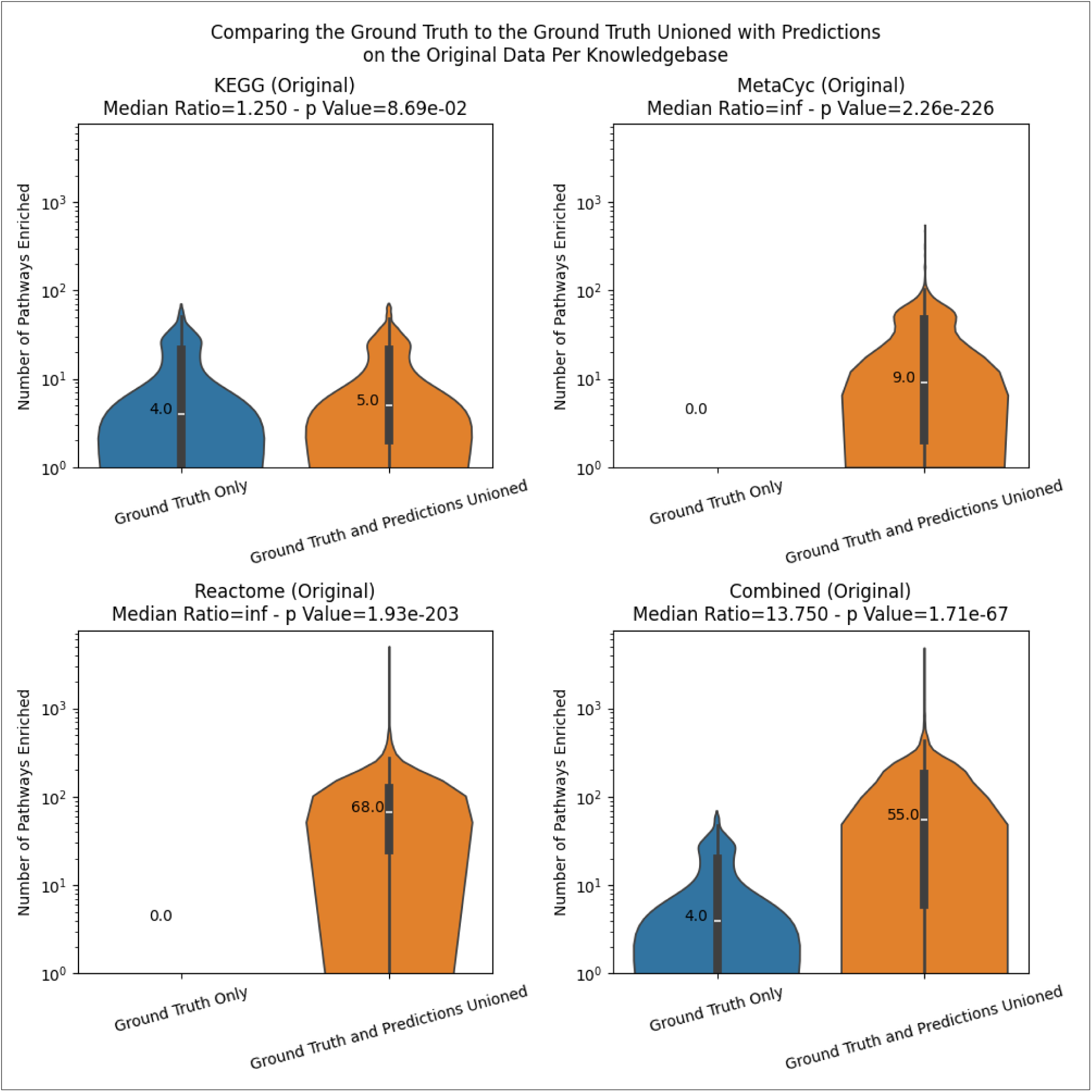
– Same as Figure S18 but for individual pathways.

### Information Loss and Gain

**Figure S21.**
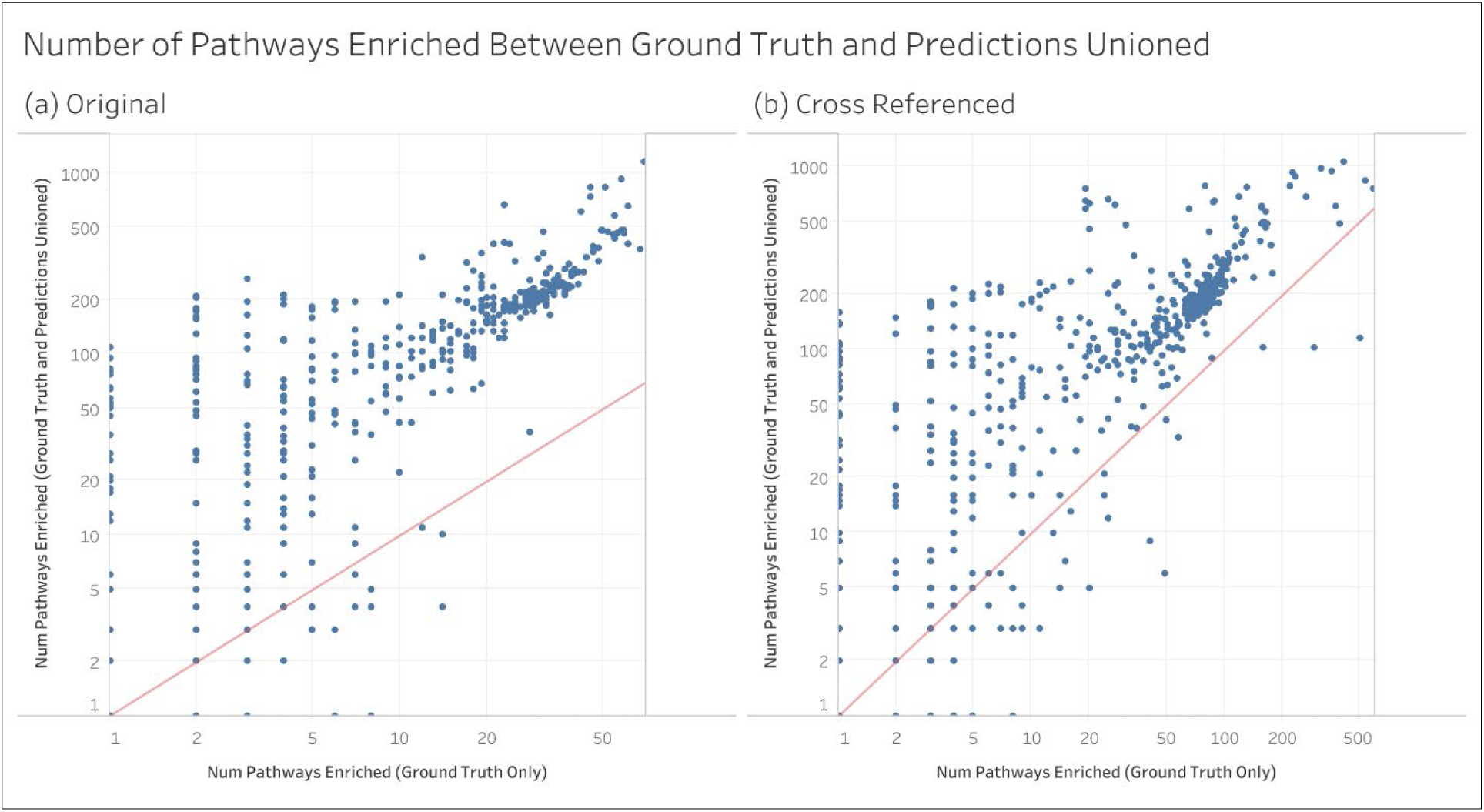
– The per dataset change in the number of enriched pathways between the ground truth and the ground truth unioned with the predictions. Each point is a separate MW dataset. Results are for individual pathway IDs.

**Figure S22.**
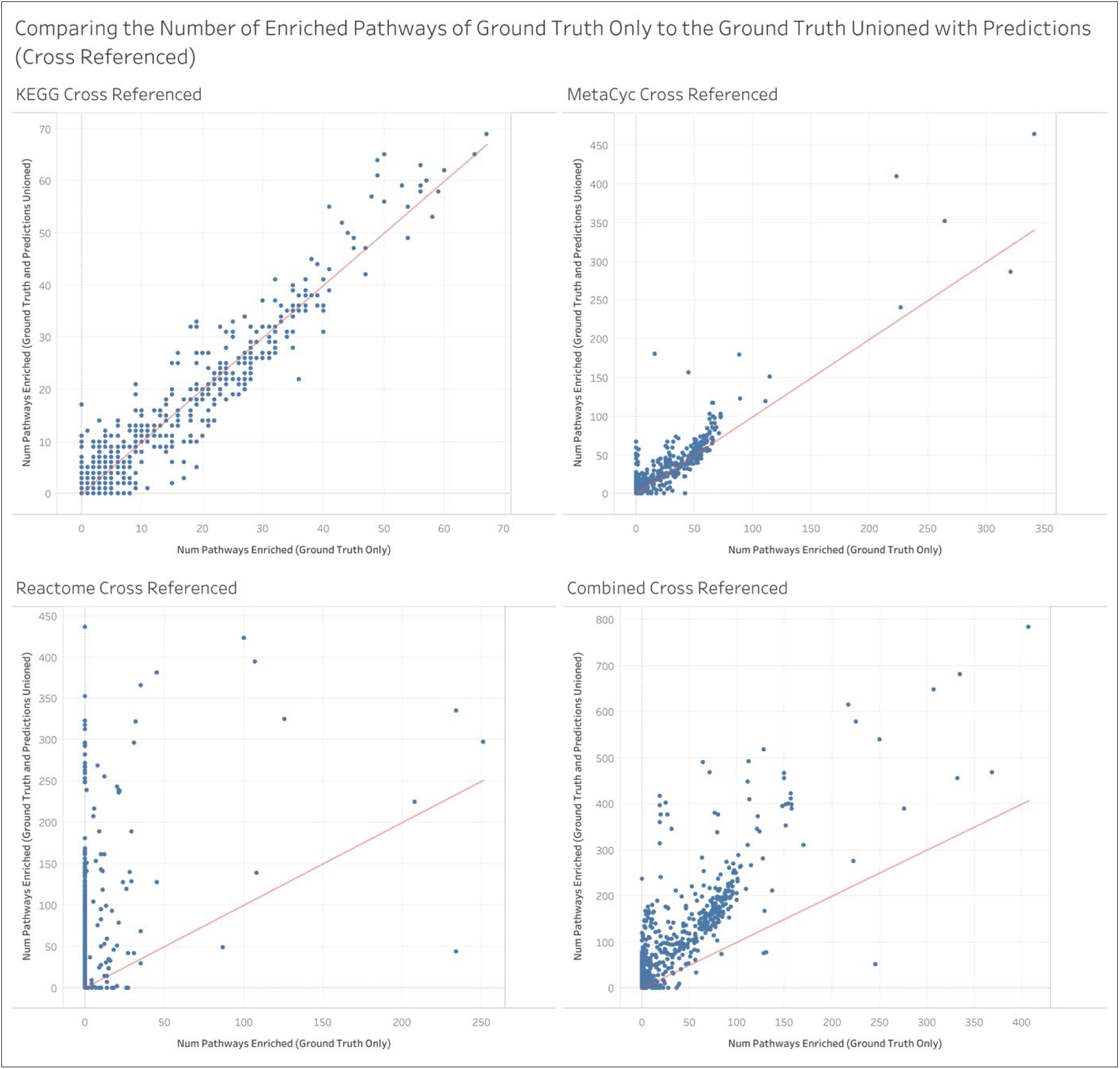
– Per analysis view of the change in the number of enriched pathways between the ground truth pathway annotations and those unioned with the predicted annotations for each knowledgebase on the cross-referenced MW datasets. Results are for unique pathway definitions.

**Figure S23.**
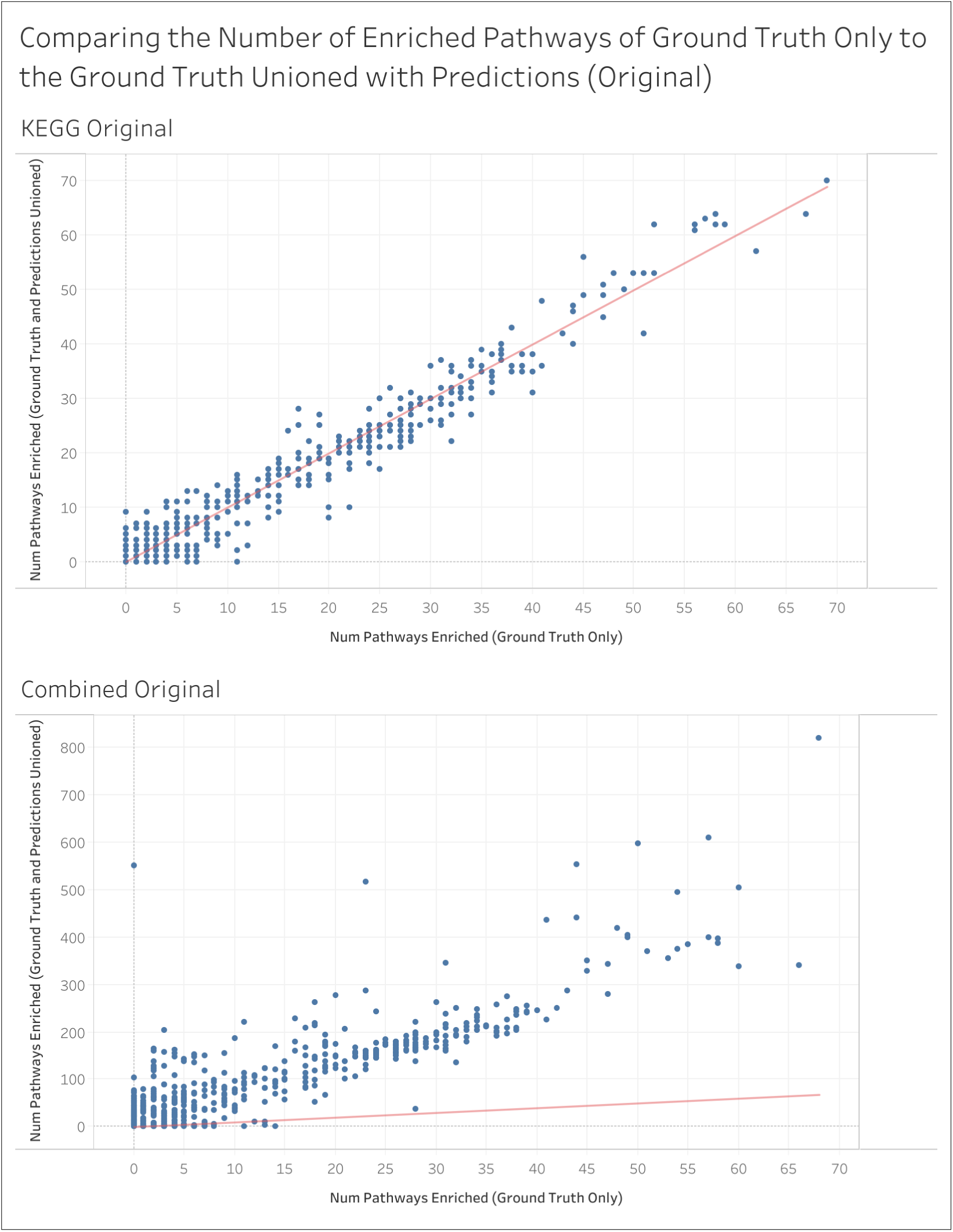
– Per analysis view of the change in the number of enriched pathways between the ground truth pathway annotations and those unioned with the predicted annotations for each knowledgebase on the original MW datasets. Note that MetaCyc and Reactome are not shown because these had no metabolite IDs available in the original datasets.

### Results are for unique pathway definitions

**Figure S24.**
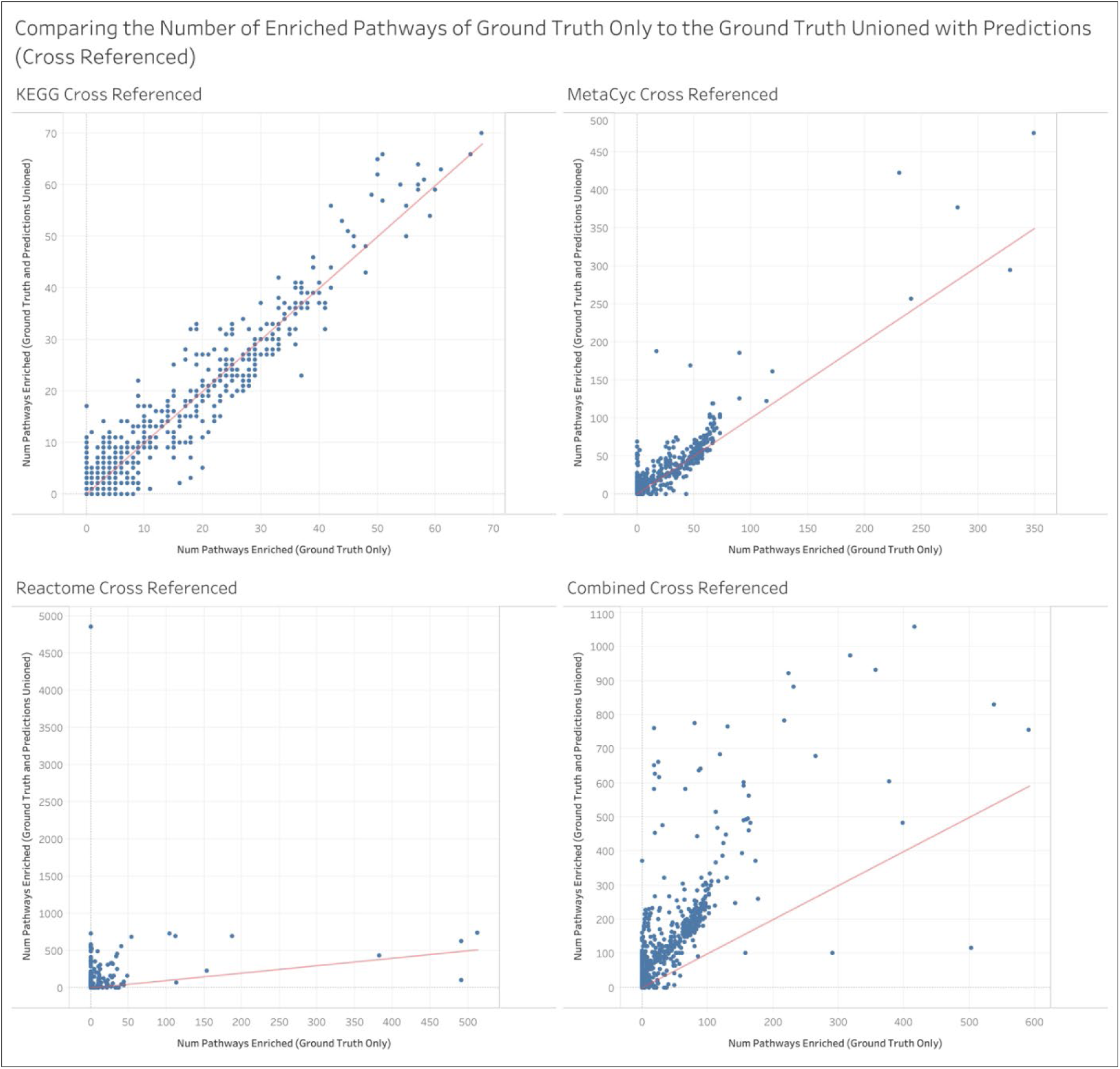
– Same as Figure S22 but for individual pathway IDs.

**Figure S25.**
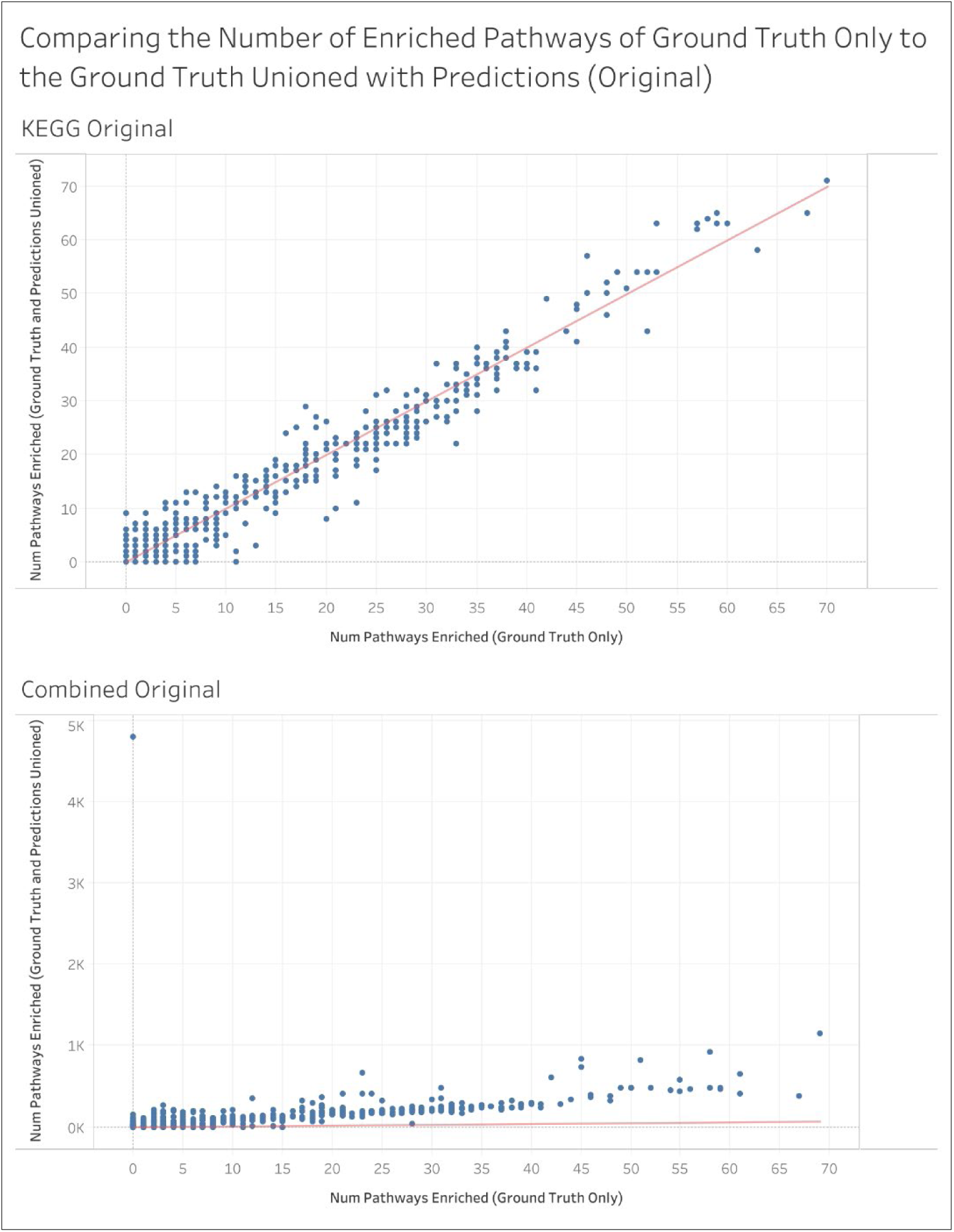
– Same as Figure S23 but for individual pathway IDs.

**Table S2.** –The amount of information gain and information loss summed across all enriched pathways and MW datasets. Results are for individual pathway IDs.

| Analysis Type | Annotation Type | Num Ground Truth Detected Only | Num Predictions Detected Only | Num Both Detected | Total Detected | Num Lost % | Num Gained % | Unchanged % |
| --- | --- | --- | --- | --- | --- | --- | --- | --- |
| Cross Referenced | Ground Truth and Predictions Unioned | 4,137 | 65,032 | 22,084 | 91,253 | 4.5 | 71.3 | 24.2 |
|  | Predictions Only | 4,080 | 69,234 | 22,141 | 95,455 | 4.3 | 72.5 | 23.2 |
| Original | Ground Truth and Predictions Unioned | 1,040 | 69,082 | 6,709 | 76,831 | 1.4 | 89.9 | 8.7 |
|  | Predictions Only | 1,030 | 69,204 | 6,719 | 76,953 | 1.3 | 89.9 | 8.7 |

**Figure S26.**
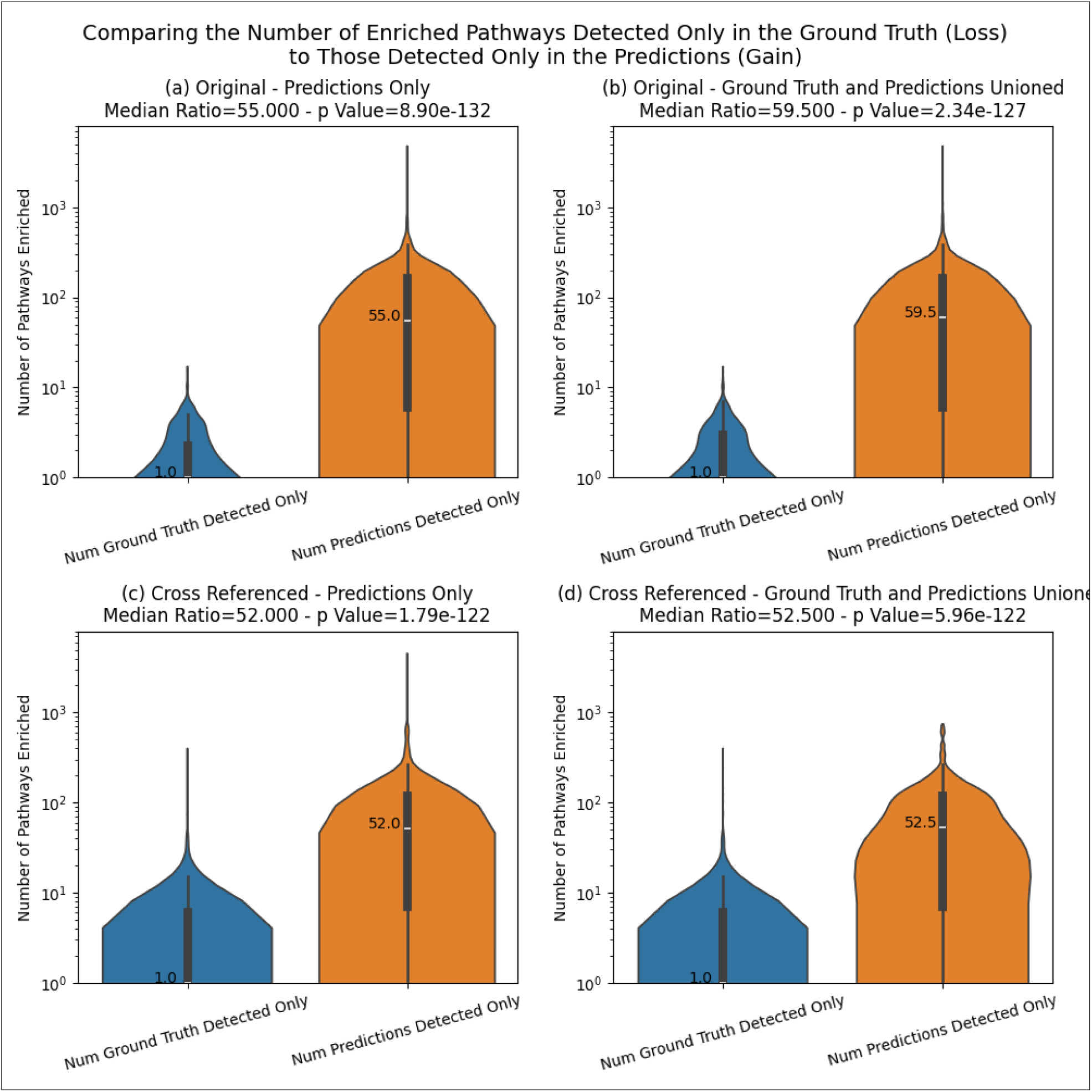
– Comparing the distribution of information loss to that of information gain across MW datasets. Results are for individual pathway IDs.

**Figure S27.**
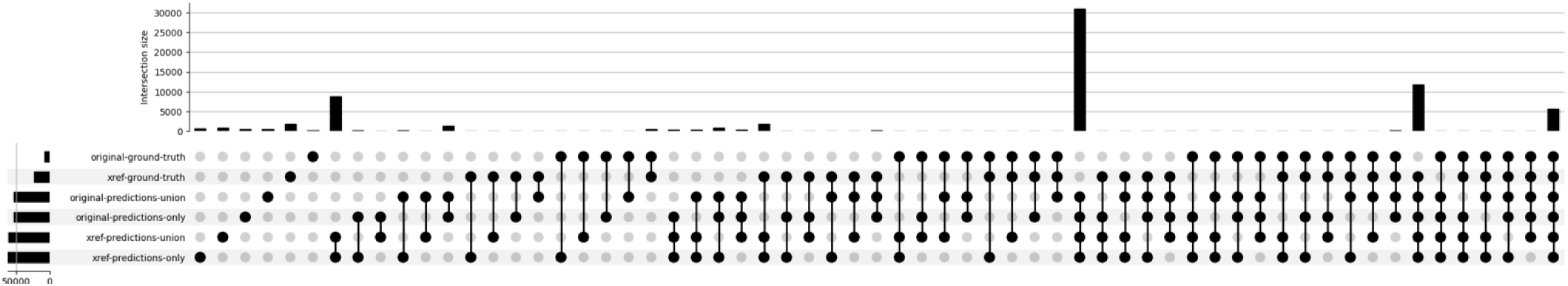
– Represents the information loss and information gain of enriched pathways between ground truth and predictions by showing the overlap of detected pathways. Results are for unique pathway definitions.

**Figure S28.**
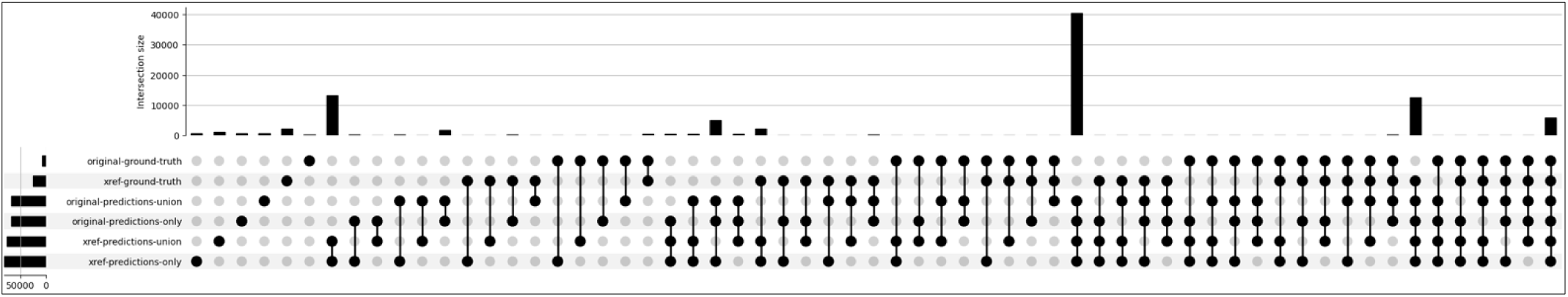
– Same as Figure S27 but for individual pathway IDs.

**Figure S29.**
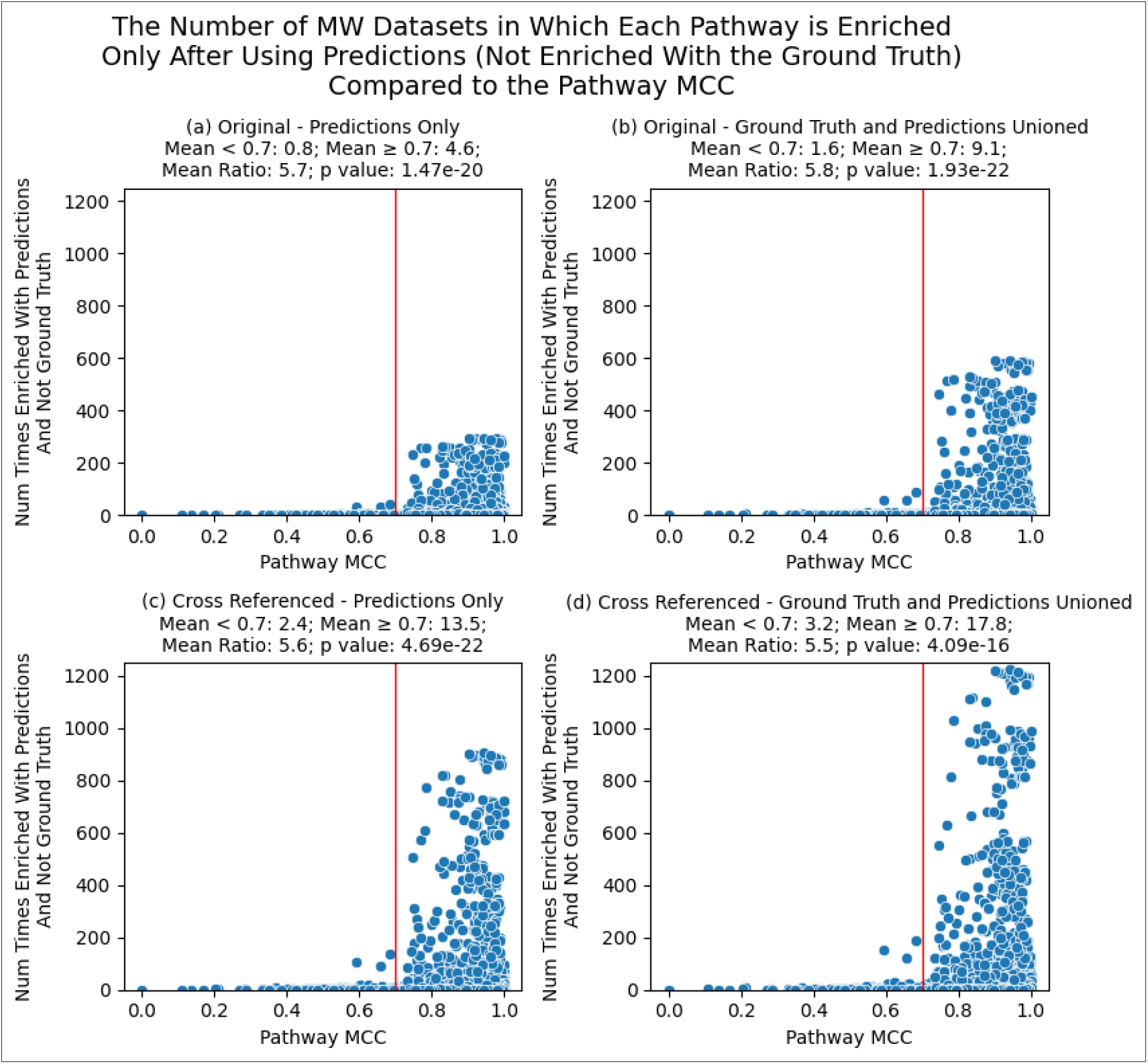
– Comparing the pathway MCC to the number of MW datasets wherein each pathway was not enriched when using ground truth annotations but was enriched when using predicted annotations. The red line is drawn at overall MCC = 0.7. Each point is a different pathway enriched in at least one dataset. Results are for individual pathway IDs.

**Table S3.** – The number of unique metabolites as part of metabolite-pathway annotations across MW datasets.

| Analysis Type | Annotation Type | Mean | Median | Max | #Datasets |
| --- | --- | --- | --- | --- | --- |
| Original | Ground Truth | 1919.1 | 0.0 | 6501 | 990 |
| Original | Predictions Unioned | 5102.1 | 0.0 | 18,411 | 990 |
| Cross Referenced | Ground Truth | 4905.4 | 0.0 | 18,141 | 990 |
| Cross Referenced | Predictions Unioned | 5241.3 | 228.0 | 18,424 | 990 |

**Table S4.** – Sum of 1 minus average Jaccard distance between enriched pathway definitions and every other pathway definition.

| Analysis Type | Annotation Type | Mean | Median | Max | #Datasets |
| --- | --- | --- | --- | --- | --- |
| Original | Ground Truth | 6.8 | 0.0 | 64.3 | 990 |
| Original | Predictions Unioned | 63.4 | 0.0 | 2574.5 | 990 |
| Cross Referenced | Ground Truth | 23.7 | 0.0 | 530.9 | 990 |
| Cross Referenced | Predictions Unioned | 73.6 | 1.0 | 1002.9 | 990 |

**Table 5.** – Hyperparameters used for the ML model.

| Activation Function | Hidden Layer Size | Num Layers | Dropout | Learning Rate | Beta 1 | Beta 2 | EPS | Weight Decay | Prediction Threshold |
| --- | --- | --- | --- | --- | --- | --- | --- | --- | --- |
| ReLU6 | 1229 | 3 | 0.063739 | 0.000719 | 0.565180 | 0.497934 | 0.000817 | 2.698780e-06 | 0.678975 |

## References

1. Voet, D.; Voet, J.G.; Pratt, C.W. Fundamentals of Biochemistry: Life at the Molecular; 5th ed.; Wiley, 2016;

2. Berg, J.M.; Tymoczko, J.L.; Gatto, G.J.; Stryer, L. Biochemistry; 9th ed.; W. H. Freeman: New York, NY, USA, 2019;

3. Nelson, D.L.; Cox, M.M. principles of biochemistry; 8th ed.; W. H. Freeman: New York, NY, USA, 2021;

4. Kim, S.J.; Kim, S.H.; Kim, J.H.; Hwang, S.; Yoo, H.J. Understanding metabolomics in biomedical research. Endocrinol Metab (Seoul) 2016, 31, 7–16, doi:10.3803/EnM.2016.31.1.7.

5. Gonzalez-Covarrubias, V.; Martínez-Martínez, E.; Del Bosque-Plata, L. The potential of metabolomics in biomedical applications. Metabolites 2022, 12, doi:10.3390/metabo12020194.

6. Kankainen, M.; Gopalacharyulu, P.; Holm, L.; Oresic, M. MPEA--metabolite pathway enrichment analysis. Bioinformatics 2011, 27, 1878–1879, doi:10.1093/bioinformatics/btr278.

7. Persicke, M.; Rückert, C.; Plassmeier, J.; Stutz, L.J.; Kessler, N.; Kalinowski, J.; Goesmann, A.; Neuweger, H. MSEA: metabolite set enrichment analysis in the MeltDB metabolomics software platform: metabolic profiling of Corynebacterium glutamicum as an example. Metabolomics 2012, 8, 310–322, doi:10.1007/s11306-011-0311-6.

8. Marco-Ramell, A.; Palau-Rodriguez, M.; Alay, A.; Tulipani, S.; Urpi-Sarda, M.; Sanchez-Pla, A.; Andres-Lacueva, C. Evaluation and comparison of bioinformatic tools for the enrichment analysis of metabolomics data. BMC Bioinformatics 2018, 19, 1, doi:10.1186/s12859-017-2006-0.

9. Acevedo, A.; Duran, C.; Ciucci, S.; Gerl, M.; Cannistraci, C.V. LIPEA: lipid pathway enrichment analysis. BioRxiv 2018, doi:10.1101/274969.

10. Zhang, B.; Hu, S.; Baskin, E.; Patt, A.; Siddiqui, J.K.; Mathé, E.A. Ramp: A comprehensive relational database of metabolomics pathways for pathway enrichment analysis of genes and metabolites. Metabolites 2018, 8, doi:10.3390/metabo8010016.

11. Lu, Y.; Pang, Z.; Xia, J. Comprehensive investigation of pathway enrichment methods for functional interpretation of LC-MS global metabolomics data. Brief. Bioinformatics 2023, 24, bbac553, doi:10.1093/bib/bbac553.

12. Barupal, D.K.; Fan, S.; Fiehn, O. Integrating bioinformatics approaches for a comprehensive interpretation of metabolomics datasets. Curr. Opin. Biotechnol. 2018, 54, 1–9, doi:10.1016/j.copbio.2018.01.010.

13. Kaever, A.; Landesfeind, M.; Feussner, K.; Mosblech, A.; Heilmann, I.; Morgenstern, B.; Feussner, I.; Meinicke, P. MarVis-Pathway: integrative and exploratory pathway analysis of non-targeted metabolomics data. Metabolomics 2015, 11, 764–777, doi:10.1007/s11306-014-0734-y.

14. Chagoyen, M.; Pazos, F. MBRole: enrichment analysis of metabolomic data. Bioinformatics 2011, 27, 730–731, doi:10.1093/bioinformatics/btr001.

15. Chen, Y.; Li, E.-M.; Xu, L.-Y. Guide to metabolomics analysis: A bioinformatics workflow. Metabolites 2022, 12, doi:10.3390/metabo12040357.

16. Kamburov, A.; Cavill, R.; Ebbels, T.M.D.; Herwig, R.; Keun, H.C. Integrated pathway-level analysis of transcriptomics and metabolomics data with IMPaLA. Bioinformatics 2011, 27, 2917–2918, doi:10.1093/bioinformatics/btr499.

17. Xia, J.; Wishart, D.S. MetPA: a web-based metabolomics tool for pathway analysis and visualization. Bioinformatics 2010, 26, 2342–2344, doi:10.1093/bioinformatics/btq418.

18. Kanehisa, M.; Goto, S. KEGG: Kyoto encyclopedia of genes and genomes. Nucleic Acids Res. 2000, 28, 27–30, doi:10.1093/nar/28.1.27.

19. Kanehisa, M. Toward understanding the origin and evolution of cellular organisms. Protein Sci. 2019, 28, 1947–1951, doi:10.1002/pro.3715.

20. Kanehisa, M.; Furumichi, M.; Sato, Y.; Kawashima, M.; Ishiguro-Watanabe, M. KEGG for taxonomy-based analysis of pathways and genomes. Nucleic Acids Res. 2023, 51, D587–D592, doi:10.1093/nar/gkac963.

21. Caspi, R.; Billington, R.; Keseler, I.M.; Kothari, A.; Krummenacker, M.; Midford, P.E.; Ong, W.K.; Paley, S.; Subhraveti, P.; Karp, P.D. The MetaCyc database of metabolic pathways and enzymesa 2019 update. Nucleic Acids Res. 2020, 48, D445–D453, doi:10.1093/nar/gkz862.

22. Milacic, M.; Beavers, D.; Conley, P.; Gong, C.; Gillespie, M.; Griss, J.; Haw, R.; Jassal, B.; Matthews, L.; May, B.; Petryszak, R.; Ragueneau, E.; Rothfels, K.; Sevilla, C.; Shamovsky, V.; Stephan, R.; Tiwari, K.; Varusai, T.; Weiser, J.; Wright, A.; Wu, G.; Stein, L.; Hermjakob, H.; D’Eustachio, P. The reactome pathway knowledgebase 2024. Nucleic Acids Res. 2024, 52, D672–D678, doi:10.1093/nar/gkad1025.

23. Wieder, C.; Frainay, C.; Poupin, N.; Rodríguez-Mier, P.; Vinson, F.; Cooke, J.; Lai, R.P.; Bundy, J.G.; Jourdan, F.; Ebbels, T. Pathway analysis in metabolomics: Recommendations for the use of over-representation analysis. PLoS Comput. Biol. 2021, 17, e1009105, doi:10.1371/journal.pcbi.1009105.

24. Stobbe, M.D.; Houten, S.M.; Jansen, G.A.; van Kampen, A.H.C.; Moerland, P.D. Critical assessment of human metabolic pathway databases: a stepping stone for future integration. BMC Syst. Biol. 2011, 5, 165, doi:10.1186/1752-0509-5-165.

25. Huckvale, E.D.; Powell, C.D.; Jin, H.; Moseley, H.N.B. Benchmark dataset for training machine learning models to predict the pathway involvement of metabolites. Metabolites 2023, 13, 1120, doi:10.3390/metabo13111120.

26. Starke, C.; Wegner, A. Metamdb: metabolic atom mapping database. Metabolites 2022, 12, doi:10.3390/metabo12020122.

27. Heller, S.; McNaught, A.; Stein, S.; Tchekhovskoi, D.; Pletnev, I. InChI – the worldwide chemical structure identifier standard. J. Cheminform. 2013, 5, 7, doi:10.1186/1758-2946-5-7.

28. Heller, S.R.; McNaught, A.; Pletnev, I.; Stein, S.; Tchekhovskoi, D. Inchi, the IUPAC international chemical identifier. J. Cheminform. 2015, 7, 23, doi:10.1186/s13321-015-0068-4.

29. Goodman, J.M.; Pletnev, I.; Thiessen, P.; Bolton, E.; Heller, S.R. InChI version 1.06: now more than 99.99% reliable. J. Cheminform. 2021, 13, 40, doi:10.1186/s13321-021-00517-z.

30. Joachimiak, M.P.; Caufield, J.H.; Harris, N.L.; Kim, H.; Mungall, C.J. Gene set summarization using large language models. arXiv 2024, doi:10.48550/arxiv.2305.13338.

31. Hu, M.; Alkhairy, S.; Lee, I.; Pillich, R.T.; Fong, D.; Smith, K.; Bachelder, R.; Ideker, T.; Pratt, D. Evaluation of large language models for discovery of gene set function. Nat. Methods 2025, 22, 82–91, doi:10.1038/s41592-024-02525-x.

32. Huckvale, E.D.; Moseley, H.N.B. A cautionary tale about properly vetting datasets used in supervised learning predicting metabolic pathway involvement. PLoS ONE 2024, 19, e0299583, doi:10.1371/journal.pone.0299583.

33. Huckvale, E.D.; Moseley, H.N.B. Predicting the pathway involvement of metabolites based on combined metabolite and pathway features. Metabolites 2024, 14, 266, doi:10.3390/metabo14050266.

34. Huckvale, E.D.; Moseley, H.N.B. Predicting the Association of Metabolites with Both Pathway Categories and Individual Pathways. Metabolites 2024, 14, 510, doi:10.3390/metabo14090510.

35. Huckvale, E.D.; Moseley, H.N.B. Predicting the pathway involvement of all pathway and associated compound entries defined in the kyoto encyclopedia of genes and genomes. Metabolites 2024, 14, 582, doi:10.3390/metabo14110582.

36. Huckvale, E.D.; Moseley, H.N.B. Predicting the pathway involvement of compounds annotated in the reactome knowledgebase. Metabolites 2025, 15, 161, doi:10.3390/metabo15030161.

37. Unable to find information for 17159280.

38. Huckvale, E.D.; Moseley, H.N.B. Chemical representation standardization needed to generalize metabolic pathway involvement prediction across the Kyoto Encyclopedia of Genes and Genomes, Reactome, and MetaCyc knowledgebases. Metabolites 2026, 16, 357, doi:10.3390/metabo16060357.

39. Jin, H.; Mitchell, J.M.; Moseley, H.N.B. Atom Identifiers Generated by a Neighborhood-Specific Graph Coloring Method Enable Compound Harmonization across Metabolic Databases. Metabolites 2020, 10, 368, doi:10.3390/metabo10090368.

40. Jin, H.; Moseley, H.N.B. Hierarchical Harmonization of Atom-Resolved Metabolic Reactions across Metabolic Databases. Metabolites 2021, 11, 431, doi:10.3390/metabo11070431.

41. Jin, H.; Moseley, H.N.B. md_harmonize: A Python Package for Atom-Level Harmonization of Public Metabolic Databases. Metabolites 2023, 13, 1199, doi:10.3390/metabo13121199.

42. Zhang, S.; Tong, H.; Xu, J.; Maciejewski, R. Graph convolutional networks: a comprehensive review. Compu Social Networls 2019, 6, 11, doi:10.1186/s40649-019-0069-y.

43. Khemani, B.; Patil, S.; Kotecha, K.; Tanwar, S. A review of graph neural networks: concepts, architectures, techniques, challenges, datasets, applications, and future directions. J. Big Data 2024, 11, 18, doi:10.1186/s40537-023-00876-4.

44. Zhang, Y.; Yang, Q. An overview of multitask learning. Natl Sci Rev 2017, doi:10.1093/nsr/nwx105.

45. Zhang, Y.; Yang, Q. A Survey on Multi-Task Learning. IEEE Trans. Knowl. Data Eng. 2021, 1–1, doi:10.1109/TKDE.2021.3070203.

46. Caruana, R. Multitask Learning. Springer Science and Business Media LLC 1997, 28, 41–75, doi:10.1023/a:1007379606734.

47. Bengio, S.; Dembczynski, K.; Joachims, T.; Kloft, M.; Varma, M. Extreme Classification (Dagstuhl Seminar 18291). Schloss Dagstuhl – Leibniz-Zentrum für Informatik 2019, doi:10.4230/dagrep.8.7.62.

48. Varma, M. Extreme classification. Commun. ACM 2019, 62, 44–45, doi:10.1145/3355628.

49. Weiss, K.; Khoshgoftaar, T.M.; Wang, D. A survey of transfer learning. J. Big Data 2016, 3, 9, doi:10.1186/s40537-016-0043-6.

50. Zhuang, F.; Qi, Z.; Duan, K.; Xi, D.; Zhu, Y.; Zhu, H.; Xiong, H.; He, Q. A comprehensive survey on transfer learning. Proc. IEEE 2020, 109, 43–76, doi:10.1109/JPROC.2020.3004555.

51. Iman, M.; Arabnia, H.R.; Rasheed, K. A review of deep transfer learning and recent advancements. Technologies 2023, 11, 40, doi:10.3390/technologies11020040.

52. Chicco, D.; Jurman, G. The advantages of the Matthews correlation coefficient (MCC) over F1 score and accuracy in binary classification evaluation. BMC Genomics 2020, 21, 6, doi:10.1186/s12864-019-6413-7.

53. Sud, M.; Fahy, E.; Cotter, D.; Azam, K.; Vadivelu, I.; Burant, C.; Edison, A.; Fiehn, O.; Higashi, R.; Nair, K.S.; Sumner, S.; Subramaniam, S. Metabolomics Workbench: An international repository for metabolomics data and metadata, metabolite standards, protocols, tutorials and training, and analysis tools. Nucleic Acids Res. 2016, 44, D463–70, doi:10.1093/nar/gkv1042.

54. Smelter, A.; Moseley, H.N.B. A Python library for FAIRer access and deposition to the Metabolomics Workbench Data Repository. Metabolomics 2018, 14, 64, doi:10.1007/s11306-018-1356-6.

55. Powell, C.D.; Moseley, H.N.B. The mwtab Python Library for RESTful Access and Enhanced Quality Control, Deposition, and Curation of the Metabolomics Workbench Data Repository. Metabolites 2021, 11, doi:10.3390/metabo11030163.

56. Powell, C.D.; Moseley, H.N.B. The metabolomics workbench file status website: a metadata repository promoting FAIR principles of metabolomics data. BMC Bioinformatics 2023, 24, 299, doi:10.1186/s12859-023-05423-9.

57. Kim, S.; Thiessen, P.A.; Bolton, E.E.; Chen, J.; Fu, G.; Gindulyte, A.; Han, L.; He, J.; He, S.; Shoemaker, B.A.; Wang, J.; Yu, B.; Zhang, J.; Bryant, S.H. PubChem Substance and Compound databases. Nucleic Acids Res. 2016, 44, D1202–13, doi:10.1093/nar/gkv951.

58. Hähnke, V.D.; Kim, S.; Bolton, E.E. PubChem chemical structure standardization. J. Cheminform. 2018, 10, 36, doi:10.1186/s13321-018-0293-8.

59. Dalby, A.; Nourse, J.G.; Hounshell, W.D.; Gushurst, A.K.I.; Grier, D.L.; Leland, B.A.; Laufer, J. Description of several chemical structure file formats used by computer programs developed at Molecular Design Limited. J. Chem. Inf. Model. 1992, 32, 244–255, doi:10.1021/ci00007a012.

60. Hastings, J.; Owen, G.; Dekker, A.; Ennis, M.; Kale, N.; Muthukrishnan, V.; Turner, S.; Swainston, N.; Mendes, P.; Steinbeck, C. ChEBI in 2016: Improved services and an expanding collection of metabolites. Nucleic Acids Res. 2016, 44, D1214–9, doi:10.1093/nar/gkv1031.

61. Cohen, A. fuzzywuzzy; PyPi, 2020;

62. Flight, R.M.; Harrison, B.J.; Mohammad, F.; Bunge, M.B.; Moon, L.D.F.; Petruska, J.C.; Rouchka, E.C. categoryCompare, an analytical tool based on feature annotations. Front. Genet. 2014, 5, 98, doi:10.3389/fgene.2014.00098.

63. Flight, R.M. categoryCompare2; Moseley Bioinformatics Lab, 2025;

64. Subramanian, A.; Tamayo, P.; Mootha, V.K.; Mukherjee, S.; Ebert, B.L.; Gillette, M.A.; Paulovich, A.; Pomeroy, S.L.; Golub, T.R.; Lander, E.S.; Mesirov, J.P. Gene set enrichment analysis: a knowledge-based approach for interpreting genome-wide expression profiles. Proc Natl Acad Sci USA 2005, 102, 15545–15550, doi:10.1073/pnas.0506580102.

65. Korotkevich, G.; Sukhov, V.; Budin, N.; Shpak, B.; Artyomov, M.N.; Sergushichev, A. Fast gene set enrichment analysis. BioRxiv 2016, doi:10.1101/060012.

66. Maćkiewicz, A.; Ratajczak, W. Principal components analysis (PCA). Comput. Geosci. 1993, 19, 303–342, doi:10.1016/0098-3004(93)90090-R.

67. Benjamini, Y.; Hochberg, Y. Controlling the false discovery rate: a practical and powerful approach to multiple testing. Journal of the Royal Statistical Society: Series B (Methodological) 1995, 57, 289–300, doi:10.1111/j.2517-6161.1995.tb02031.x.

68. McKnight, P.E.; Najab, J. Mann-Whitney U Test. In The corsini encyclopedia of psychology; Weiner, I. B., Craighead, W. E., Eds.; John Wiley & Sons, Inc.: Hoboken, NJ, USA, 2010 ISBN 9780470479216.

69. Team, R.D.C. R: A language and environment for statistical computing. *R Found*. Stat. Comput. 2016.

70. Rossum, G.V.; Drake, F.L. Python 3 Reference Manual; CreateSpace, 2009; ISBN 1441412697.

71. The pandas development team pandas-dev/pandas: Pandas 1.0.3. Zenodo 2020, doi:10.5281/zenodo.3509134.

72. Harris, C.R.; Millman, K.J.; van der Walt, S.J.; Gommers, R.; Virtanen, P.; Cournapeau, D.; Wieser, E.; Taylor, J.; Berg, S.; Smith, N.J.; Kern, R.; Picus, M.; Hoyer, S.; van Kerkwijk, M.H.; Brett, M.; Haldane, A.; Del Río, J.F.; Wiebe, M.; Peterson, P.; Gérard-Marchant, P.; Sheppard, K.; Reddy, T.; Weckesser, W.; Abbasi, H.; Gohlke, C.; Oliphant, T.E. Array programming with NumPy. Nature 2020, 585, 357–362, doi:10.1038/s41586-020-2649-2.

73. Pedregosa, F.; Varoquaux, G.; Gramfort, A.; Michel, V.; Thirion, B.; Grisel, O.; Blondel, M.; Müller, A.; Nothman, J.; Louppe, G.; Prettenhofer, P.; Weiss, R.; Dubourg, V.; Vanderplas, J.; Passos, A.; Cournapeau, D.; Brucher, M.; Perrot, M.; Duchesnay, É. Scikit-learn: Machine Learning in Python. arXiv 2012, doi:10.48550/arxiv.1201.0490.

74. Virtanen, P.; Gommers, R.; Oliphant, T.E.; Haberland, M.; Reddy, T.; Cournapeau, D.; Burovski, E.; Peterson, P.; Weckesser, W.; Bright, J.; van der Walt, S.J.; Brett, M.; Wilson, J.; Millman, K.J.; Mayorov, N.; Nelson, A.R.J.; Jones, E.; Kern, R.; Larson, E.; Carey, C.J.; Polat, İ.; Feng, Y.; Moore, E.W.; VanderPlas, J.; Laxalde, D.; Perktold, J.; Cimrman, R.; Henriksen, I.; Quintero, E.A.; Harris, C.R.; Archibald, A.M.; Ribeiro, A.H.; Pedregosa, F.; van Mulbregt, P.; SciPy 1.0 Contributors SciPy 1.0: fundamental algorithms for scientific computing in Python. Nat. Methods 2020, 17, 261–272, doi:10.1038/s41592-019-0686-2.

75. Salesforce Tableau Public; Salesforce, 2024;

76. Waskom, M. seaborn: statistical data visualization. JOSS 2021, 6, 3021, doi:10.21105/joss.03021.

77. Hunter, J.D. Matplotlib: A 2D Graphics Environment. Comput. Sci. Eng. 2007, 9, 90–95, doi:10.1109/MCSE.2007.55.

78. Kluyver, T.; Ragan-Kelley, B.; Pérez, F.; Granger, B.; Bussonnier, M.; Frederic, J.; Kelley, K.; Hamrick, J.; Grout, J.; Corlay, S.; Ivanov, P.; Avila, D.; Abdalla, S.; Willing, C.; Team, J. development Jupyter Notebooks – a publishing format for reproducible computational workflows. In Positioning and Power in Academic Publishing: Players, Agents and Agendas; Loizides, F., Scmidt, B., Eds.; IOS Press: Netherlands, 2016; pp. 87–90.

